# Neanderthal introgressed ancestry reveals human genomic regions enriched with recessive deleterious mutations

**DOI:** 10.1101/2025.05.07.652751

**Authors:** Xinjun Zhang, Lingxuan Zhu, Jiongxuan Yang, Nina Sachdev, Jazlyn Mooney, Harold Z. Wang, Sriram Sankararaman, Kirk E. Lohmueller

## Abstract

Negative selection on deleterious mutations plays a key role in shaping human genetic variation, yet the dominance effects of these mutations remain poorly understood because existing statistical methods often cannot distinguish dominance effects from the overall selective effects. In this work, we take a fundamentally different approach by leveraging the distribution of Neanderthal ancestry across the human genome. Simulations show that recessive deleterious mutations can increase archaic introgressed ancestry through heterosis in the absence of positive selection, in contrast to the depletion expected under additive effects. We use this signal to develop *DominL*, a machine learning classifier trained on simulations of human demography with Neanderthal introgression to identify megabase-scale genomic windows enriched with recessive mutations. *DominL* demonstrates robust accuracy, with particularly high power in exon-dense regions. Applied to 7 non-African populations from the 1000 Genomes Project, *DominL* identifies approximately 3-9% of the genome as enriched for recessive mutations, with most high-confidence regions shared across populations. Predicted regions show patterns consistent with expected signatures from recessive mutations, including weakened background selection, depletion of runs of homozygosity, and enrichment of non-additive trait-associated variants. These regions also contain genes associated with metabolic and immune-related functions.

## Introduction

Dominance is one of the most fundamental concepts in genetics^1–4^ and has many implications for a myriad of phenomena in population genetics, including the magnitude of inbreeding depression, the rate and efficiency of natural selection, and the prevalence and evolution of disease^5–8^. Despite the inarguable importance of dominance, its distribution and biological effects have only been characterized in a small number of organisms, not including humans. Due to the poor understanding of dominance in humans, most population genetic studies impose an arbitrary assumption that dominance across the genome is a constant, being either additive, recessive, or a value in between. For example, many studies estimating the distribution of fitness effects or predicting polygenic disease risks assume strict additivity (*h*=0.5)^9,10^ of all genetic variants. This assumption, however, poorly fits the empirical data in most model organisms^11–13^, and can bias downstream inferences, resulting in unreliable conclusions with major consequences in biomedical applications^14,15^. Therefore, there is a pressing need for an accurate measure of dominance.

While historically there have been two main approaches to estimate dominance, both suffer from serious limitations. One approach is to estimate dominance through experimental methods^12,13,16–20^ such as gene knock-outs. However, this approach is limited by two factors. First, it is difficult to be scaled genome-wide. Second, and more importantly, experimental knockouts are inapplicable to most of the vertebrate species that are not manipulatable in laboratories^13^, including humans. To bypass the experimental constraints, another common approach to estimate dominance is to infer its distribution through the summaries of genomic variation data, such as the site frequency spectrum (SFS)^21–23^. While it is straightforward to tabulate a SFS on genetic variants found in whole genome sequence data from individuals in natural populations such as humans, the allele frequency-based approach suffers from an inability to disentangle the selection coefficient of a deleterious mutation (*s*) from its dominance coefficient (*h*). Studies on model organisms suggest that most strongly deleterious variants tend to be recessive^2,24^. Consequently, a negative relationship between the strength of negative selection and the dominance coefficient has been observed^23^, which is known as the *hs* relationship. It turns out that different combinations of *h* and *s* can produce highly similar patterns of empirical statistics such as SFS^25^, meaning that allele frequency patterns could only provide estimates of the *hs* compound rather than *h* by itself.

Therefore, the key to detect dominance is to identify a genomic feature that is sensitive to changes in dominance (*h*), but remains insensitive to variation in selection coefficients (*s*). The spatial pattern of introgressed archaic hominin ancestry in modern human genomes, such as the Neanderthal and Denisovan ancestry, is such a genomic feature. Previous studies showed that archaic introgressed ancestry in modern humans may have increased drastically in regions of the genome carrying many recessive deleterious mutations^26–28^, even in the absence of positive selection. Archaic ancestry increases in humans because by the time of archaic introgression (approx. 50,000 years ago), modern humans and archaic hominins had been separated from each other for approximately half a million years^29^, during which time they accumulated a substantial number of private genetic variants, including recessive deleterious variants. Upon introgression, the hybrids are expected to be mostly heterozygous at these private variants. If most deleterious private variants at a given genomic region are recessive, the deleterious fitness effects of these variants would be masked in the Neanderthal-human hybrid individuals, leading to an increase in Neanderthal ancestry at these loci through heterosis (Fig. 1a). In contrast, if most deleterious mutations have additive effects, negative selection would remove the Neanderthal introgressed ancestry, as shown in, empirical human genomes^5,30,31^ as well as simulations (Fig. 1b). Our earlier work^28^ further showed that this heterosis pattern is particularly pronounced in regions of the genome with high exon density, as recessive deleterious mutations accumulate faster in such regions.

**Figure 1:**
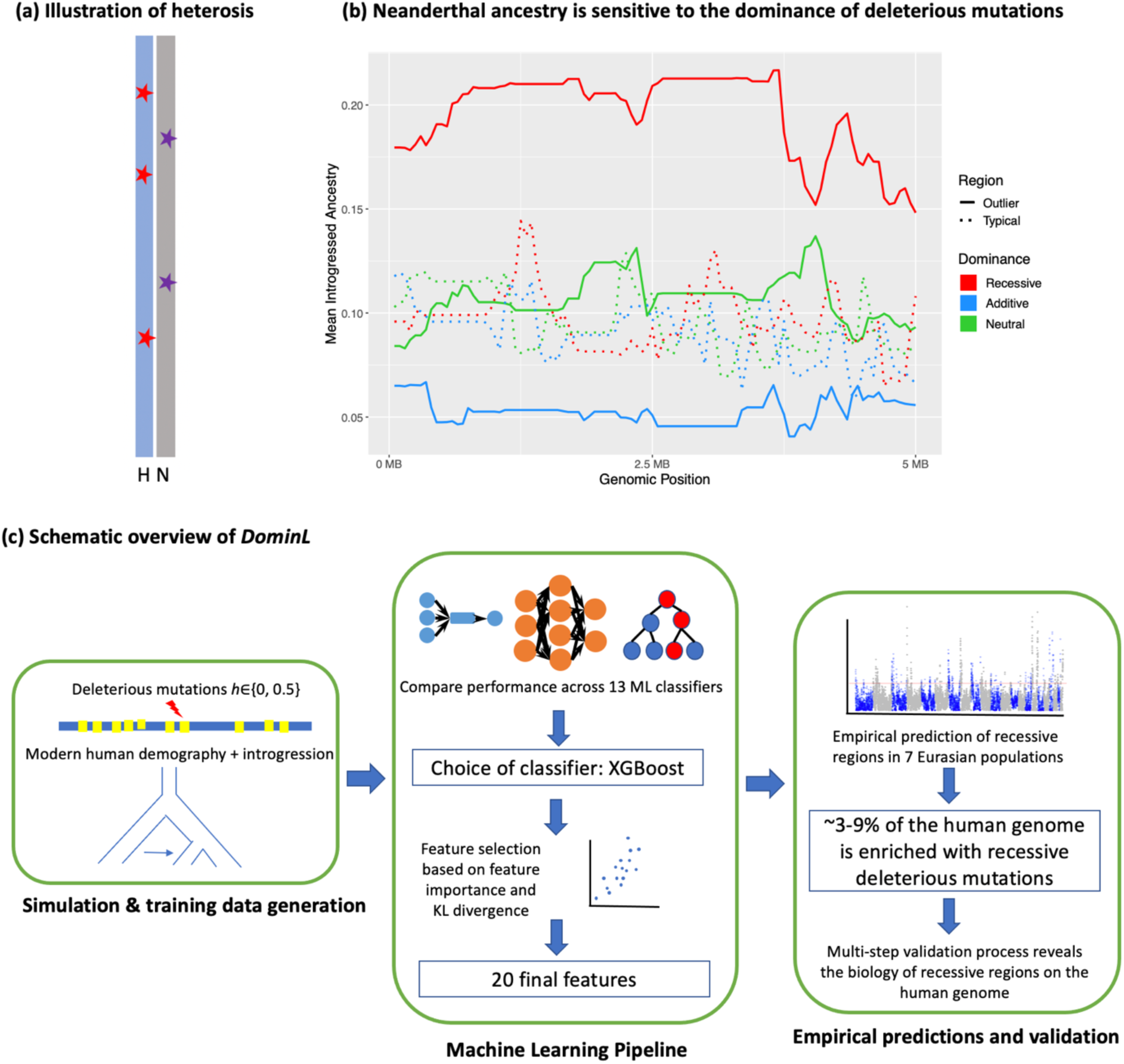
Overview of *DominL*. (**a**) The genetic mechanism of the heterosis effect. A Human(H)-Neanderthal(N) F1 hybrid individual carries distinct haplotypes. The stars represent recessive mutations private to human and Neanderthal populations, respectively. After Neanderthal introgression, the deleterious fitness effects of such variants are masked in heterozygotes, leading to an increase in Neanderthal ancestry in the human population. (**b**) Demonstration of the heterosis effect. Simulations show that an increase in Neanderthal ancestry in genomic segments with excessive exon density (“outlier”) with fully recessive deleterious mutations (red solid line), in contrast to segments with average exon density (“typical”, dashed lines), or those that accumulates neutral (green) or additive deleterious mutations (blue). (**c**) The schematic workflow of DominL. We use simulations of genomic segments with various distributions of exon density and recombination rates to compute Neanderthal ancestry and other informative statistics as features to identify the most optimized machine learning classifier that detects whether a genomic segment is enriched with recessive deleterious mutations. We applied the trained model to empirical human genomic data and validated the predictions through a multi-step process.

Here, we leverage this unique pattern of Neanderthal introgressed ancestry being sensitive to dominance, and develop a fundamentally different approach to detect recessive variants in human genomes. We train a supervised machine learning classifier called *DominL* (**Domin**ance **L**earner) to learn the evolutionary signature of recessive deleterious variants based on the heterosis mechanism. We then apply the classifier to map dominance effects along the genome, testing for 1Mb windows enriched with recessive deleterious mutations. *DominL* shows robust accuracy at detecting genomic regions enriched with recessive deleterious mutations at megabase resolution and has reliable power with high precision in regions with high exon density. We apply *DominL* to whole genome sequence data from individuals from 7 European and Asian populations from the 1000 Genome Project^32^, and show that approximately 3-9% of the human genome shows strong evidence of harboring recessive deleterious variants. Furthermore, these recessive regions are largely shared across human populations and are enriched in biological pathways related to vital metabolic and immune functions.

## Results

### Description of *DominL* and simulation of training data

The essential task for *DominL* is to solve a binary classification problem: whether a given genomic window is enriched for recessive versus additive deleterious mutations. Genome-wide analyses have shown that introgressed Neanderthal ancestry is generally depleted in humans due to negative selection against deleterious variants^33,34^. This depletion is expected under an additive model where archaic hominins with smaller effective population sizes are thought to carry more deleterious mutations than modern humans^26,34,35^, leading to strong purging of archaic ancestry. In contrast, if a region is enriched with fully recessive mutations private to modern humans, archaic introgression can mask these deleterious effects in hybrids, leading to relatively higher levels of archaic ancestry over time (the heterosis effect). Therefore, in genomic regions not affected by positive selection, variation in archaic ancestry is expected to reflect underlying dominance through the relative deviation from the additive baseline.

As a supervised machine learning classifier (Fig. 1c), *DominL* learns the signature of recessive and additive mutations by integrating multiple population genetic summary statistics, including the measures of archaic ancestry, genetic diversity, introgression, natural selection, and genomic structure (*i.e.* exon density, recombination rates). While Neanderthal introgression provides a central biological motivation for the framework, introgressed ancestry frequency is only one component of the information used by the classifier, consistent with feature-importance analyses shown in Supplementary Figure 7. *DominL* trains on extensive simulations of genomic segments sampled from a non-African human population that experienced Neanderthal introgression, where all deleterious nonsynonymous mutations are fully recessive (labeled as class “recessive”; *h*=0), or strictly additive (labeled as class “non-recessive”, *h*=0.5). Training on these extremes allows the classifier to learn the clearest boundary between dominance classes, and we subsequently assess classifier performance across a range of intermediate dominance values. The simulations are performed using the forward-in-time simulation tool SLiM^36^. The simulated evolutionary history is based on a demographic model of modern Europeans described by Gravel et al. 2011^37^ and Neanderthal introgression history inferred by Prüfer et al. 2014^29^, where the ancestral Eurasians received a single pulse of Neanderthal introgression after the Out-of-Africa migrations. For the simulated segments, we use the genic structure and recombination maps of 1000 randomly sampled 5MB windows across the human genome.

We divide each simulated segment into 1MB sliding windows (step size = 200kb), and compute the Neanderthal introgressed ancestry and other features (or summary statistics) in these windows. Selection coefficients for nonsynonymous mutations were drawn from a human-specific gamma distribution of fitness effect (DFE)^27^. For each set of simulations, the dominance of deleterious mutations is fixed as a constant randomly drawn at either *h*=0 (fully recessive) or *h*=0.5 (additive), making the ratio between the two classes approximately 1:1 in the training data. All genomic windows from a recessive simulation are labeled as class “recessive”. And similarly, all genomic windows from simulations drawn from an additive DFE are labeled as class “additive”. For each combination of dominance, segment, and demographic parameters, we obtain 100 replicates, and we combine the simulated data into a training data set of size 100,000 (n1=1e5). Separately, for examining the model performance, we obtain testing data set of smaller sample size (n2=1e3) using a different set of genomic segments unobserved in the training data.

### Selection of the ML classifier

To evaluate ML classifier performance, we used independent testing data that was not observed during the training phase. Receiver Operating Characteristic (ROC) curves and Precision-Recall curves were generated for each classifier, and Area Under the Curve (AUC) scores were stratified based on exon density of the genomic segments simulated (Fig. 2a, Fig. S1). We observed that classifier performance improved with increasing exon density, with the highest AUC values occurring in exon-rich regions. Additionally, we examined True Positive Rate (TPR) rankings at a fixed False Positive Rate (FPR) of 0.01, and assessed prediction consistency by calculating pairwise correlation coefficients between the predictions of different classifiers. Based on these criteria, 3 out of 13 classifiers emerged as top performers: XGBoost^38^, CatBoost^39^, and MLP^40^. These classifiers consistently demonstrated the highest power (TPR between 0.23 to 0.24) and highest precision (False Discovery Rate or FDR between 0.03 to 0.04) at FPR = 0.01 and exhibited strong agreement in their predictions across test datasets (Fig. 2b, Fig. S2-3). Given the difficulty of inferring dominance effects from population genetic data alone, this level of sensitivity at a stringent FPR indicates that *DominL* captures informative signals of recessive deleterious variants.

**Figure 2:**
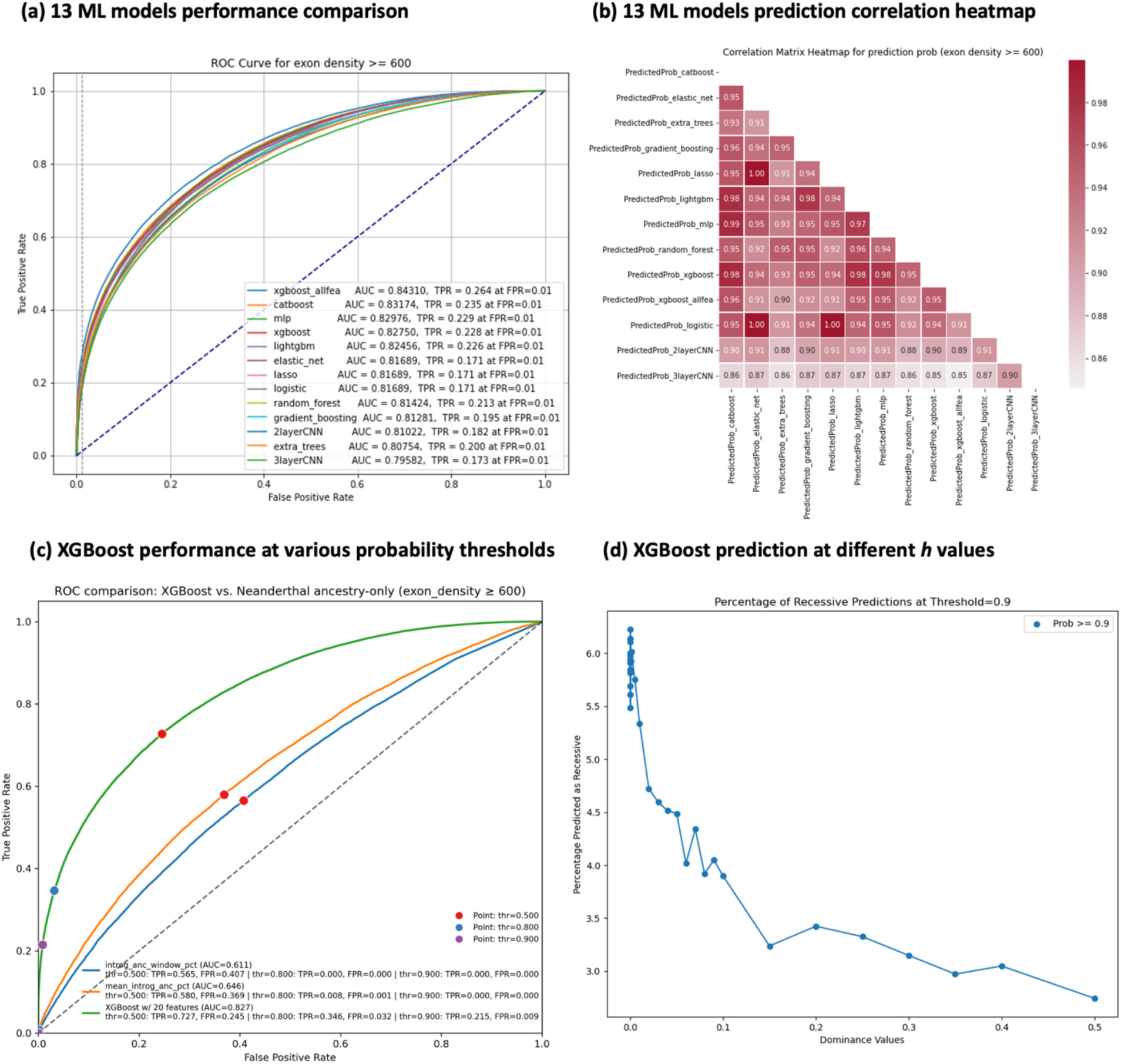
Evaluation of the accuracy and robustness of *DominL*. (**a**) Performance of 13 ML classifiers on simulated testing data generated from genomic windows with exon density larger than 600/5MB. We plot the Receiver Operating Characteristics (ROC) curves and highlight the Area Under the Curve (AUC) of each classifier as well as their power (True positive rate, TPR) when False Positive Rate (FPR) is capped at 1%. (**b**) Correlation matrix heatmap showing the concordance of the classifier predictions between pairs of classifiers on the same testing data. Prediction probabilities closer to 1 indicating higher consistency between classifiers. (**c**) Performance in terms of power and FPR at various prediction probability thresholds for the winning classifier, XGBoost (green line), comparing against prediction using Neanderthal ancestry only at 1MB resolution (blue line) or 5MB resolution (orange line). (**d**) The percentage of windows predicted as “recessive” using XGBoost at P_recessive_ > 0.9 when deleterious mutations have different dominance coefficients (x-axis). The proportion drops when h is larger than 0.1 for deleterious mutations, and at h=0.5, the proportion drops to 0.4%.

To further evaluate the performance of the ML classifiers, we applied the trained models to testing data stratified by exon density within a 5MB genomic window, with categories defined as fewer than 200 exons, more than 400 exons, more than 600 exons, and more than 800 exons per 5MB. We observe that classifier power improves as exon density increases. For example, in XGBoost, when the FPR is held at 0.01, the TPR increases from 0.12 when exon density is >200 per 5MB to 0.28 when exon density is >800 per 5MB (Fig. S4). Additionally, when the FPR is held at 0.01, the FDR remains consistently between 0.03 to 0.04 across all exon densities, showing high precision at detecting recessive windows (Fig. S4). Overall, all classifiers exhibit substantial power and high precision at detecting recessive deleterious mutations in regions with more than 600 exons per 5MB (Fig. 2a, 2c). We additionally evaluated whether dominance architecture could be resolved at finer genomic scales by retraining *DominL* on 100 kb windows derived from the same simulated 1 Mb segments. Although this approach increases spatial resolution, classifier performance was substantially reduced at 100 kb (lower ROC AUC; Fig. S5), indicating that the aggregated signal required to distinguish recessive from additive architectures is diluted at smaller scales. We therefore proceeded using 1 Mb as the minimal window size.

### Evaluation of classifier robustness

To further assess the generalizability of our trained classifiers, we tested their robustness by applying them to simulated data where deleterious mutations had intermediate dominance coefficients (0 < *h* < 0.5). Since the classifiers were originally trained on only the two extreme cases (*h* = 0 for purely recessive mutations and *h* = 0.5 for additive mutations), this step was critical in evaluating how well they could generalize across more biologically realistic intermediate dominance coefficients.

We assessed robustness by determining the proportion of intermediate *h*-value simulations classified as recessive. Among the three top-performing classifiers, XGBoost demonstrated the highest robustness in distinguishing intermediate dominance values. Specifically, when applying a stringent probability threshold (only assigning cases with probability ≥ 0.9 as “recessive”) to simulations where *h* < 0.1, we observe the highest percentage of *h* values being predicted as “recessive”. In contrast, when *h* > 0.1, the probability of a window being assigned as recessive rapidly declined (Fig. 2d). The other classifiers also performed well, but XGBoost exhibited the most precise separation between recessive and non-recessive cases across the intermediate *h* range (Fig. S6).

Based on this evaluation, we selected XGBoost as the final classifier of *DominL* for application to empirical data. The most reliable application of *DominL* is at 1MB windows, particularly in high exon density regions with more than 600 exons per 5MB. To further refine empirical predictions, we selected a probability threshold of *P_recessive_* ≥ 0.9 to maximize precision. We also performed a feature selection process to optimize the ML model performance (see Methods) and retained 20 out of 33 original features based on feature importance ranking and the Kullback-Leibler (KL)^41^ divergence of features between simulation and empirical data (Supplementary Table 1, Fig. S7-S9).

Given that Neanderthal ancestry is a key feature used by *DominL* and that true local ancestry is unknown in empirical data, we assessed the robustness of *DominL* to ancestry inference uncertainty. To do so, we applied the CRF-based ancestry inference framework of Sankararaman et al.^42^ to the simulated genomes and recomputed *DominL* features using the inferred ancestry tracts, instead of the true simulated ancestry. *DominL* predictions remained highly consistent under this setting, with similar or slightly lower false-positive rates and comparable power in exon-dense regions (Fig. S8). Importantly, using either true ancestry or inferred ancestry in the model, *DominL* significantly outperforms using Neanderthal ancestry-only as a predictor (Fig. 2c), indicating that it captures more complex signatures of dominance beyond simple shifts in archaic ancestry driven by heterosis.

To evaluate robustness to model misspecification, we applied *DominL* to additional simulated datasets with altered evolutionary parameters, including variation in recombination rates and the presence of positive selection. Classifier performance remained broadly consistent under these conditions, with similar ROC curves and only modest changes in power (Fig. S8). These results indicate that *DominL* captures signals specific to recessive deleterious variation rather than being driven by confounding effects of recombination rate heterogeneity or selective sweeps. We additionally evaluated false positive rates using neutral simulations lacking deleterious mutations. Although true positives are not defined in this setting, *DominL* showed an increase in false positive rate (FPR=0.085, compare to FPR=0.005 under the baseline model), as expected given the absence of purifying selection that distinguish additive from recessive dominance. However, it is important to note that such neutral scenarios do not recapitulate the consistent functional and evolutionary signatures observed in the empirical data, indicating that drift alone does not generate the recessive signal captured by *DominL*.

### *DominL* reveals recessive deleterious mutations in the human genome

Next, we applied the trained *DominL* classifier to the empirical modern human genomic data from 7 non-African populations from the 1000 Genomes Project (1KG), including 4 European populations (CEU, GBR, FIN, IBS) and 3 East Asian populations (CHB, CHS, JPT). These populations correspond to those for which CRF-inferred Neanderthal ancestry maps are available, providing high-confidence per-haplotype introgression calls required for computing *DominL* ancestry-based features. For each of the populations, we computed summary statistics in overlapping 1MB windows the same way as done for the training data, including genetic diversity-related statistics, exon density, and recombination rate. For the archaic human ancestry in genomic windows, we used the Neanderthal ancestry computed by Sankararaman et al.^43^ in the 7 non-African populations at the individual level, and averaged over all individuals in a population to obtain Neanderthal ancestry in 1MB windows for the population. We used a stringent prediction threshold (*P_recessive_* >0.9) in all populations, and converted the prediction probability to a log-scaled Recessive Score in Manhattan plots (Fig. 3a, Fig. S10, Supplementary Table 2). We further group the empirical predictions by whether the empirical genomic windows lie within 5MB genomic context where the exon density exceeds 600 exons. Additionally, we removed all adaptive introgression candidate regions^34,43–47^ from the genome-wide analyses, as archaic ancestry in such regions is driven by positive selection rather than deleterious mutations. The results below only apply *DominL* to the 22 autosomes.

**Figure 3:**
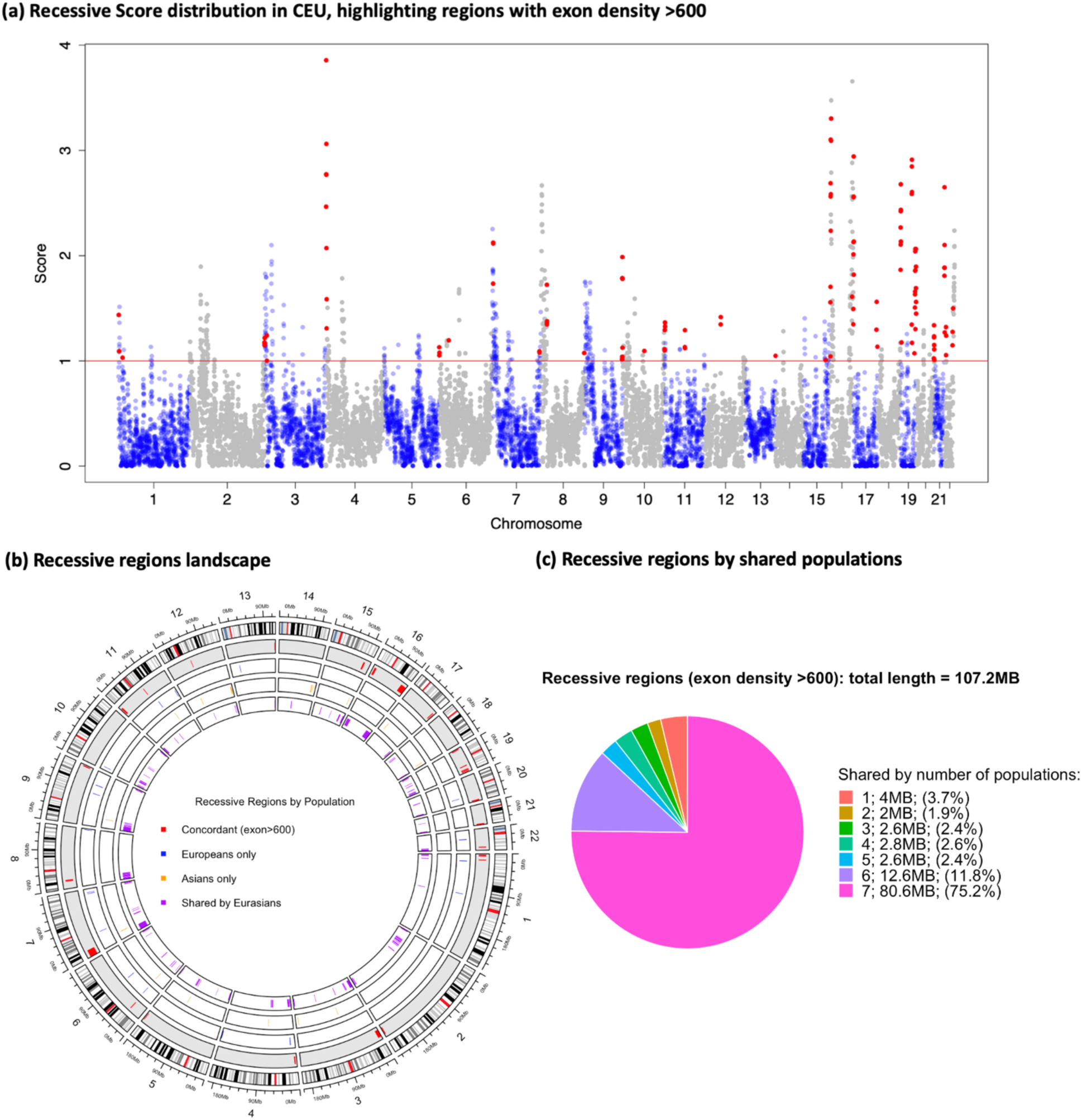
Modern human genomic windows enriched with recessive variants. (**a**) The genome-wide distribution of regions predicted as recessive in the CEU population (significance threshold as red horizontal line, which is defined as P_recessive_ ≥0.9), with red points highlighting regions with exon density >600/5MB. (**b**) The distribution of recessive genomic regions detected by DominL. The outermost circle shows the 22 autosomes. The second, gray circle shows the “concordant recessive regions” in red shades, defined by a region having exon density >600/5MB (where DominL has the highest performance) and is detected as recessive in at least 6 out of 7 Eurasian populations. The other circles show the recessive regions detected in Europeans (blue), East Asians (yellow), and at least one European and one East Asian population (purple) respectively. (**c**) Among the 107.2MB of genomic regions (exon density>600/5MB) detected as recessive by DominL, the vast majority are shared by nearly all non-African populations, and only a small fraction is detected in a single population or continent.

We concatenated the neighboring or overlapping windows predicted as recessive together for each population, and annotated these regions by the population in which they were detected. We further labeled the regions by superpopulations such as “European only”, “Asian only”, and “Shared by Eurasians” if a region was detected in at least one European population and one Asian population. Fig. 3b illustrates the distribution of the recessive regions across all 22 autosomes and the different superpopulations where they were found. We also highlight a category that we define as “concordant” that shows recessive segments that are found in at least 6 out of all 7 populations. In total, 249.2MB of the modern human autosomes exhibit evidence of recessive deleterious mutations (approx. 8.7% of the human autosomes). Under purely additive simulations, *DominL* yields a False Positive Rate of approximately 1% at the same prediction threshold. Therefore, if all genomic regions were truly additive, only 1% of the genome would be expected to be labeled as recessive by chance. The observed 8.7% genome-wide proportion of recessive regions substantially exceeds the baseline expectation, indicating that the empirical signal cannot be explained by false positive alone.

Of these windows showing recessive effects, 107.2MB (approx. 3.2% of the human autosome) fall within regions where the genomic context has exon density larger than 600 exons/5MB where *DominL* makes the most reliable predictions (Fig. 3c). Notably, exon-dense regions comprise 21.7% of the autosomes overall (Fig. S17), whereas 43.0% of recessive-predicted sequence lies within such regions, representing approximately a two-fold enrichment relative to the genomic baseline. Additionally, we show that among the 107MB in high exon density regions, 94.3MB are consistently detected across all human populations examined, meaning that the vast majority of high-confidence predictions are shared. While, the remaining windows were detected in only a subset of populations, we caution that such differences likely reflect variation in statistical power, demographic history, and stochastic variation rather than true population-specific dominance effects. For example, within Europe, the Finnish population (FIN) shows the largest number of uniquely detected windows (Fig. S11), whereas GBR shows few such cases. Given Finland’s demographic history characterized by bottlenecks and extreme genetic drift, this pattern is consistent with increased stochastic variation, rather than distinct biological enrichment. In East Asians, windows detected in only a subset of populations are distributed more evenly across groups (Fig. S11).

### Validation of empirical predictions

Given the difficulty of direct functional validation for the empirical predictions of recessive windows in humans, we attempted to validate our recessive predictions in three ways. First, we stratify the predictions into 3 categories depending on their recessive scores, including “Non-recessive” (*P_recessive_* ≤ 0.5), “Probable Recessive” (*P_recessive_* > 0.5 and < 0.9), and “Recessive” (*P_recessive_* ≥ 0.9). As a consistency check, we examined whether *DominL* predicted recessive windows exhibit higher Neanderthal ancestry, a pattern expected under the heterosis effect that motivated the framework (Fig. S12). Indeed, we observe exactly this pattern and find a positive correlation between the recessive scores and Neanderthal ancestry. We also examine the distribution of exon density in the 3 types of predictions (Fig. S13). We found that although *DominL* has the highest accuracy at detecting the recessive signature in exon-dense regions, there is little correlation between recessive scores and exon density, indicating the prediction of *DominL* is not biased toward exon-dense regions. These results are consistent with what is expected from the heterosis-based mechanism. However, because Neanderthal ancestry and exon density are features used by the classifier, these analyses should be interpreted as a reasonableness check rather than as independent validation.

Second, we asked how different pairs of the four key features related to the heterosis effect, including Neanderthal ancestry, heterozygosity, exon density, and recombination rates, correlate in the training data and the recessive simulations, and how that compares to the distribution of pairs of features in the empirical data and the empirical recessive predictions (Fig. S14). This comparison helps validate the method by confirming that the predicted recessive-enriched regions exhibit biologically plausible relationships between features that are broadly consistent with simulation expectations, while any discrepancies can be explained by known evolutionary forces absent from the simulation design.

In both simulations and empirical data, heterozygosity shows only weak association with exon density. In simulations, the Pearson correlation is r = 0.171 (Spearman rho = 0.156), while in empirical genome-wide data it is r = −0.039 (Spearman rho = −0.113). Although statistically detectable, given the large number of windows, these effect sizes are small in magnitude. The weak and inconsistent direction of association across simulated and empirical data indicates that heterozygosity is not strongly determined by exon density and contributes information beyond simple functional density. We also observe a slight positive correlation between Neanderthal ancestry and heterozygosity in both simulations and empirical data, which is also expected as we expect the recessive windows to contain higher Neanderthal ancestry due to heterosis. A slight negative slope in the empirical recessive windows can be explained by the particular high heterozygosity in a few windows at intermediate to low Neanderthal ancestry range. Additionally, in simulations, we expect more Neanderthal ancestry in exon dense regions, as half of the simulations contain only fully recessive mutations, which drives the heterosis effect. In the empirical data, however, we observe a strong negative correlation between exon density and Neanderthal ancestry, which has been repeatedly reported by various studies as most of the introgressed Neanderthal variants are selected against by negative selection^34,42,43^, thus more exons lead to stronger purging effect of the Neanderthal ancestry.

Finally, we examined whether windows predicted to have recessive deleterious mutations also showed evidence of non-additive effects for associations with 1,060 phenotypes in the UK Biobank (UKBB)^48^ (Fig. 4a) reported by Palmer et al.^48^ In that genome wide association study (GWAS), each variant was tested under both additive and non-additive models to identify loci where dominance effects beyond additivity contributed to phenotypic variation. Using their published summary statistics, we extracted variants with significant evidence of non-additive effects by retaining loci that pass a genome-wide Bonferroni cutoff under the non-additive model that also had smaller p-values than under the additive model. This filtering yielded 565,600 variants. If *DominL* correctly identifies genomic regions enriched for recessive deleterious mutations, we expect these regions to contain an excess of variants showing non-additive effects in phenotypic association tests. To test this, we quantified the number of UKBB non-additive variants overlapping each 1MB window. We determined the number of non-additive UKBB variants in all windows in the 3 categories of regions stratified by recessive scores, including “Non-recessive”, “Probable Recessive”, and “Recessive”. Non-additive UKBB variants are significantly enriched in the stringently recessive group when compared to the non-recessive and probable recessive groups (Wilcoxon p-values 3.95e-9 and 0.0043; Fig. 4a). To quantify the enrichment, we modeled variant counts using negative binominal regression adjusting for exon density, recombination rate, and GC content. We observed a strong enrichment of non-additive variants in *DominL*-predicted recessive windows (incidence rate ratio IRR = 1.48, 95% CI: 1.44-1.52), corresponding to a 48% more non-additive variants per window relative to non-recessive regions. In contrast, variants showing additive effects did not exhibit enrichment in recessive windows (IRR=0.93, 95% CI: 0.78-1.10), serving as a negative control (Fig. 4c). This analysis suggests that the signal captured by *DominL* is specific to non-additive genetic architecture rather than reflecting a general concentration of GWAS variants.

**Figure 4:**
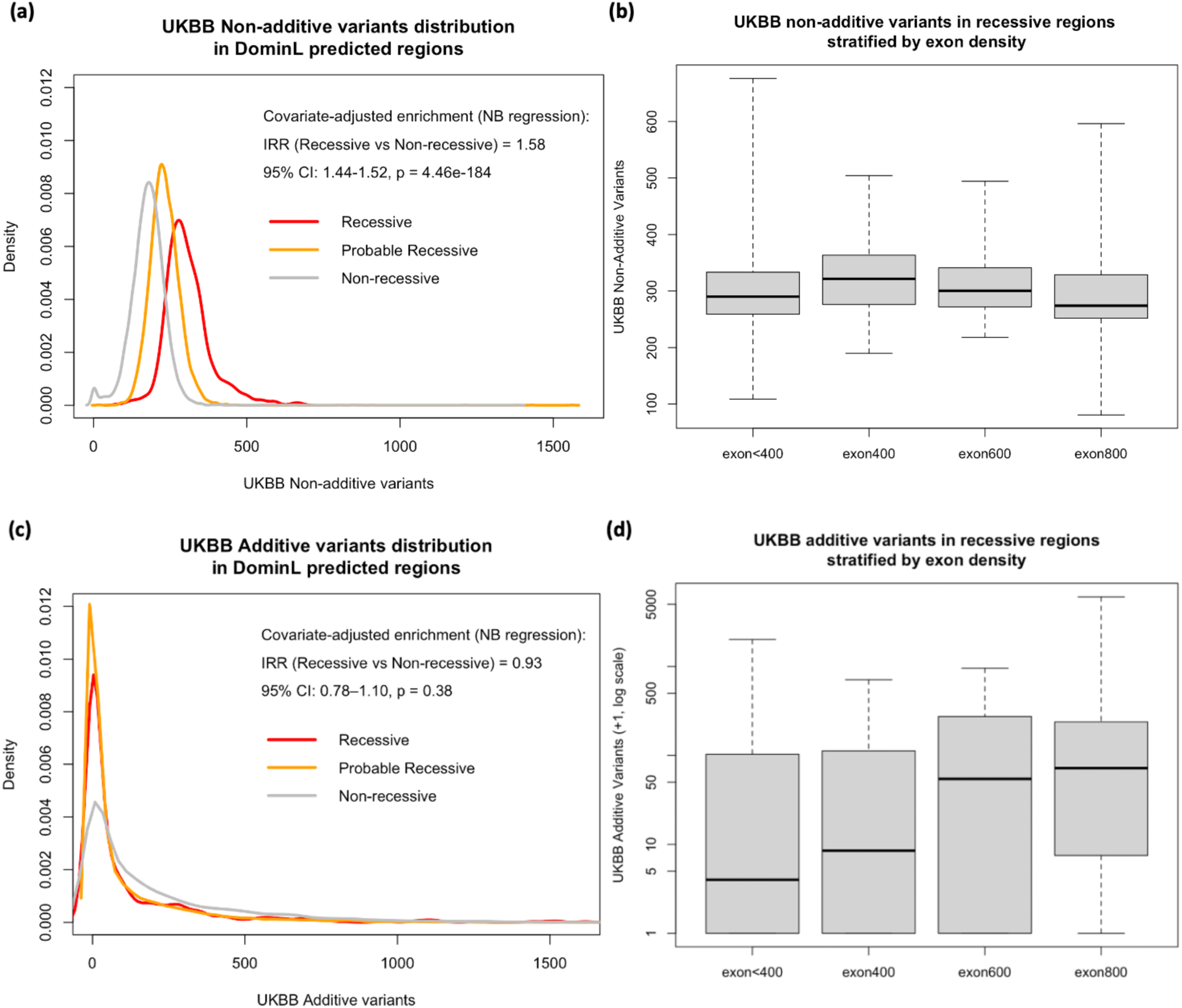
Predicted *DominL* recessive regions are enriched with UKBB non-additive variants, but not additive variants. We intersected the number of UKBB non-additive variants^48^ with the 1MB windows in our study, and examined the distribution of such variants in windows predicted as Recessive (P_recessive_ ≥ 0.9), Probable Recessive (P_recessive_ > 0.5 and <0.9), and Non-recessive (P_recessive_ ≤ 0.5). Panel **a** shows the distribution of UKBB non-additive variants significantly differs (via covariate-adjusted regression and block-based permutation) between any pair of DominL groupings. Non-additive variant counts were analyzed using negative-binomial regression adjusted for exon density, recombination rate, and GC content. Statistical significance was assessed using a two-sided Wald test (IRR = 1.48, 95% CI = 1.44-1.52, P = 4.46e-184). Panel **b** shows that such difference is not driven by varying exon density in genomic windows (Wilcoxon one-sided p-value between exon800 and exon<400 group = 0.130). On the other hand, as a control, we intersected the number of additive UKBB variants with the recessive windows (**c**). Additive variant counts were analyzed using the same negative-binomial regression framework; the association with recessive status was not significant (IRR = 0.93, 95% CI = 0.780-1.10, two-sided Wald P = 0.384). (d). Further, the density of variants showing additive associations increases with exon density. Sample sizes are n = 415, 108, 54, and 75 genomic windows for <400, 400–599, 600–799, and ≥800 exons per 5 Mb, respectively. Center lines indicate medians, boxes indicate the interquartile range (25th–75th percentiles), and whiskers indicate the minimum and maximum values.

Because *DominL* performs more accurately in exon-dense regions, we examined whether the enrichment of non-additive variants could simply reflect functional density. To test for this, we counted the number of trait associated variants in windows stratified by the number of exons. Reassuringly, the number of variants with non-additive effects shows little correlation with exon density in recessive windows (Fig. 4b, Fig. S15-16). The number of variants with additive effects, on the other hand, increases strongly with exon density (Fig. 4d), consistent with the known observations from GWAS^49,50^. This pattern suggests that the observed enrichment of non-additive variants is not driven by exon density alone. In sum, all three biological validation steps provide confidence that *DominL* picks up genuine signatures of recessive genetic effects.

### Biology of the recessive genomic regions

We next sought to understand the biology and genomic properties of the 3-9% of the human genome predicted to contain recessive deleterious mutations. Here, we focused on recessive windows shared across all Eurasian populations that also passed the filter of exon density greater than 600 exons/5MB as the test set, yielding a total of 94MB genomic sequence. To evaluate whether these regions exhibit distinct genomic or biological properties, we compared them to covariate-matched control windows sampled from the remainder of the genome. Control windows were selected to match key genomic features known to influence patterns of diversity and selection, including exon density, recombination rate, GC content, or heterozygosity, depending on the downstream analysis (see further in Methods section; Fig. S18). We then intersected these windows with other annotations to test specific biological hypotheses. For each annotation, statistical comparisons were performed using regression models that adjust for genomic covariates, and significance and effect sizes were estimated using block-based permutation and bootstrapping procedures to account for spatial correlation between windows (see Methods).

First, we hypothesize that runs of homozygosity (ROHs) would be depleted in genomic regions enriched with recessive deleterious mutations, because ROHs expose recessive deleterious alleles in homozygotes and are thus expected to be selected against. To test this hypothesis, we compared patterns of ROHs between recessive windows and covariate-matched control windows (see Methods). Because local heterozygosity strongly influences the distribution of ROHs, control windows for this analysis were matched to recessive windows on exon density, recombination rate, and heterozygosity. All analyses use ROHs inferred at the individual level in the 1KG populations (see Methods). We quantify three complementary ROH metrics within each genomic window: the number of individual runs overlapping the region (“number of runs”), the total length of all ROHs overlapping the region (“total length”), and the maximum ROH length overlapping the region (“max length”). To capture ROHs arising from different demographic processes, we evaluate these metrics separately for intermediate ROHs (0.5-2MB) and long ROHs (>2MB).

As expected, across all three metrics and both ROH length categories, recessive windows show a consistent significant depletion of ROHs relative to control windows (Fig. 5a, Fig. S19), indicating that genomic regions predicted by *DominL* to harbor recessive deleterious mutations are systematically less likely to form extended homozygous tracts. Interestingly, when we removed regions on chromosome 6 that overlap with the *HLA* gene cluster from both the recessive and control sets, the depletion of the intermediate and long ROHs became even more pronounced (Fig. S20). This observation is particularly interesting because the *HLA* region is known to harbor notably high levels of identity-by-descent (IBD) due to recent, strong natural selection^51–53^, which can lead to elevated ROH or IBD signals relative to the genomic background. Thus, while ROHs are generally depleted in genomic regions enriched for recessive variants, the *HLA* locus remains an exception, warranting further study.

**Figure 5:**
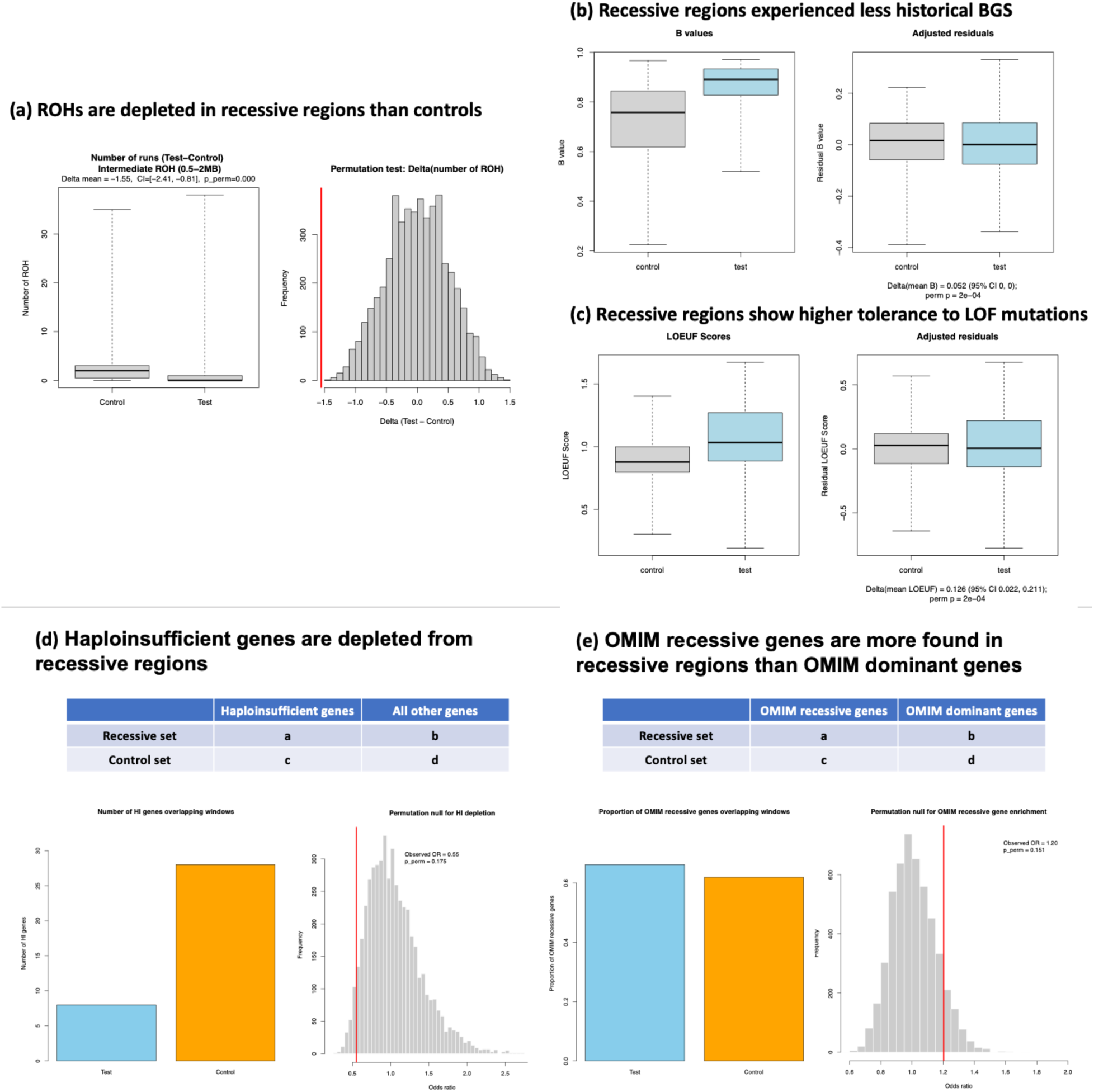
Multi-step validation process reveals the biology of recessive regions. (**a**) Number of intermediate-length ROHs (0.5-2MB) overlapping DominL-predicted recessive windows and covariate-matched control windows. Recessive windows show a significant depletion of ROHs. The left panel shows ROH counts per window, and the right panel shows the permutation null distribution of the mean difference (test-control), with the observed value highlighted in red. Statistical significance was assessed using a two-sided block-permutation test, with 95% CIs obtained by block bootstrap. Additional ROH metrics are shown in Supplementary Figure 19. (**b**) McVicker’s B values in recessive and control regions. A higher B value indicates weaker historical background selection. The left panel shows the raw values and the right panel shows covariate-adjusted residuals. (**c**) LOEUF in recessive and control windows. Higher LOEUF score indicates greater tolerance to LOF mutations. The left panel shows raw values and the right panel shows covariate-adjusted residuals. For panel b-c, differences between recessive and matched control windows were evaluated using covariate-adjusted linear regression; p-values were obtained using two-sided block-permutation tests and 95% CIs using block bootstrap. (**d**) Haploinsufficient genes overlap with recessive and control windows. Histogram shows permutation null distribution of the odds ratio for the contingency table, with the red line highlighting the observed value. (**e**) Relative abundance of OMIM recessive genes to dominant genes in recessive and control windows. Histogram shows permutation null distribution of the odds ratio for the contingency table, with the red line highlighting the observed value. For panel d-e, statistical significance of the observed odds ratio was assessed using a two-sided block-permutation test. For panels a-e, n = 123 for DominL-predicted recessive 1Mb genomic windows and n = 123 for covariate-matched control windows. In all boxplots, center lines indicate medians, boxes indicate the interquartile range (25th–75th percentiles), and whiskers indicate the minimum and maximum values.

To understand the evolutionary pressures and functional constraints affecting regions of the genome containing recessive deleterious mutations, we examined the distribution of McVicker’s *B* values^54^ and the LOEUF scores^55^. A lower *B* value indicates more intense historical background selection (BGS), while a lower LOEUF score indicates less tolerance to Loss-of-Function (LOF) mutations. We compared recessive windows to covariate-matched control windows and evaluated differences using regression models that adjust for exon density, recombination rate, GC content, and heterozygosity. We assessed statistical significance using block permutation and obtained confidence intervals using block bootstrapping (see Methods). We observe a significantly greater *B* values in the recessive set than the controls (Delta mean=0.052, permutation p=2e-4, Fig. 5b, Fig. S21), indicating weaker historical BGS in the recessive regions than the rest of the genome. This observation is consistent with the expectation that negative selection acts more efficiently at removing the additive deleterious mutations than recessive deleterious mutations^56,57^. Furthermore, previous work has noted that the BGS signal is stronger in genomic regions without associative overdominance^57^, which also is concordant with what we observe that recessive, non-overdominant regions experienced less BGS. Similarly, we observe significantly higher LOEUF scores in the recessive windows compared to the control windows (Delta mean=0.126, permutation p=2e-4, Fig. 5c).

Because higher LOEUF scores indicate greater tolerance to LOF mutations, this result suggests that recessive regions are overall less functionally constrained and are more tolerant to gene-disrupting variation.

Next, we examine the distribution of the nearly 300 known haploinsufficient genes in humans^58^. Haploinsufficient genes, where a single functional copy is insufficient to maintain normal function, are usually caused by LOF mutations and are typically associated with dominant diseases^59^. Because such genes are expected to exhibit dominant rather than recessive effects, we tested whether they are underrepresented in *DominL*-predicted recessive regions using a gene-centric null. We compared the frequency of HI genes overlapping recessive windows relative to all other genes overlapping recessive windows. Statistical significance was assessed using block permutation tests (see Methods). Although the depletion does not reach statistical significance (permutation p=0.175), which likely is due to the limited power from relatively small number of haploinsufficient genes, the observed odds ratio is below 1 (OR=0.55), indicating a directional depletion of haploinsufficient genes in recessive regions (Fig. 5d). This pattern is consistent with the expectation that genomic regions enriched for recessive variants are less likely to contain genes whose pathogenic effects are primarily dominant.

Similarly, we examine the distribution of autosomal recessive and autosomal dominant genes as annotated by the OMIM database^60,61^. If *DominL* correctly identifies regions enriched for recessive variants, we expect those regions to contain relatively more autosomal recessive genes and fewer autosomal dominant genes. Because disease genes tend to reside in highly constrained genomic regions^62^, we focused on the relative abundance of OMIM recessive versus dominant genes between recessive windows and matched control windows. Using a contingency table framework, statistical significance was assessed using block permutation. Although the permutation test did not reach statistical significance (permutation p=0.151), the observed odds ratio is greater than 1 (OR=1.20, Fig. 5e), indicating that OMIM autosomal recessive genes are relatively more common than autosomal dominant genes within *DominL*-predicted recessive regions. Given the limited number of curated OMIM disease genes, this analysis likely has limited statistical power. Nevertheless, the directional trend is consistent with the expectation that genomic regions enriched for recessive deleterious mutations are more likely to harbor genes associated with recessive rather than dominant diseases.

We additionally noted that *DominL*-predicted recessive regions were disproportionately located near chromosome ends (Fig. S22). This pattern may in part reflect genomic structure associated with model power: across the genome, highly exon-dense windows show a modest tendency to occur closer to chromosome ends, while *DominL* has substantially greater power in such exon-dense regions. To determine whether chromosome-end localization could account for the empirical properties of predicted regions, we repeated the downstream analyses while additionally including distance to the nearest chromosome end as a covariate, otherwise retaining the analysis framework and test/control regions used above (Fig. S22; Supplementary Table 3). The elevated B values and enrichment of non-additive UK Biobank associations remained significant after this adjustment. ROH depletion was attenuated, but remained negative across all six measures, with maximum overlapping ROH length remaining significantly depleted for both intermediate and long ROHs. The positive LOEUF shift was also attenuated and was no longer statistically significant after chromosome-end adjustment, although the directionality of effect remained unchanged. Overall, these sensitivity analyses indicate that chromosome-end localization does not fully account for the empirical patterns associated with *DominL*-predicted recessive regions and does not provide a sufficient alternative explanation for the inferred recessive signal.

Lastly, we investigate the biological pathways that are most influenced by recessive deleterious mutations. We use the program Enrichr^63^ to perform a Gene Ontology enrichment analysis for biological processes and molecular functions respectively, and visualize the top 10 pathways from each of the enrichment test. We show that for the recessive regions shared in all Eurasian populations, 25 biological processes yielded significant p-values from the enrichment test, with the most prominent processes being vital metabolism-related functions, particularly in Oxygen Transport functions (Fig. 6a, Supplementary Table 4). On the other hand, only 8 molecular functions yielded significant p-values from the enrichment test (Fig. 6b, Supplementary Table 5), with the most significant enrichment coming from olfactory receptor activities, followed by immune-related activities such as the MHC receptors, which are generally all related to individual fitness. When we examine the recessive regions that are found in only one population or one continent (Fig. S23), the most biological functions showing the strongest enrichment appear to be less shared between populations, and are overall mostly related to functions that influence oxygen transport, immune functions, and metabolism.

**Figure 6:**
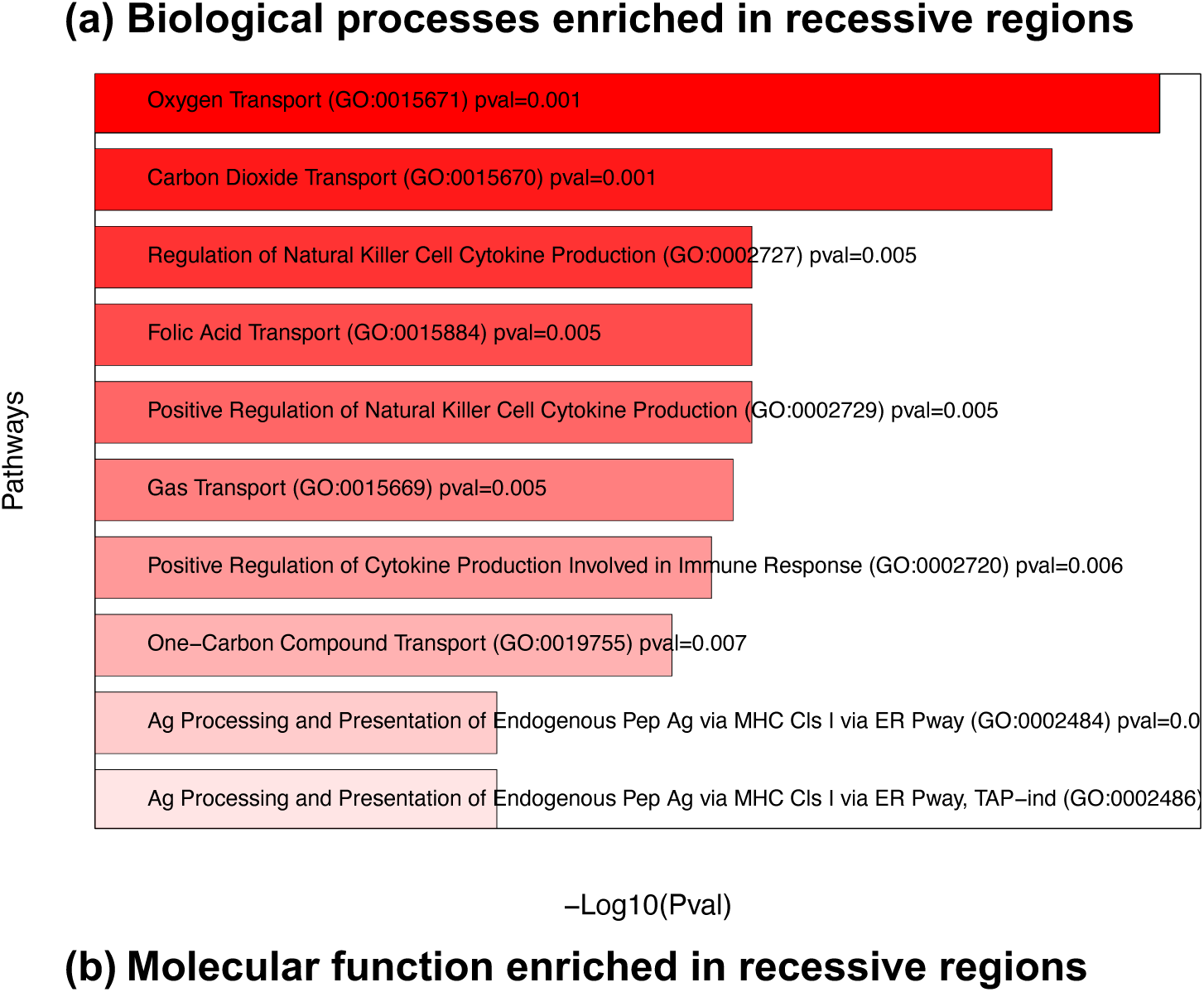
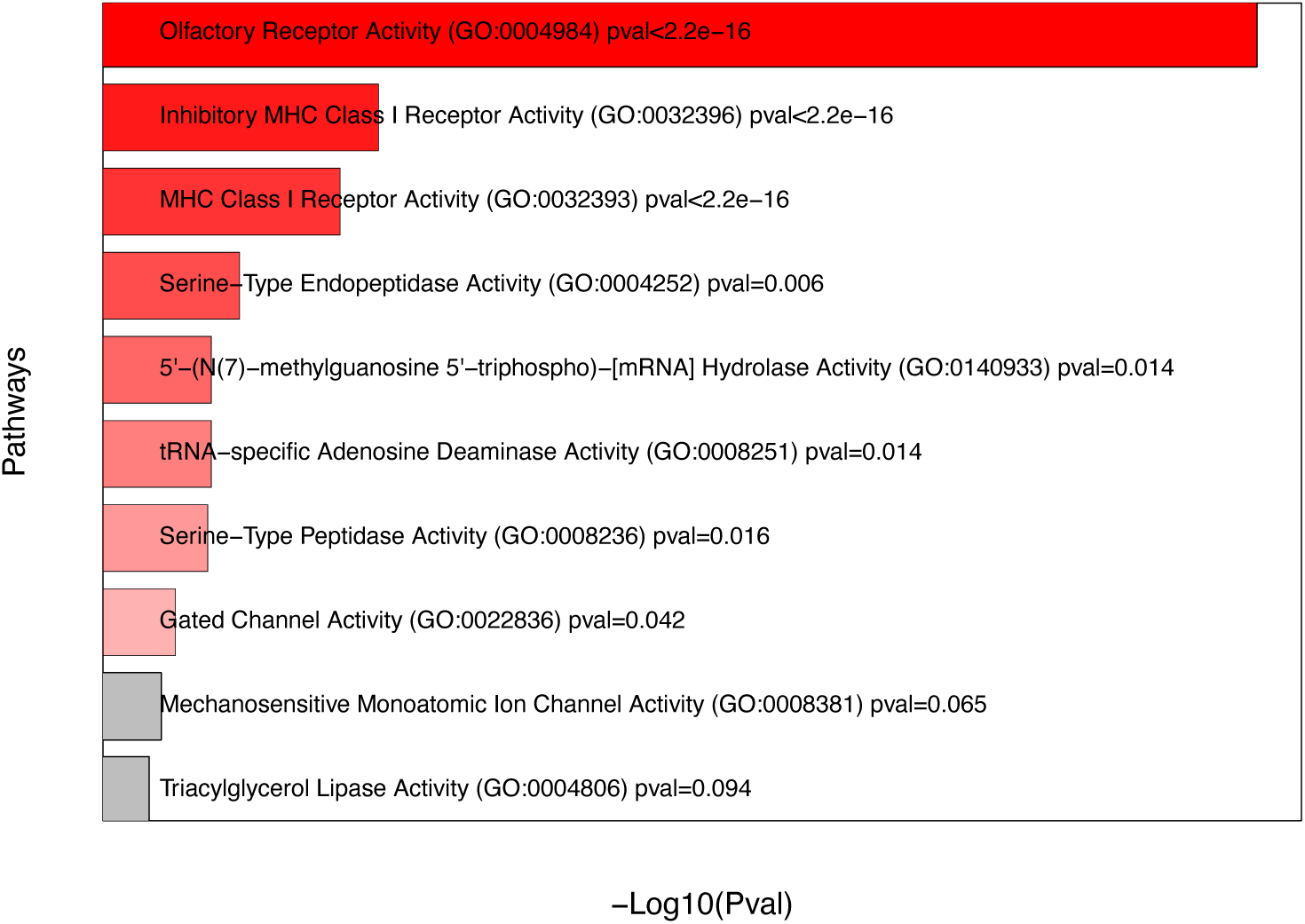
Recessive regions are mostly found in biological functions related to vital metabolic and immune functions. Gene Ontology (GO) enrichment analyses using Enrichr for human biological processes (**a**), and molecular functions (**b**). The top 10 processes are shown, with ascending p-values for each analysis, with red indicating a significant p-value and gray indicating non-significant p-values. The intensity of the red color as well as the length of the bars both indicate how significant a process is from the enrichment test. All p-values are corrected for multiple testing using Benjamini-Hochberg False Discovery Rate^64^ method. GO enrichment was performed using Enrichr using a one-sided Fisher’s exact test for over-representation. P-values were adjusted for multiple testing using the Benjamini-Hochberg false discovery rate procedure.

## Discussion

Dominance is one of the most fundamental concepts in genetics and has many implications in population and evolutionary genetics^1–8^. Despite its importance, dominance is poorly understood for most mutations in the human genome. Previous studies of dominance have been underpowered to detect dominance effects separately from selective effects. Indeed, models where all deleterious mutations were assumed to be additive provided an adequate fit to the human SFS for nonsynonymous mutations, suggesting that either additive or recessive models could explain the data^4,25,65^. In this work, by leveraging the distribution of Neanderthal introgressed ancestry, we present a novel machine learning-based method, *DominL*, to test for recessive deleterious mutations in the human genome. Our study provides strong statistical support for the presence of recessive deleterious mutations in the human genome.

The signal leveraged by *DominL* arises from how dominance shapes the fate of introgressed ancestry. While our framework is motivated by the contrast between masking of recessive deleterious mutations and the purging of additive mutations, it is important to note that heterosis does not reverse the overall depletion of archaic ancestry, but instead modulates its extent across the genome. In this context, *DominL* captures the relative deviation from the additive expectation, rather than absolute increases in archaic ancestry, providing a genome-wide signal of the dominance distribution.

By applying *DominL* to 1KG non-African modern human populations, we provide striking and compelling evidence that recessive deleterious mutations are present in the human genome and are not uniformly distributed. Instead, they are concentrated in specific genomic regions, covering up to 9% of the autosome - or 3% when restricting to regions where *DominL* performs with the highest accuracy. Importantly, at the selected prediction threshold, the number of regions identified far exceeds what would be expected from false positive predictions alone, indicating that the observed signal cannot be explained by chance. Because our inference is based on a stringent threshold designed to maximize precision, these estimates are likely conservative, suggesting that the true extent of recessive-enriched regions is likely even larger. These results point to a substantial and non-uniform concentration of recessive deleterious mutations shaping human genomic variation.

Our finding of recessive deleterious mutations has significant implications for understanding the genetics of complex traits, especially in improving polygenic models for complex phenotype prediction. Traditional quantitative genetic methods used in human genetics model phenotypes assuming strict additivity among variants associated with the trait^66–71^. For example, widely-used polygenic risk scores (PRS) typically assume variants act in an additive manner. Further, empirical data across traits and species suggest that most of the genetic variance is additive^72–75^. However, inconsistency in phenotype and heritability estimation across populations and study cohorts has been frequently reported^76–78^, and recessive effects on common diseases have been reported in small populations with high autozygosity^79,80^. Although classic studies using deviation tests led to a widely accepted conclusion of mainly additive genetic variance in complex traits^72^, such studies only consider the non-additive effects from trait-associated SNPs, which omits the effects from genomic backgrounds harboring these SNPs. If some trait-associated variants, especially those with larger effect sizes on the trait, fall in recessive windows, phenotypic prediction accuracy may be improved using a recessive model than traditional additive models. Future studies could incorporate empirical dominance maps to refine these models and improve their predictive power.

*DominL*’s reliability is further validated by its consistency with biological predictions from empirical data, which remain robust under covariate-matched controls and block-based resampling tests. We observe that genomic regions with an excessive amount of recessive deleterious variants generally tend to harbor fewer haploinsufficient genes, more autosomal recessive genes, and show a strong depletion of intermediate and long runs of homozygosity (ROHs). Moreover, the recessive regions exhibit higher tolerance to loss-of-function (LOF) mutations, suggesting that recessive mutations in these areas are less constrained and more tolerant of functional disruptions. We also found that recessive regions experienced weaker historical background selection (BGS), confirming what we expect from the efficiency of negative selection at removing recessive deleterious mutations in contrast to additive or dominant mutations. These observations affirm *DominL* as a powerful and reliable tool for identifying genomic regions enriched with recessive deleterious mutations. Although the current ML classifier used in *DominL* has a resolution limitation of 1MB, this does not diminish the significance of the predictions. Given the limited knowledge about the genomic distribution of recessive mutations in humans, our findings provide a valuable foundation for future research.

One notable property of *DominL*-predicted regions is their enrichment near chromosome ends. High exon density regions are themselves modestly enriched toward chromosome ends, and because *DominL* has greater power in exon-dense regions, this genomic structure may contribute to the localization of high-confidence predictions. However, sensitivity analyses explicitly accounting for distance to the nearest chromosome end showed that several of our principal empirical observations, including elevated B values and enrichment of non-additive UK Biobank associations, remained robust. ROH depletion was partially attenuated but remained consistently negative across all measures, with maximum overlapping ROH length remaining significant for both intermediate and long length categories. The LOEUF association was more sensitive to chromosome position but remained directionally consistent. We therefore interpret chromosome-end localization as an important genomic context of *DominL* predictions, rather than as a potential confounder.

*DominL*’s results also reveal insights about the connection of evolutionarily deleterious mutations to phenotypes. In general, the relationship between a mutation’s effect on phenotype and its effect on fitness is complicated, and may depend on the particular trait being considered and its connection to fitness^81,82^. Here, we found that regions of the genome enriched with recessive deleterious mutations also show more non-additive variants associated with complex traits in UKBB. This suggests that the historical fitness effects being quantified through patterns of Neanderthal introgression are concordant, at least broadly, with the phenotypic associations with complex traits in present-day populations. This suggests a connection between the evolutionary effects of mutations and their effect on phenotype.

The distribution of recessive mutations also reveals a shared evolutionary history among human populations, with substantial overlap of recessive regions in different populations across continents. This shared history is particularly evident in regions with high exon density, where *DominL* demonstrates the greatest power. These regions likely accumulate more mutations in all human populations, leading to a stronger heterosis effect upon archaic introgression.

One of the most interesting findings is the signal observed in the *HLA* region. This region’s enrichment with recessive mutations and the heterosis effect may have significantly influenced its genetic diversity. Our evidence suggests that the repeatedly reported balancing selection signature in *HLA*^28,52,53,83,84^ may partly reflect the presence of recessive deleterious mutations in the region. However, unlike other regions, the *HLA* region is not depleted of long ROHs like the rest of recessive regions. This anomaly appears compatible with what was previously noted by Albretchsen et al. 2010^51^ that *HLA* region contains excessive IBD shares in humans due to strong recent selection at this locus, thus can lead to more ROHs in subsequent generations. This explanation suggests unique evolutionary pressures and mechanisms at play in the *HLA* region, warranting further investigation to fully understand its genetic dynamics. Our analysis also shows that many recessive windows are enriched in vital biological pathways especially in functions related to metabolism. This enrichment highlights the potential role of recessive mutations in shaping metabolism and fitness functions. Understanding the implications of these findings could provide valuable insights into the genetic basis of many complex diseases.

Looking forward, there is room for improvement to *DominL*. Currently, *DominL* predicts at 1MB resolution, which is much larger than the size of a gene. To enhance the resolution of these predictions, alternative approaches such as deep learning methods could be explored for finer-scale inference, which has the great potential at revealing more detailed insights into the distribution and impact of recessive mutations on individual genes. Additionally, since *DominL*’s prediction mechanism relies on the heterosis effect upon archaic introgression, it currently is limited to application in non-African populations. Furthermore, given that we rely on populations with high-resolution Neanderthal ancestry maps, the application of *DominL* is further limited to Eurasian populations. Extending the analysis to other populations, such as recently admixed American populations, would require substantial methodological modifications. For example, Neanderthal ancestry in American populations is distributed across heterogenous local ancestry backgrounds, thus disentangling recessive effects associated with introgressed ancestry from such populations would therefore require explicit modeling of admixture history. It will be highly valuable to identify alternative methods or features to adapt *DominL* for predicting in other non-Eurasian populations, including the Americans and African populations or even archaic hominins, as the knowledge of dominance distribution can facilitate a broader understanding of the patterns and evolution of recessive mutations throughout human evolution, and represents an important direction of future work.

We also note that non-additive genetic effects can also arise from gene-environment (GxE) interactions, in which allelic effects vary across environmental contexts. In such cases, an allele may appear recessive in some environments but more additive in others, particularly in loci involved in immune response (e.g. *HLA*). *DominL* does not explicitly model GxE interactions and therefore captures an effective, population-averaged recessive enrichment inferred from introgression patterns. While this averaging mitigates highly context-specific effects, future extensions incorporating environmental or population-specific contexts will be important for refining dominance inference.

In conclusion, *DominL* provides a novel and powerful framework for exploring the landscape of recessive deleterious mutations in the human genome, filling a critical knowledge gap in human genetics. Our findings highlight the non-uniform distribution of these mutations, their evolutionary conservation, and their potential implications for genetic diversity and disease. Future research leveraging *DominL*’s results and expanding its applications will continue to reveal the complexities of recessive genetic variation and facilitate numerous evolutionary genetics and disease genetics studies.

## Methods

### Features used by *DominL*

In addition to Neanderthal ancestry, we computed a set of biologically meaningful summary statistics as features for training the *DominL* classifier. These features are computed in sliding 1MB windows (step size 200kb) throughout the simulated segments, and they generally capture information about genetic variation (e.g. Watterson’s θ, sequence divergence, Tajima’s *D*, heterozygosity), natural selection (eg. Haplotype diversity, Garud’s H12 statistics), and archaic introgression (eg. The *F* statistics, ABBA-BABA’s *D*, the *U* and *Q* statistics), recombination rate, and genic structure. Overall, these features provide insight into the change in genetic variation after Neanderthal introgression, and provide quantitative measures as to how the allele frequency distributions shift due to subsequent evolutionary changes such as negative selection, the heterosis effect, and demographic events. Additionally, we incorporate recombination rate and genic structure (e.g. exon density) as features, as their distribution can heavily influence the magnitude of the heterosis effect.

A complete list of the features can be found in Supplementary Table 1, and all features are computed in Python3.

### ML classifier implementation and selection for *DominL*

The ultimate goal for *DominL* is to distinguish and predict genomic windows as binary classes “recessive” versus “additive”, where a recessive prediction is interpreted as a genomic region enriched with recessive deleterious variants. To identify the most effective machine learning (ML) classifier for detecting signatures of heterosis upon Neanderthal-human introgressed segments due to recessive deleterious mutations, we implemented and compared 13 different approaches. These included CatBoost^39^, Elastic Net^85^, Extra Trees Classifier^86^, Gradient Boosting^87^, Lasso^88^, LightGBM^89^, Multi-Layer Perceptron (MLP)^40^, Random Forest^90^, XGBoost^38^, Logistic Regression^91^, and two convolutional neural network (CNN)^92^ architectures (a 2-layer and a 3-layer CNN). All classifiers were trained using the same set of training data, which consisted of simulated genomic datasets under two selection regimes: one where all deleterious mutations were additive (*h* = 0.5) and another where all were recessive (*h* = 0). Models were evaluated using held-out test datasets generated under the same simulation framework. Genomic windows were classified as recessive when *DominL* prediction probability exceeded 0.9.

### XGBoost overview and advantages

XGBoost (eXtreme Gradient Boosting)^38^ is an optimized gradient boosting framework designed for high performance and efficiency. It builds a series of decision trees sequentially, where each new tree corrects errors made by the previous trees through gradient descent optimization. Regularization mechanisms, including L1 and L2 regularization, are incorporated to prevent overfitting and enhance classifier generalization. XGBoost can automatically learn the best split direction for missing values, making it robust for real-world datasets with incomplete information. Its efficient parallelization and memory optimization make it significantly faster than traditional boosting methods.

XGBoost offers several advantages, including high predictive accuracy due to ensemble learning, built-in feature selection through feature importance evaluation, robustness to multicollinearity and missing values, and efficient handling of large datasets through parallel computing. Additionally, it provides interpretability through feature importance scores. However, it is computationally intensive for very large datasets, requires careful hyperparameter tuning for optimal performance, and can be prone to overfitting if not regularized properly. XGBoost is particularly effective for feature-based supervised learning because of its ability to capture complex feature interactions while maintaining high efficiency. Its regularization mechanisms prevent overfitting, making it suitable for genomic datasets where high-dimensional features often pose challenges. Furthermore, XGBoost’s ability to rank feature importance ensures that the most informative genomic features contribute to classifier predictions, improving overall robustness and interpretability. These attributes make it the ideal choice for our analysis of recessive deleterious mutations and their genomic signatures.

To estimate computational requirements, we monitored training time and peak memory usage during model fitting. Training was performed using the XGBoost implementation in Python. Runtime was measured using time.perf_counter, and peak resident memory usage (RSS) was recorded using the psutil library by sampling memory usage at 0.1-second intervals during training. Training times and peak memory consumption were recorded for the primary *DominL* model as well as alternative feature configurations. We show that training the *DominL* classifier using XGBoost on the simulated dataset (∼187,000 windows with 20 features) required approximately 9 seconds on a single CPU core, with peak memory usage of ∼1.6 GB. Model inference is computationally inexpensive: once features are computed, prediction for all genomic windows in a population requires less than one second. Runtime scales approximately linearly with the number of genomic windows and genomes analyzed.

We also note that our XGBoost model used the default hyperparameters of the Python package. We evaluated several alternative hyperparameter configurations including tree depth, learning rate and number of boosting rounds, and observed minimal differences in predictive performance while substantially increasing computational cost and training time. We therefore retained the default parameters to avoid overfitting the classifier to specific simulation settings.

### Feature selection process

To optimize the selected classifier for empirical applications, we performed a feature selection process focusing on two primary criteria: (1) features most frequently utilized by the ML classifiers, and (2) features with comparable distributions between simulated and empirical data. During classifier development, we observed that several features had drastically different distributions in empirical datasets compared to simulations, raising concerns about prediction accuracy due to domain shift. To address this, we computed the Kullback-Leibler (KL)^41^ divergence between feature distributions in the training simulations and empirical data, retaining only features with low KL divergence values to ensure comparability between the two datasets.

Following this selection process, we retained 20 out of the original 33 summary statistics (Supplementary Table 1). We then assessed the impact of feature selection by comparing classifier performance using all features versus the reduced feature set. For example, in the case of XGBoost, using the full feature set led to lower accuracy and reduced consistency in predictions on test data compared to the refined subset. This confirmed that removing incompatible features improved classifier robustness and interpretability without sacrificing predictive power.

In the primary simulation-based evaluation, true local ancestry from the simulations was used when computing ancestry-based features. To assess the potential impact of ancestry inference uncertainty, we performed an additional validation in which simulated genomes were processed using the CRF-based ancestry inference framework of Sankararaman et al. (2014) (details see below), and inferred ancestry tracts were used in place of true ancestry when computing *DominL* input features. Classifier performance under this setting remained comparable to analyses using true ancestry, with similar or slightly lower false-positive rates and comparable power in exon-dense regions (Fig. S8), indicating that ancestry inference uncertainty does not materially affect *DominL’s* predictions.

### CRF Neanderthal ancestry inference

Neanderthal ancestry was estimated across simulated 5 Mb genomic segments using a Conditional Random Field (CRF) approach^42^. Each segment comprised 92 diploid individuals from four populations: 2 Neanderthal, 30 African, 30 European, and 30 Asian, with European individuals treated as the admixed target population and African and Neanderthal individuals serving as unadmixed reference panels. Prior to calling, duplicate chromosomal positions were resolved by retaining a single representative variant per site, genotypes at triallelic sites were recoded to the reference allele, and genetic distances were assigned to each variant using per-1 Mb bin recombination rates from the simulation metadata. The CRF was run in inference mode with pre-trained model parameters over 100 kb windows, producing a matrix of posterior probabilities of archaic origin for every variant–haplotype pair. Two summary statistics were then derived from this matrix: mean archaic ancestry, defined as the simple mean of all posterior probabilities, and a thresholded estimate obtained by classifying each variant– haplotype pair as Neanderthal-derived when its posterior exceeded 0.5 and reporting the resulting fraction. Both statistics were computed over the full 5 Mb segment as well as within each of five non-overlapping 1 Mb bins, and all results were stratified by segment identity, replicate, and dominance coefficient.

### Selecting Control Windows

To evaluate biological properties of *DominL*-predicted recessive regions, we compared these regions to covariate-matched control windows sampled from the remainder of the genome. The test set consisted of genomic windows predicted to be recessive by *DominL* (prediction probability> 0.9) that were shared across Eurasian populations and with exon density exceeded 600 exons per 5 Mb.

Covariate balance between recessive and control windows was evaluated using standardized mean differences (SMD), defined as the difference in group means divided by the pooled standard deviation. SMD values less than 0.1 indicate good covariate balance. Diagnostic plots showing the distributions of exon density, recombination rate, GC content, and heterozygosity before and after matching are shown in Fig. S18.

We found that exon density and recombination rate could be matched very closely between recessive and control windows (|SMD| ≤ 0.1). However, because recessive windows are concentrated in a narrow genomic subset of the genome, perfect matching on both GC content and heterozygosity simultaneously was not feasible. When matching on exon density, recombination rate, and GC content, residual imbalance remained for heterozygosity but was substantially reduced compared to the unmatched dataset (SMD = 1.55 versus 1.97 previously). Conversely, when matching on exon density, recombination rate, and heterozygosity, GC content showed only moderate residual imbalance (SMD = 0.82 versus 1.60 before matching). In all cases, the unmatched covariate exhibits substantially improved balance relative to the original analysis.

Because different downstream analyses are sensitive to different genomic features, we constructed two complementary control sets: 1) For annotation-based analyses including background selection, LOEUF scores, OMIM genes, and haploinsufficient genes, we used control windows matched on exon density, recombination rate, and GC content. GC content is directly associated with mutation rates and background selection and is therefore the most relevant covariate for these analyses. 2) For runs of homozygosity (ROH), we used control windows matched on exon density, recombination rate, and heterozygosity, as heterozygosity is a primary determinant of ROH distribution. In both sets of controls, windows were sampled randomly from the remainder of the genome while matching the empirical distribution of the specific covariates.

Residual differences in unmatched covariates were small (low SMD values) and were additionally controlled for by including these covariates as predictors in regression-based statistical models. To account for spatial correlations among adjacent genomic windows, statistical inference was performed using block-based resampling. Blocks consisted of contiguous genomic segments spanning 94 MB (matching with the size of recessive windows shared across all populations), and 10,000 bootstrap and permutation replicates were performed to estimate effect sizes, confidence intervals, and empirical p-values. For metrics with continuous distributions (eg. B values, LOEUF scores), effect sizes were defined as the mean difference between recessive and control windows. For metrics with binary status (eg. number of genes overlapping windows), we constructed 2×2 contingency tables and effect sizes were defined as the odds ratios. For metrics with continuous values, we additionally fit linear regression models including exon density, recombination rate, GC content and heterozygosity as covariates, and used the residuals for comparison between recessive and control windows.

### Runs of Homozygosity inference

We identified identity-by-descent (IBD) segments and runs of homozygosity (ROH) in empirical modern human genomic data from the 1000 Genomes Project (1KG)^32^, focusing on four European populations (CEU, GBR, FIN, IBS) and three east Asian populations (CHB, CHS, JPT). We used the 1KG data where variants were called using human genome build 19 and we filtered the VCF to include only biallelic sites. We used version r1206 of IBDSeq^93^ to identify both IBD segments and ROH. IBDSeq is a likelihood-based method designed to detect IBD segments in unphased sequence or array data. ROH are also output, because they are a special case of IBD within an individual rather than between individuals. We split the data by population and ran IBDSeq with default parameters, rather than running the method jointly across the entire 1KG dataset. ROHs were grouped as intermediate (0.5-2MB) or long (>2MB), and are evaluated for their overlap with *DominL*-predicted recessive windows. Because ROH boundaries are determined during the inference step rather than binning windows based on physical distance, converting physical distance to genetic distance would not alter the segments or enrichment analyses here.

To quantify ROH enrichment, we summarize ROHs using three complementary metrics. First, we counted the number of ROH under a given length category overlapping each genomic window (number of runs). Second, we calculate the total genomic lengths of ROHs overlapping each window (total length of runs). Third, we measure maximum length of a single ROH overlapping each window (max length of runs). These metrics capture complementary aspects of homozygosity, including ROH frequency, cumulative length, and maximum coverage. Enrichment analyses were performed by comparing a given ROH metric in recessive windows to covariate-matched control windows (matching heterozygosity as mentioned above) using block permutation and bootstrap. In the main text, we present the intermediate ROHs in terms of number of runs, and we show the additional metrics and long ROH results in the Supplementary Materials.

### Application to empirical human genomic data

We used the phased variant calls from the 1000 Genomes Project Phase 3 dataset (GRCh37/hg19). Variants were filtered to retain biallelic SNPs only, and all other variant types (e.g, multiallelic sites and indels) were excluded to ensure consistency with the simulation-based feature definitions. No additional filtering on minor allele frequency (MAF), Hardy–Weinberg equilibrium (HWE), or missingness was applied, as *DominL* relies primarily on window-based summary statistics that are robust to rare variants. All summary statistics were then computed directly from the filtered dataset using the same procedures applied to the simulated data.

### UK Biobank GWAS variant enrichment analysis

To evaluate whether *DominL*-predicted recessive regions show enrichment for trait-associated variants, we analyzed genome-wide association results from the UK Biobank (UKBB) reported by Palmer et al. 2023, which contains summary statistics of GWAS on 1030 fitness-related traits testing both the additive genetic effects and non-additive effects. GWAS variants were mapped onto the windows used throughout *DominL* analyses. For each window, we counted the number of: 1) non-additive GWAS variants: these are variants that show significant non-additive p-values when fitted under a non-additive model, and these p-values also needs to be more significant than the additive p-values; and 2) additive GWAS variants: these are variants that show significant additive p-values that are more significant than the non-additive p-values.

We then compared the abundance of these variants between *DominL*-predicted recessive windows and control windows. To quantify enrichment, we fit generalized linear models in which the number of variants per window was modeled as a function of window class. Effect sizes were summarized using the incidence rate ratio (IRR), representing the relative rate of GWAS variants in recessive windows compared to matched control windows. Models also included covariates including exon density, recombination rate, GC content and heterozygosity to control for genomic structure. To account for spatial correlation among windows, statistical significance was assessed using the block-based permutation and bootstrap framework described above. Both additive and non-additive variant counts were also examined as a function of exon density to assess whether signals from non-additive variants could alternatively be explained by underlying genomic structure rather than recessive mutation enrichment.

## Data Availability Statement

All datasets analyzed in this study are publicly available. Phased variant calls from the 1000 Genomes Project Phase 3 dataset (GRCh37/hg19) were obtained from the International Genome Sample Resource (IGSR). CRF-based Neanderthal ancestry maps were obtained from Sankararaman et al. (2014). Additive and non-additive UK Biobank GWAS summary statistics were obtained from Palmer et al. (2023). Gene-level loss-of-function constraint metrics (LOEUF) were obtained from gnomAD v2.1.1. Background-selection B values were obtained from the precomputed McVicker et al. background-selection map. Haploinsufficient gene annotations were obtained from the supplementary data of Dang et al. (2008), and autosomal dominant and recessive disease-gene annotations were obtained from Online Mendelian Inheritance in Man (OMIM). Source data underlying the figures are provided with this paper.

## Code Availability Statement

Custom code used to perform the simulations, calculate summary statistics, train and evaluate the *DominL* classifiers, generate empirical predictions, and perform downstream analyses is publicly available in the DominL GitHub repository: https://github.com/xzhang-popgen/DominL

## Acknowledgement

The authors thank Jeffrey Kidd, the Zhang Lab at the University of Michigan, and the Lohmueller Lab at UCLA for helpful discussions during the development of this project.

## Author Contributions statement

X.Z. conceived and designed the study, developed the original computational framework, performed and directed the analyses, interpreted the results, and wrote the manuscript. L.Z. and J.Y. implemented and evaluated the machine-learning methods, analyzed model performance, and applied the trained models to empirical data; L.Z. additionally performed extensive reanalysis and validation of the machine-learning pipeline and evaluated the models on additional test datasets. H.Z.W. implemented the CRF-based Neanderthal ancestry inference. N.S. performed exploratory simulations and generated Fig. 1b. J.M. provided the ROH data and contributed to the ROH methodology and Methods description. S.S. supervised the machine-learning components of the study and contributed to interpretation of the results. K.E.L. supervised the overall study and contributed to study design and interpretation of the results. All authors reviewed and approved the manuscript.

## Completing Interest Statement

The authors declare no competing interest.

