## Supplementary material for "Neanderthal introgressed ancestry reveals human genomic regions enriched with recessive deleterious mutations": 260826SI-Dominance.pdf

#### Supplementary Figures and Tables

##### Supplementary Figure 1: Precision-Recall curves of all ML models at exon density >=600

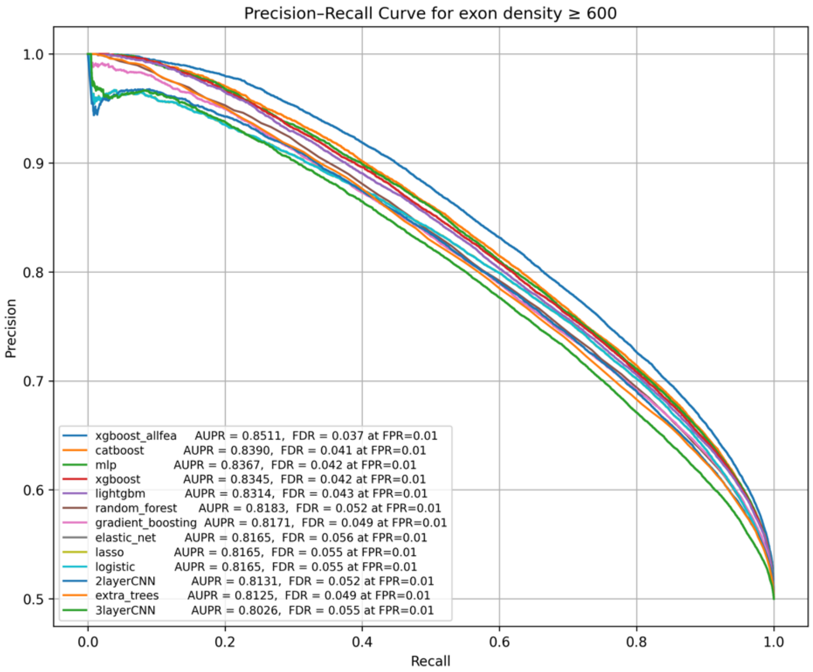


*We evaluate the performance of 13 ML classifiers on simulated testing data generated from genomic windows with exon density larger than 600/5MB. We plot the Precision-Recall curves and highlight the Area Under the Curve (AUPR) of each classifier as well as their False Discovery Rates (FDR) when the False Positive Rate (FPR) is capped at 1%.*

##### Supplementary Figure 2: Pairwise ML model comparison on empirical data using CEU and exon density >=600

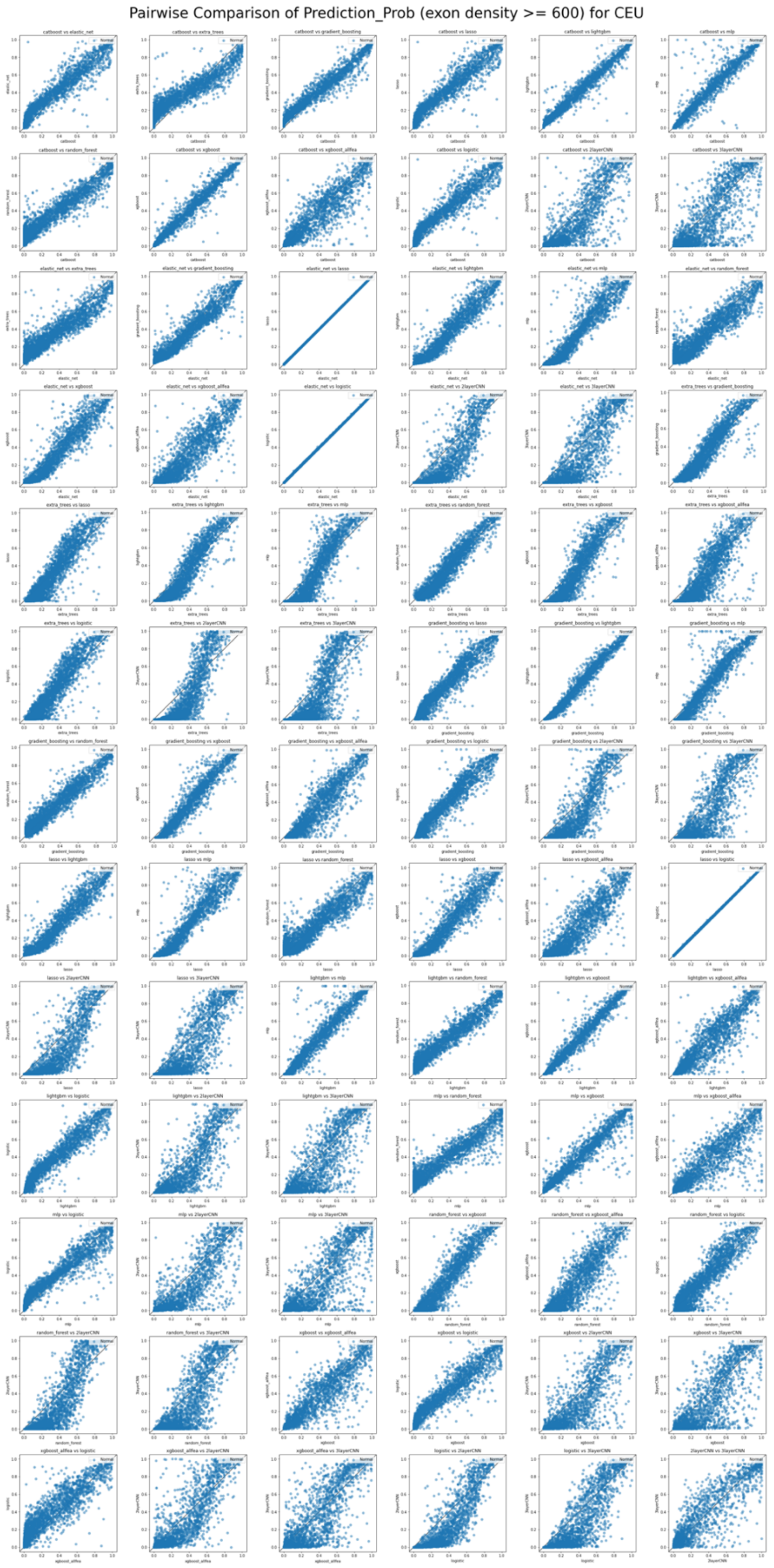

We *evaluate the consistency of predictions between classifiers by plotting the prediction probabilities of all empirical windows (filtered by exon density >=600) from CEU between different pairs of classifiers.*

#### Supplementary Figure 3: Correlation matrix between pairwise ML algorithms, stratified by the exon density distribution

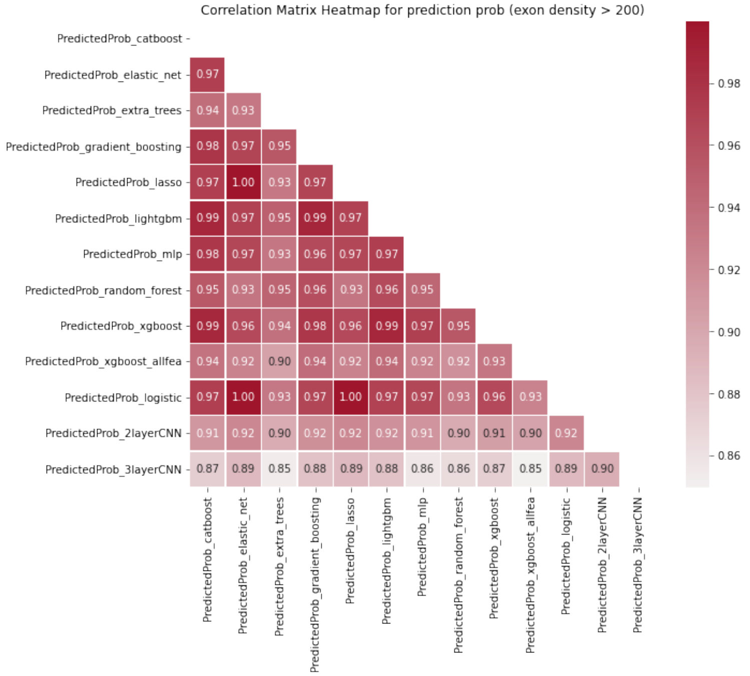


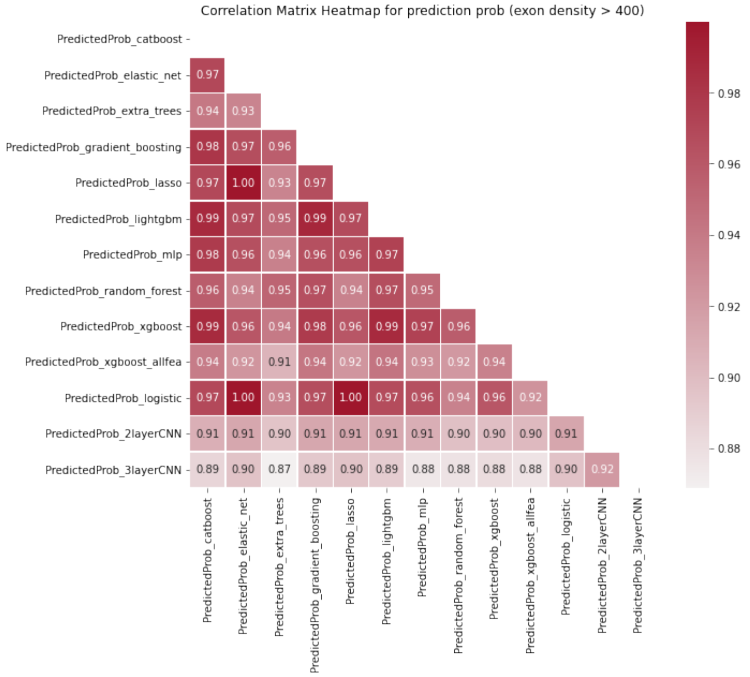

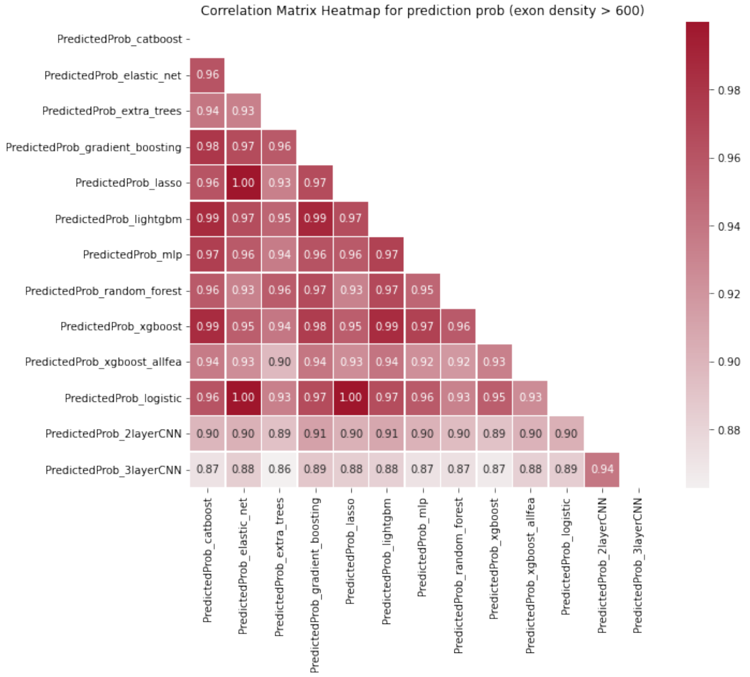


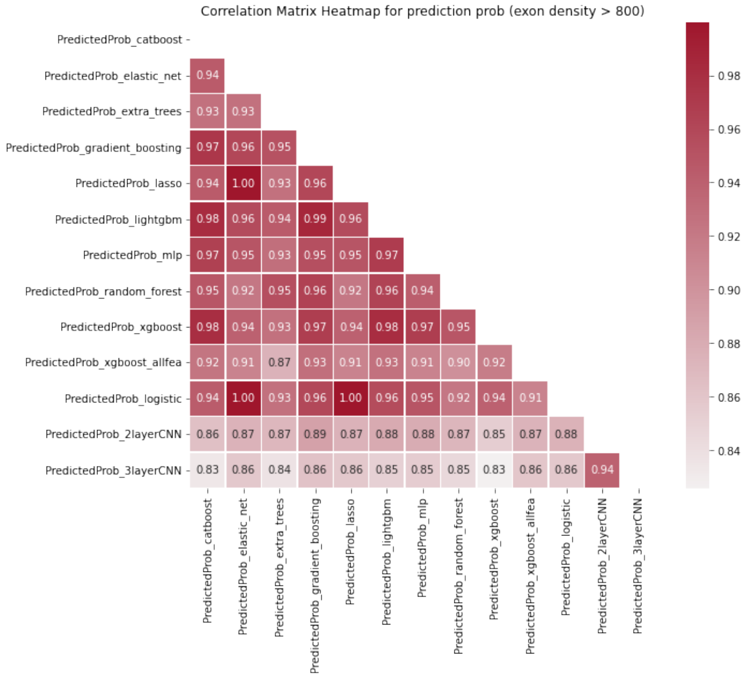


*To evaluate the consistency of predictions between classifiers, we use the prediction of all models on the same testing data, and plot the correlation matrix heatmap, with the color gradient representing the strength of linear correlation. From top to the bottom, each panel represents a correlation matrix heatmap for a given exon density threshold, including 200, 400, 600, and 800 exons per 5Mb.*

**Supplementary Figure 4: ROC and Precision-Recall curves of XGBoost at different exon density thresholds**

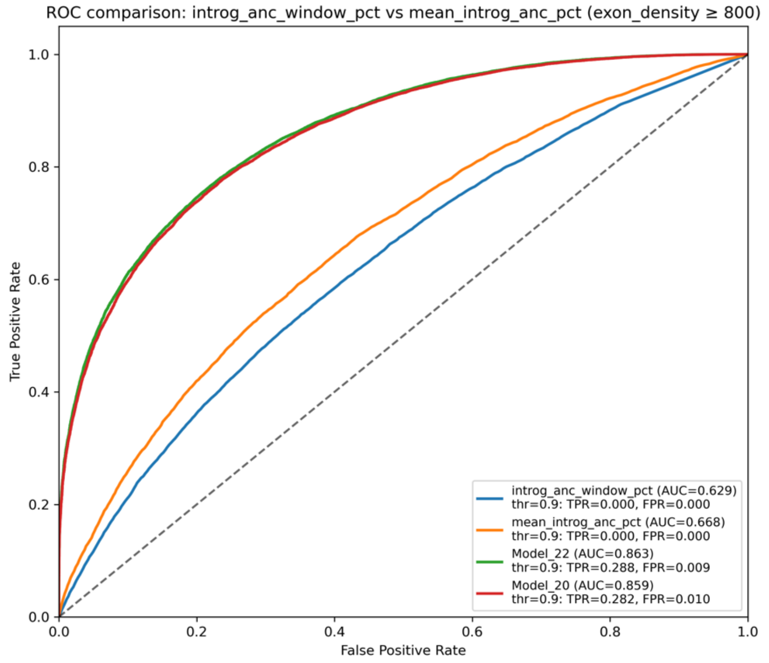


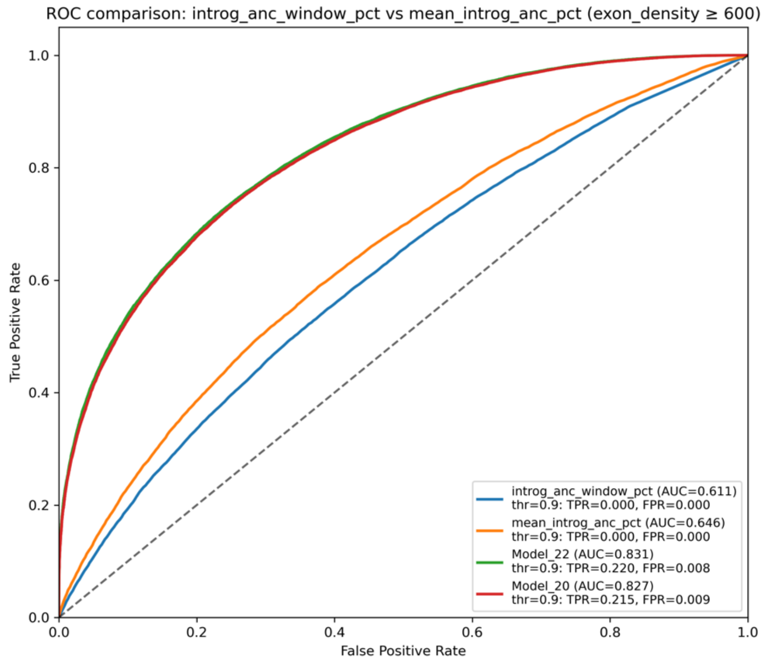

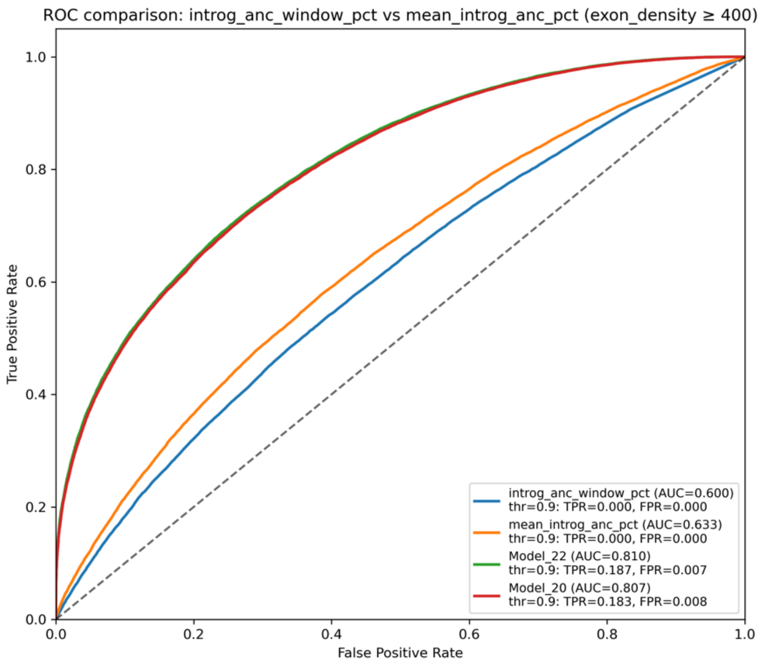


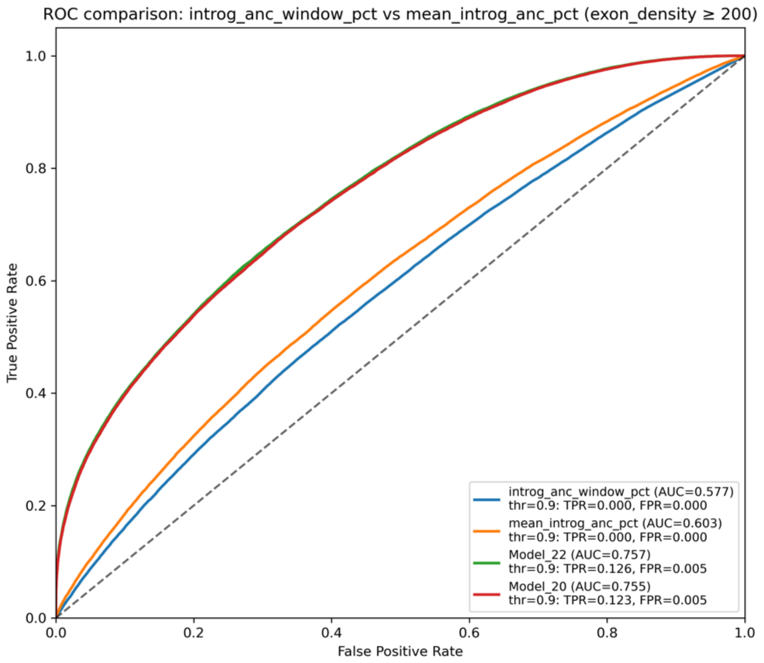


*We show the performance of XGBoost ("Model20" is the reported DominL model, red, and Model22, green, is retrained model after adding two recombination rate related features) in terms of ROC and Precision-Recall curves at various exon density thresholds, including 200, 400, 600 and 800 exons per 5Mb. On each ROC curve, we highlight the power (TPR) and False Positive Rates (FPR) at different prediction probability thresholds. We compare the performance of XGBoost against using Neanderthal ancestry-only as predictor (Neanderthal ancestry in 1MB as blue, and Neanderthal ancestry in 5MB as orange).*

**Supplementary Figure 5: ROC and Precision-Recall curves of XGBoost in different window sizes, using exon density >=800**

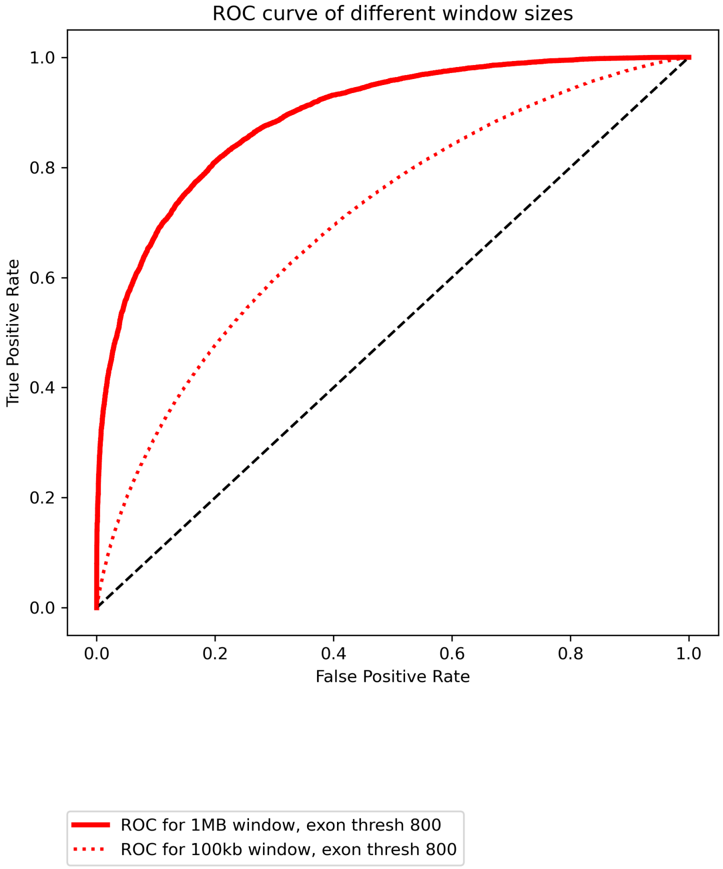

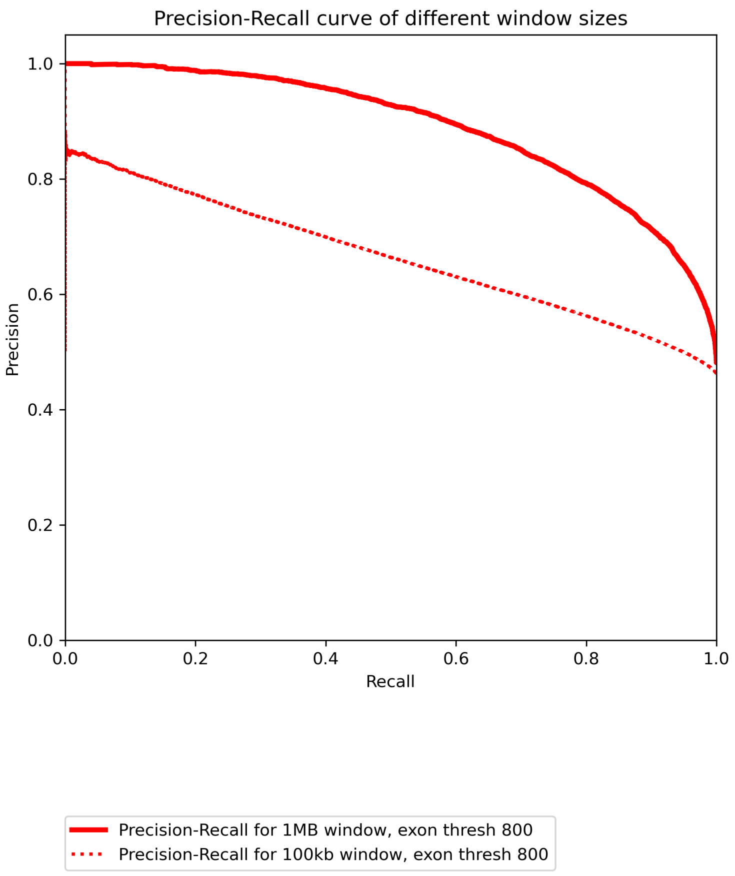


*We determine the most ideal window size used in DominL by showing the ROC and Precision-Recall curve of trained XGBoost classifier on 1MB windows versus 100kb windows, using exon density threshold of 800 exons per 5Mb. We show that the accuracy of predictions is substantially better in MB-resolution windows than smaller, kb-resolution windows.*

**Supplementary Figure 6: ML models prediction at different *h* values**

**a) Catboost**

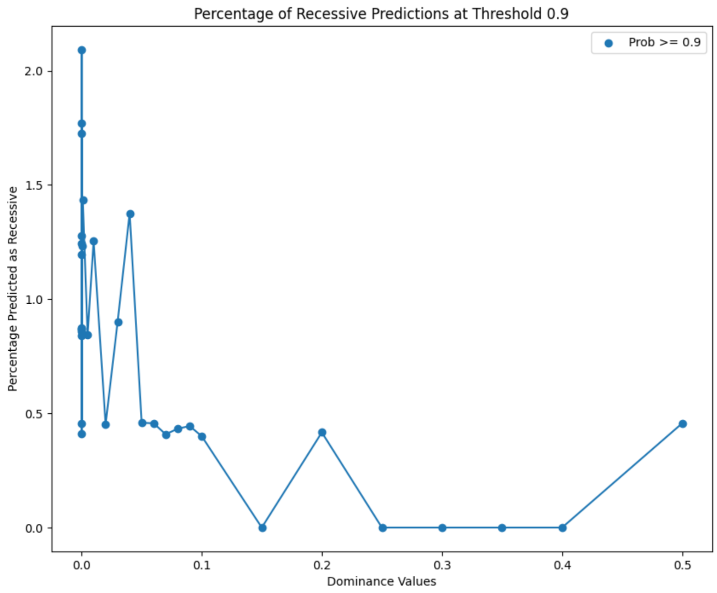


**b) Mlp**

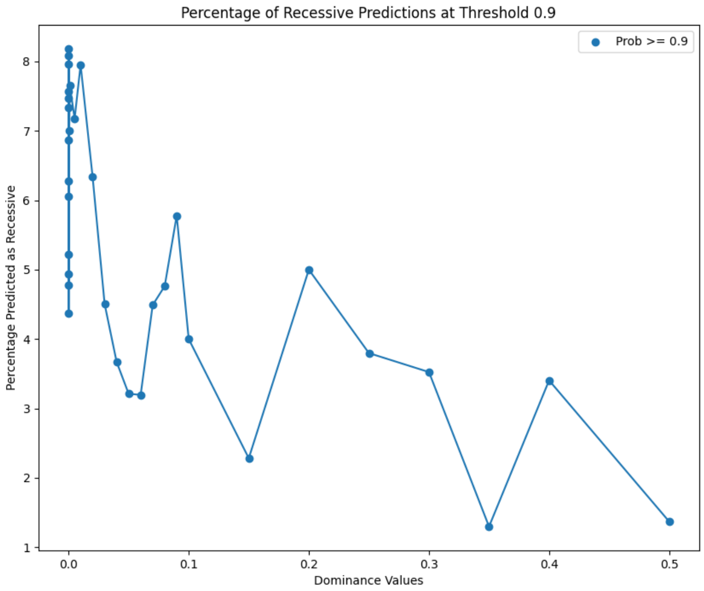


**c) Extra Tree Classifier**

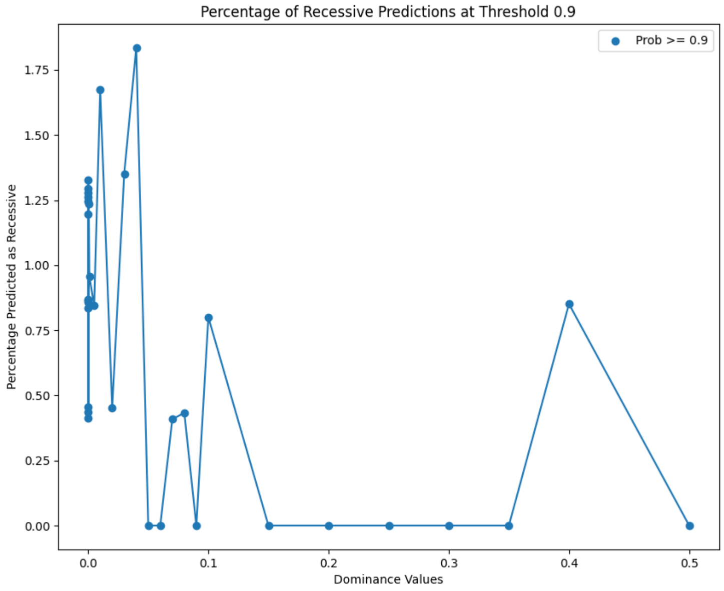


**d) Gradient Boosting**

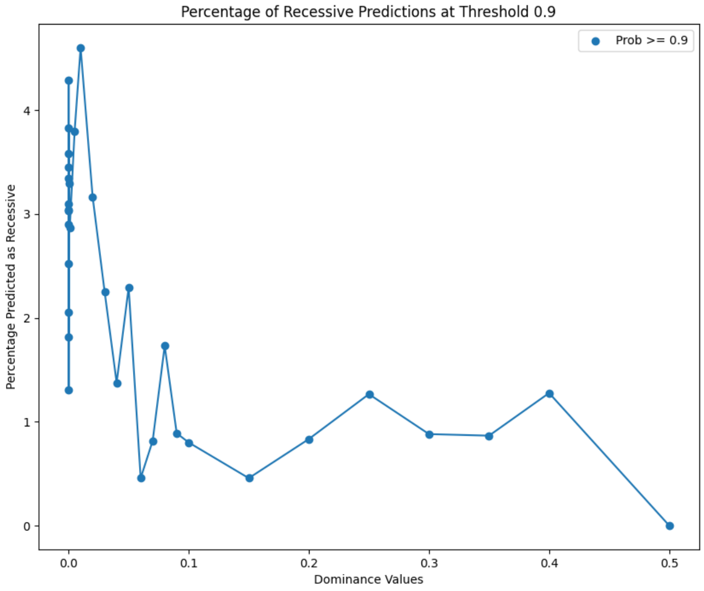


**e) LASSO**

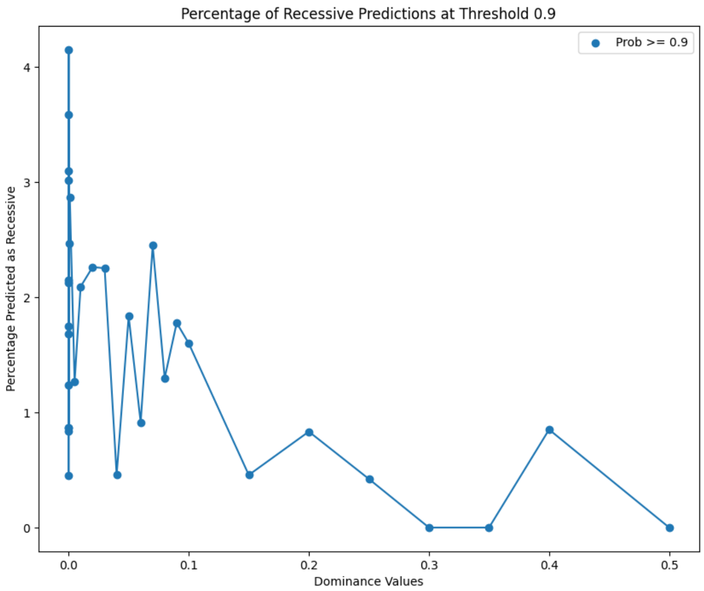


**f) Light GBM**

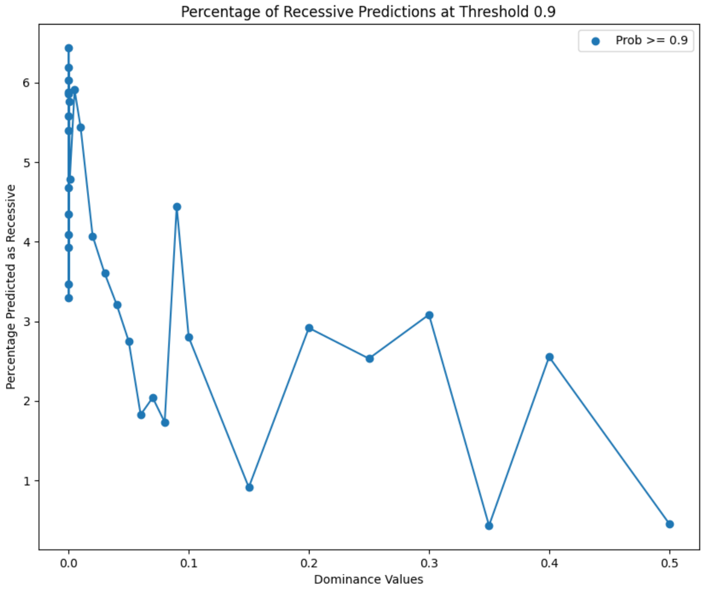


**g) Logistic regression**

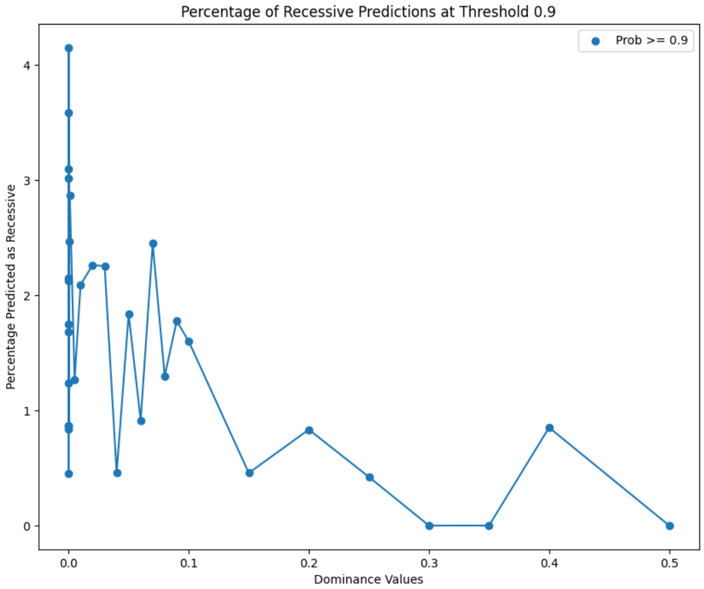


#### h) Elastic Net

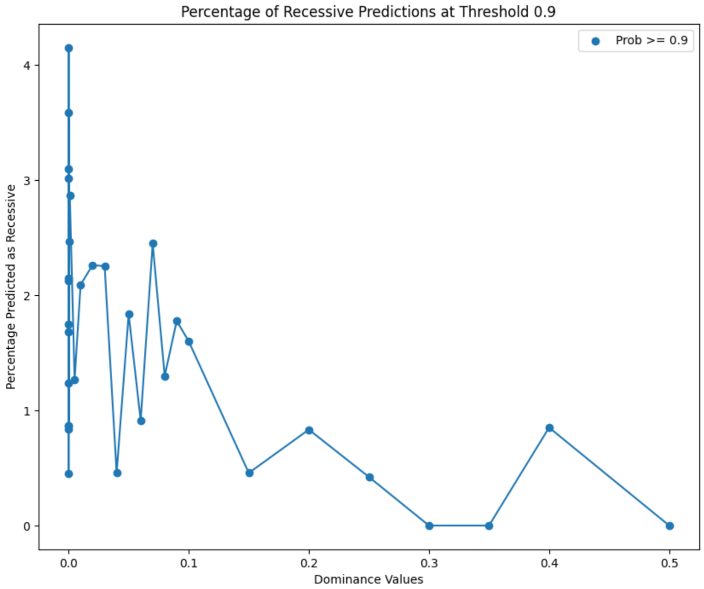


#### i) Random Forest

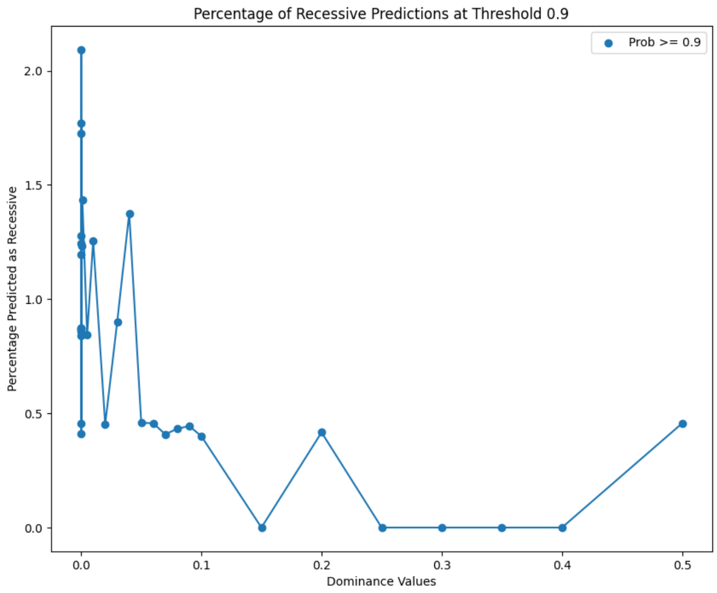


#### j) XGBoost before feature selection

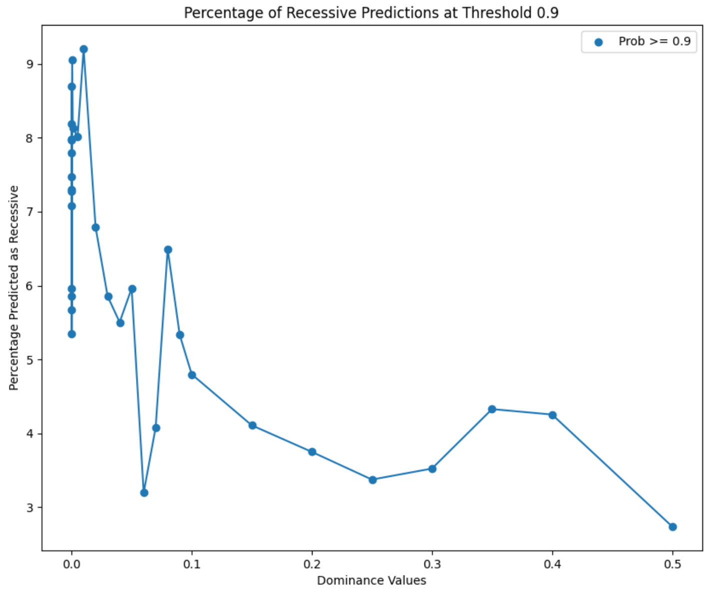


#### k) XGBoost after feature selection

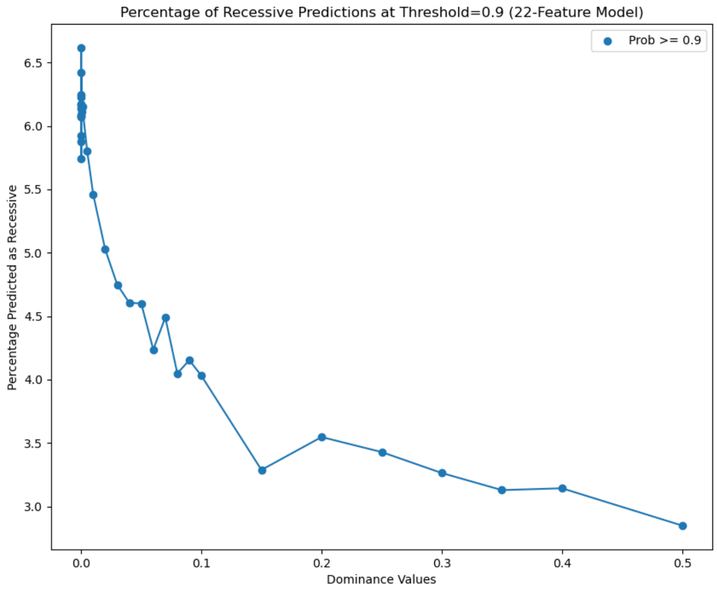


*We evaluate the performance of all ML classifiers considered by DominL by applying each trained classifier to new testing data that included simulations using non-extreme, intermediate dominance values (h between 0 and 1). For each h value category on the x-axis, we show the percentage of simulated windows being predicted as "Recessive" ($P_{recessive}$ > 0.9) on the y-axis. In general, all classifiers predicted substantially more recessive windows when h < 0.1, than for higher h values, showing that all ML classifiers considered can, in general, correctly distinguish highly recessive mutations from partially recessive mutations.*

**Supplementary Figure 7: XGBoost using 20 features performance comparison against using 22 features (20+2 recombination rate features)**

**a) All exon density, CEU**

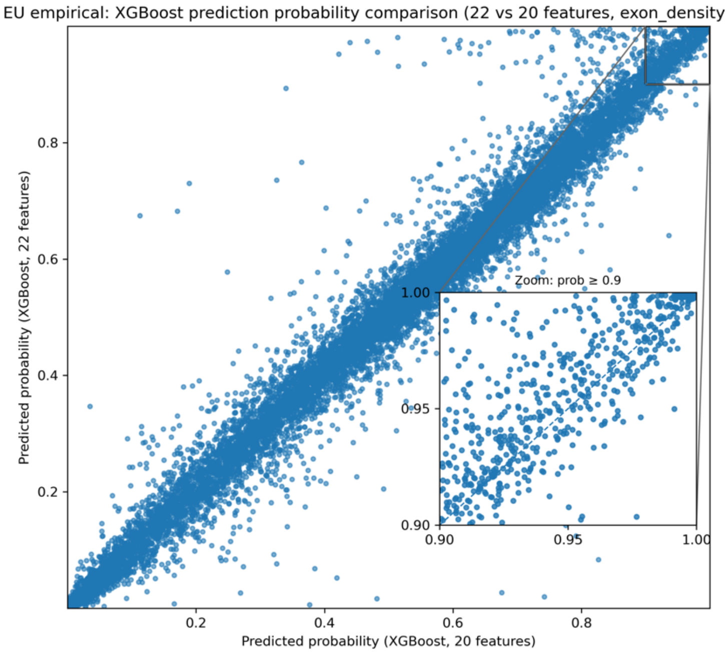


**b) Exon >=600, CEU**

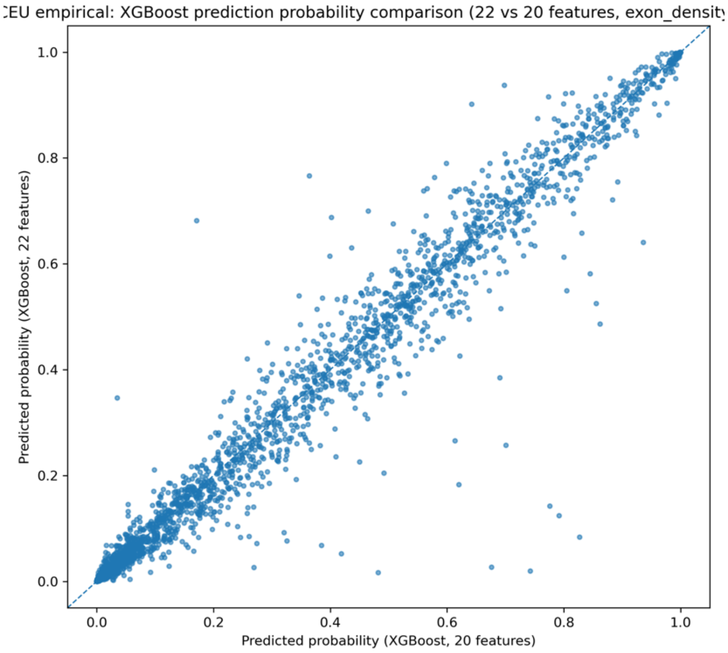


**c) All exon, CHB, show only Prediction Probability >= 0.7**

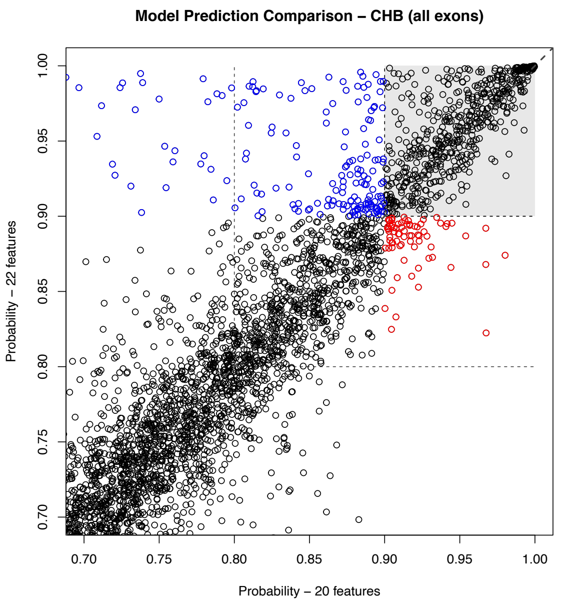


#### Supplementary Figure 8: DominL Robustness analyses, stratified by exon density

a) Misspecification of recombination rate (higher by magnitude of 10, orange, and lower by magnitude of 10, green), and with positive selection (red)

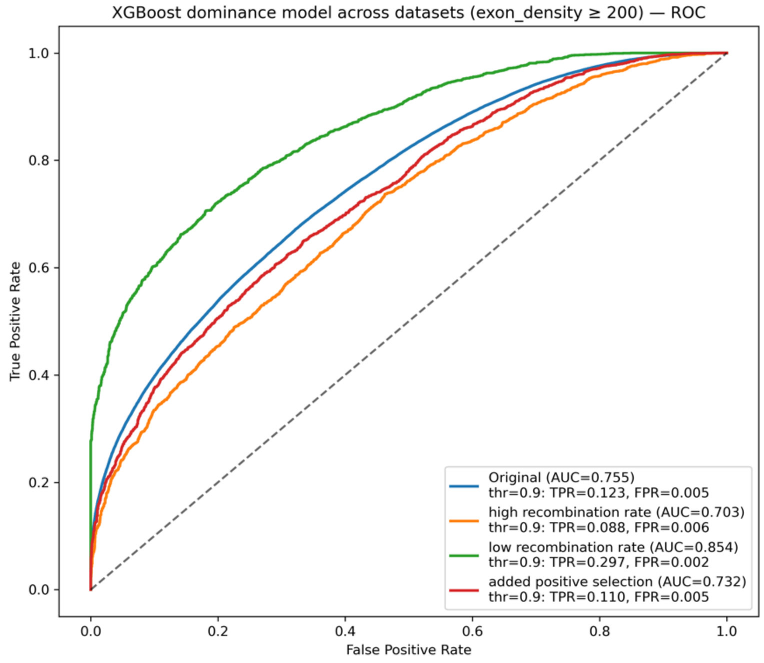

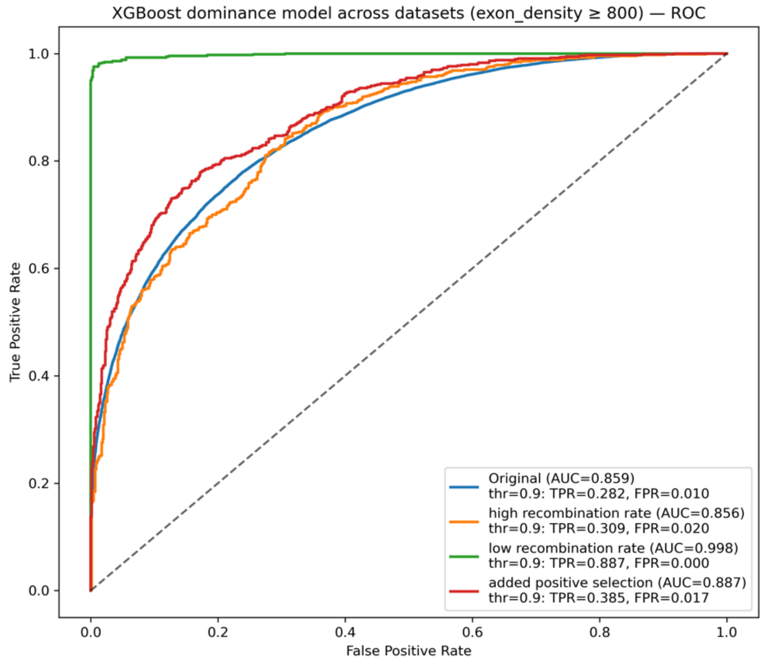
XGBoost dominance model across datasets (exon_density ≥ 800) — ROC
True Positive Rate
False Positive Rate
Original (AUC=0.859)
thr=0.9: TPR=0.282, FPR=0.010
high recombination rate (AUC=0.856)
thr=0.9: TPR=0.309, FPR=0.020
low recombination rate (AUC=0.998)
thr=0.9: TPR=0.887, FPR=0.000
added positive selection (AUC=0.887)
thr=0.9: TPR=0.385, FPR=0.017

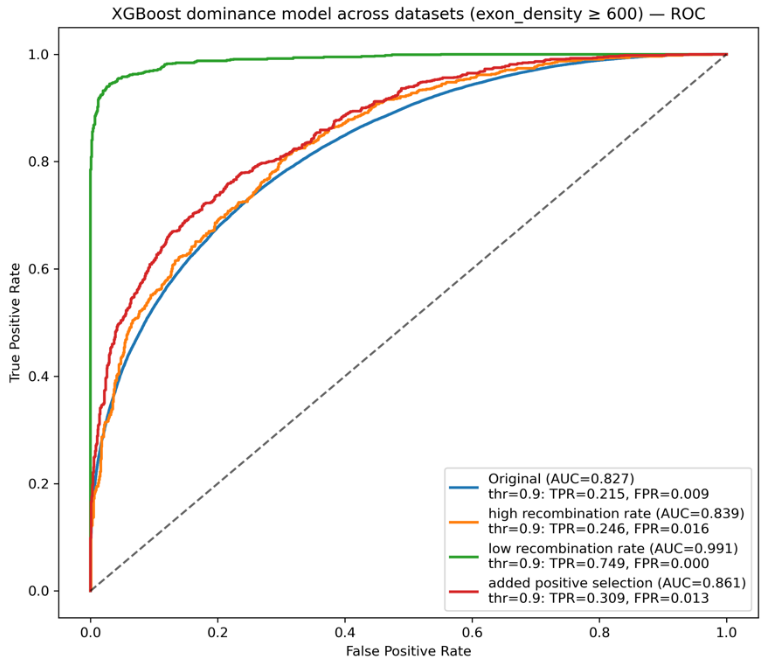
XGBoost dominance model across datasets (exon_density ≥ 600) — ROC
True Positive Rate
False Positive Rate
Original (AUC=0.827)
thr=0.9: TPR=0.215, FPR=0.009
high recombination rate (AUC=0.839)
thr=0.9: TPR=0.246, FPR=0.016
low recombination rate (AUC=0.991)
thr=0.9: TPR=0.749, FPR=0.000
added positive selection (AUC=0.861)
thr=0.9: TPR=0.309, FPR=0.013

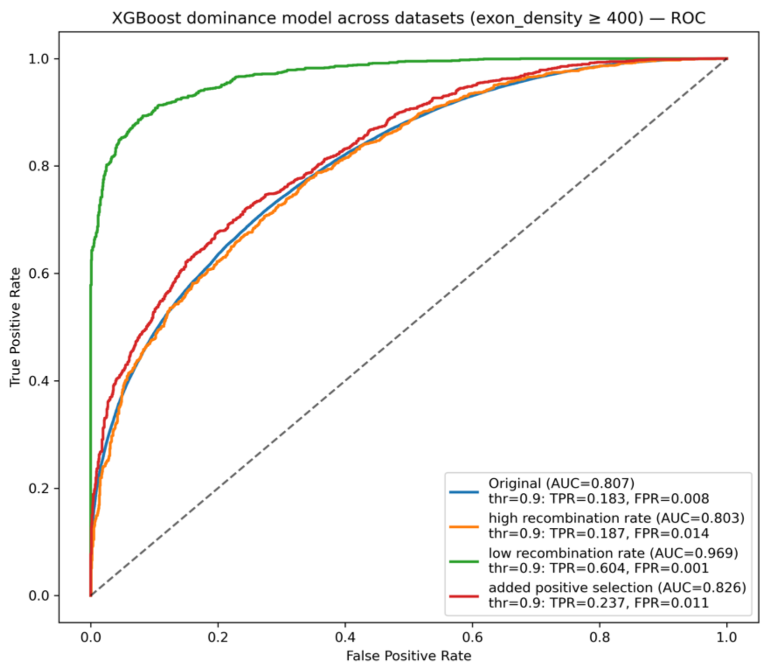


**b) Using CRF inferred Neanderthal ancestry (orange) instead of true ancestry (reported DominL, blue), stratified by exon densoty**

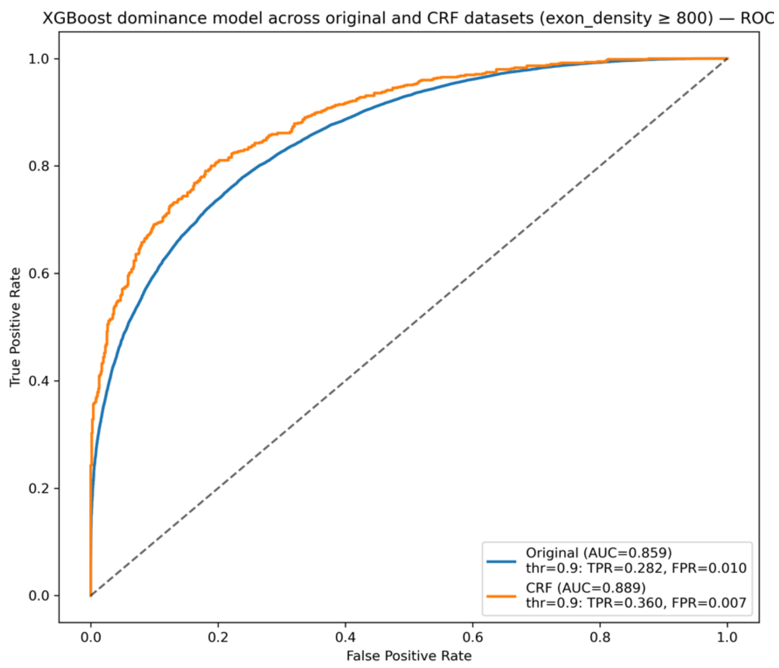
XGBoost dominance model across original and CRF datasets (exon_density ≥ 800) — ROC
True Positive Rate
False Positive Rate
Original (AUC=0.859)
thr=0.9: TPR=0.282, FPR=0.010
CRF (AUC=0.889)
thr=0.9: TPR=0.360, FPR=0.007

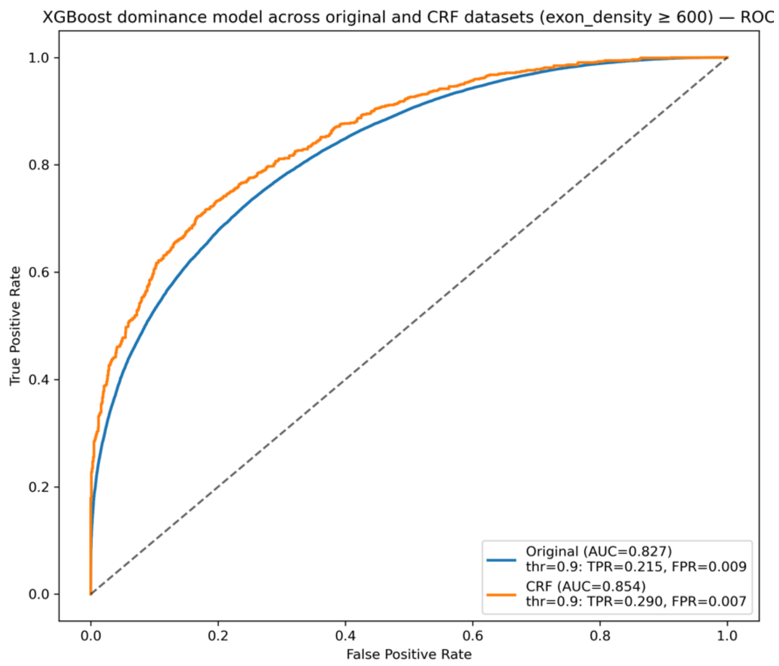
XGBoost dominance model across original and CRF datasets (exon_density ≥ 600) — ROC
True Positive Rate
False Positive Rate
Original (AUC=0.827)
thr=0.9: TPR=0.215, FPR=0.009
CRF (AUC=0.854)
thr=0.9: TPR=0.290, FPR=0.007

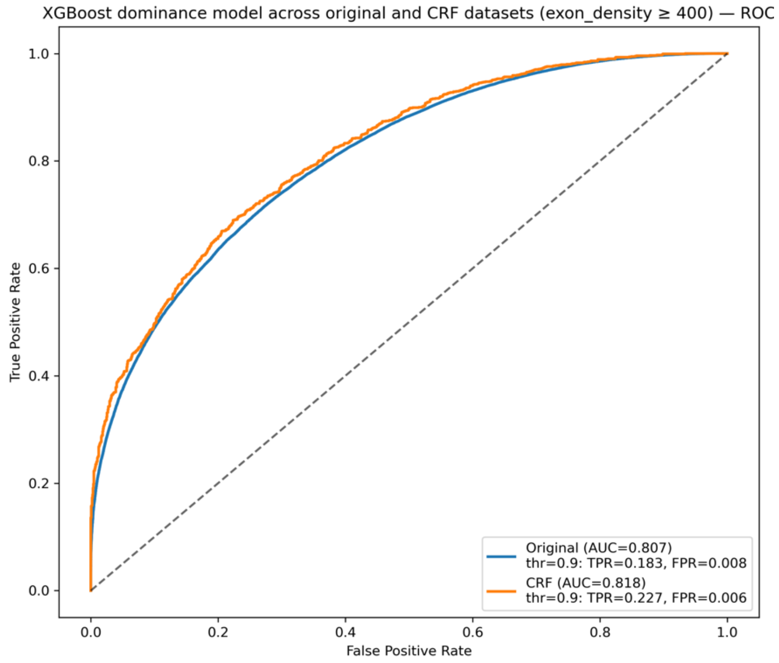


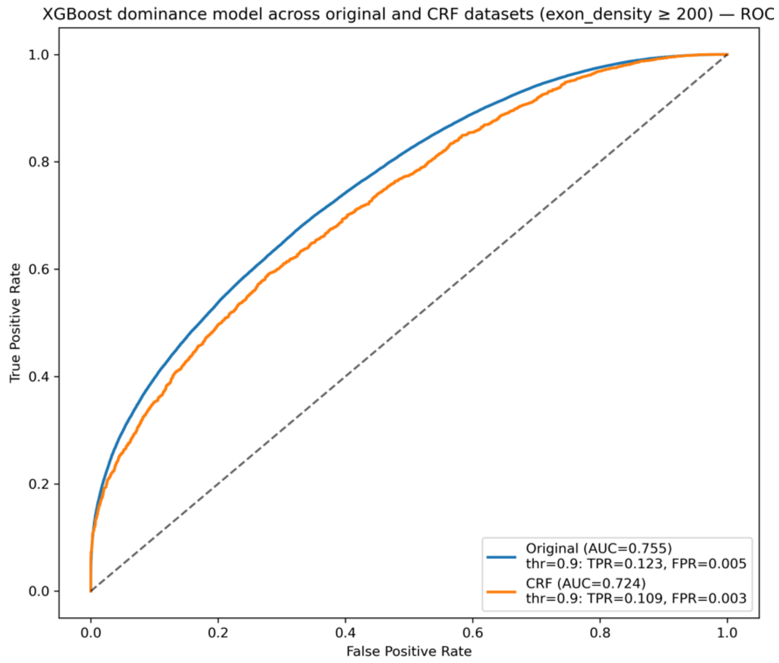


#### Supplementary Figure 9: Selected features ranked by feature importance used by XGBoost

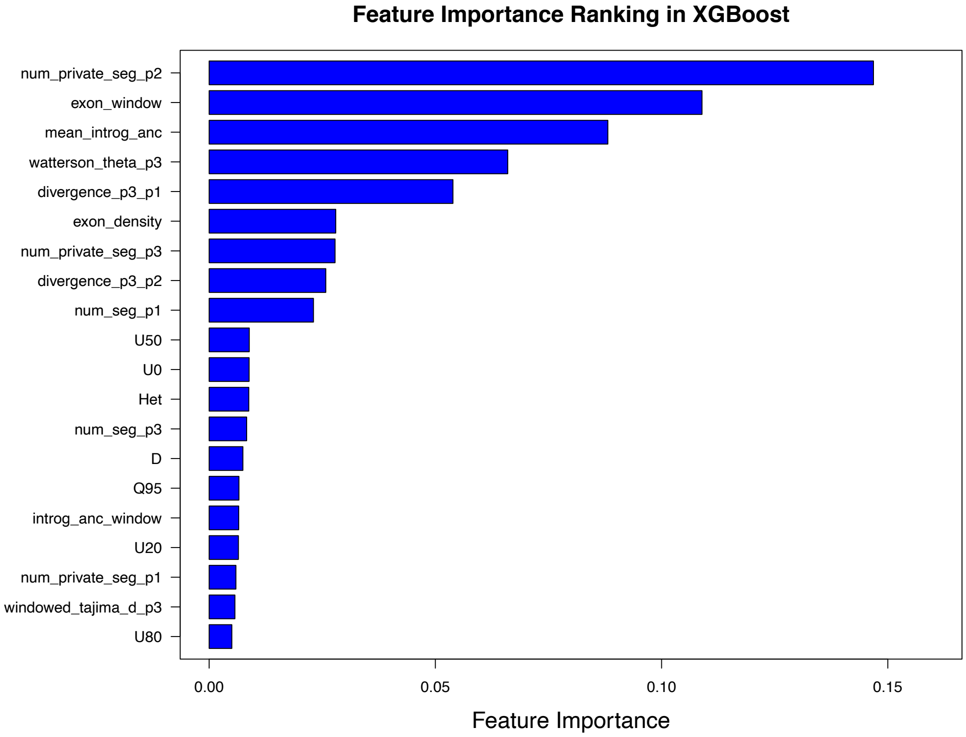


*We retrieved the feature importance scores (value between 0 to 1, all features sum up to 1) from DominL (XGBoost classifier) to understand its underlying deterministic model. We ranked and plotted the importance all features, and kept the top 20 features with both the highest feature importance score and the least discordance between simulation and empirical data, measured by the KL divergence.*

**Supplementary Figure 10: Distribution of Recessive Scores in all populations, using XGBoost after feature selection**

a) **Recessive predictions in the European population (CEU)**

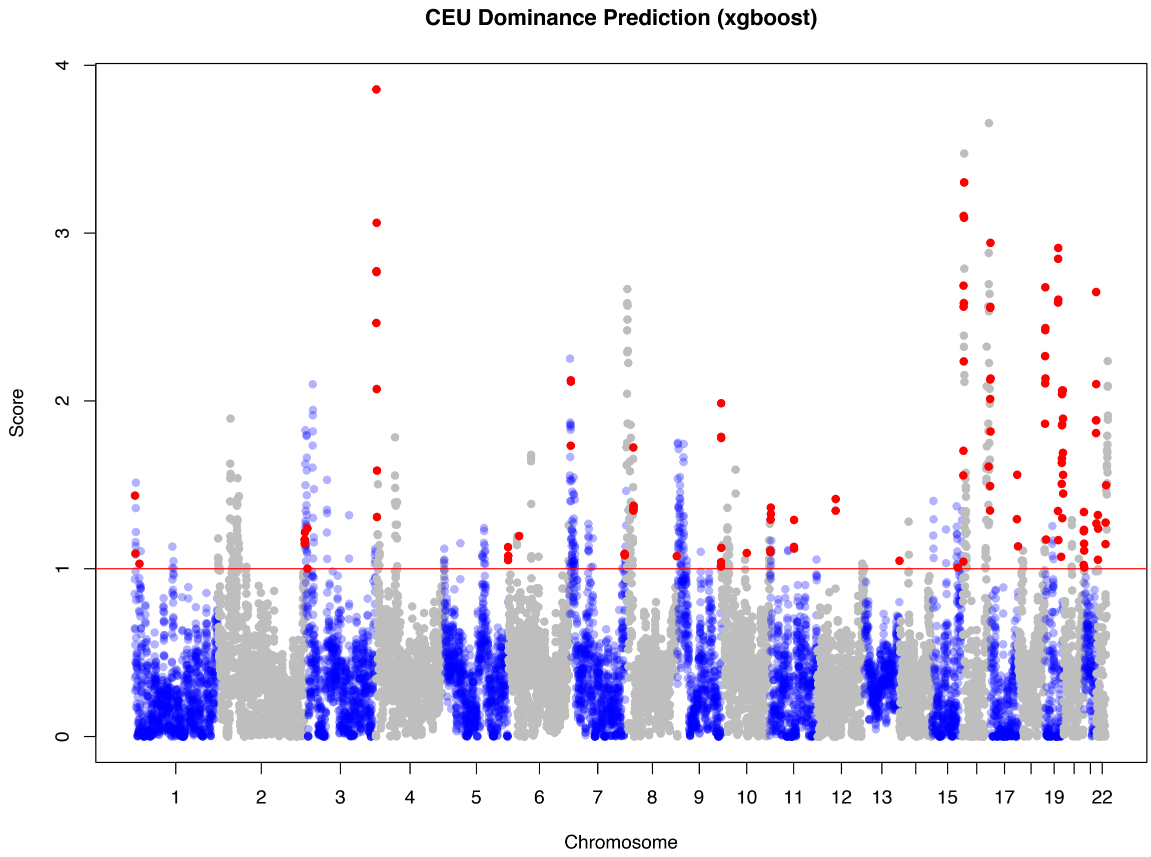


#### b) Recessive predictions in the Finnish population (FIN)

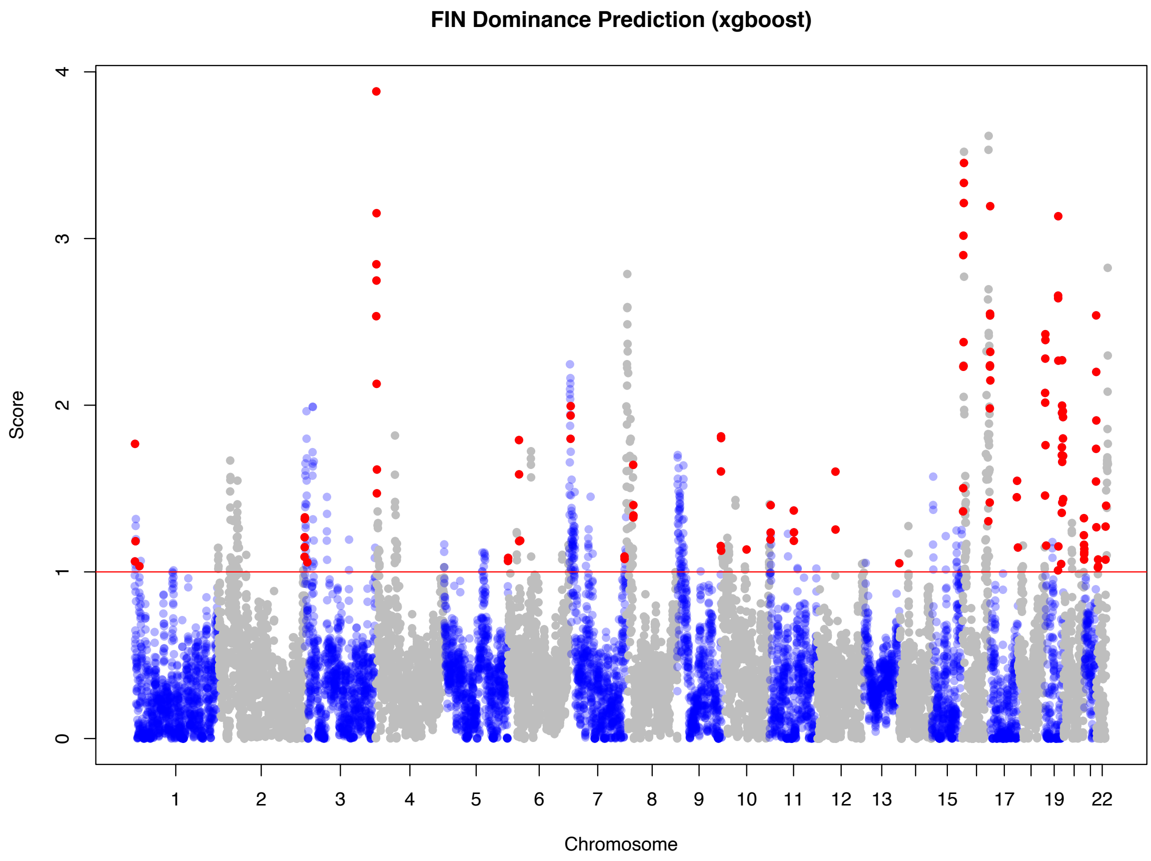


#### c) Recessive predictions in the Japanese population (JPT)

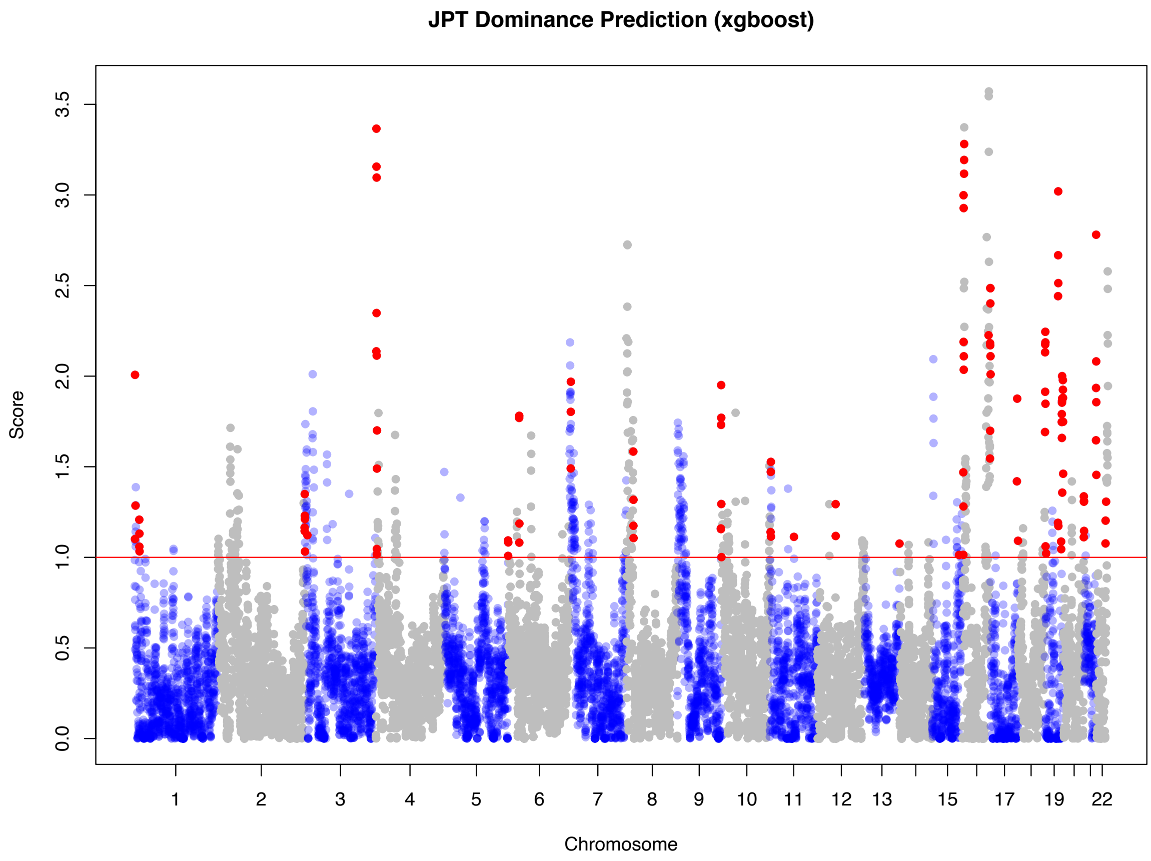


**d) Recessive predictions in the Southern Han Chinese population (CHS)**

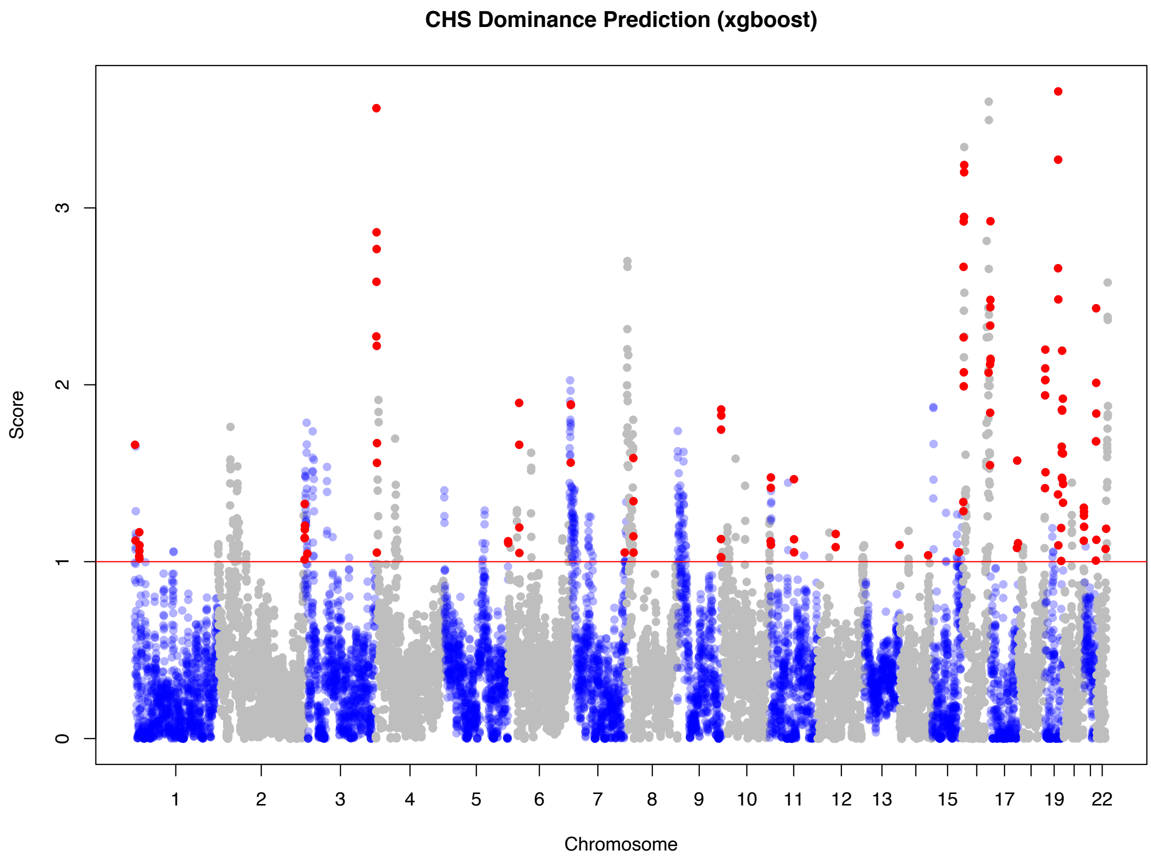


#### e) Recessive predictions in the Han Chinese population (CHB)

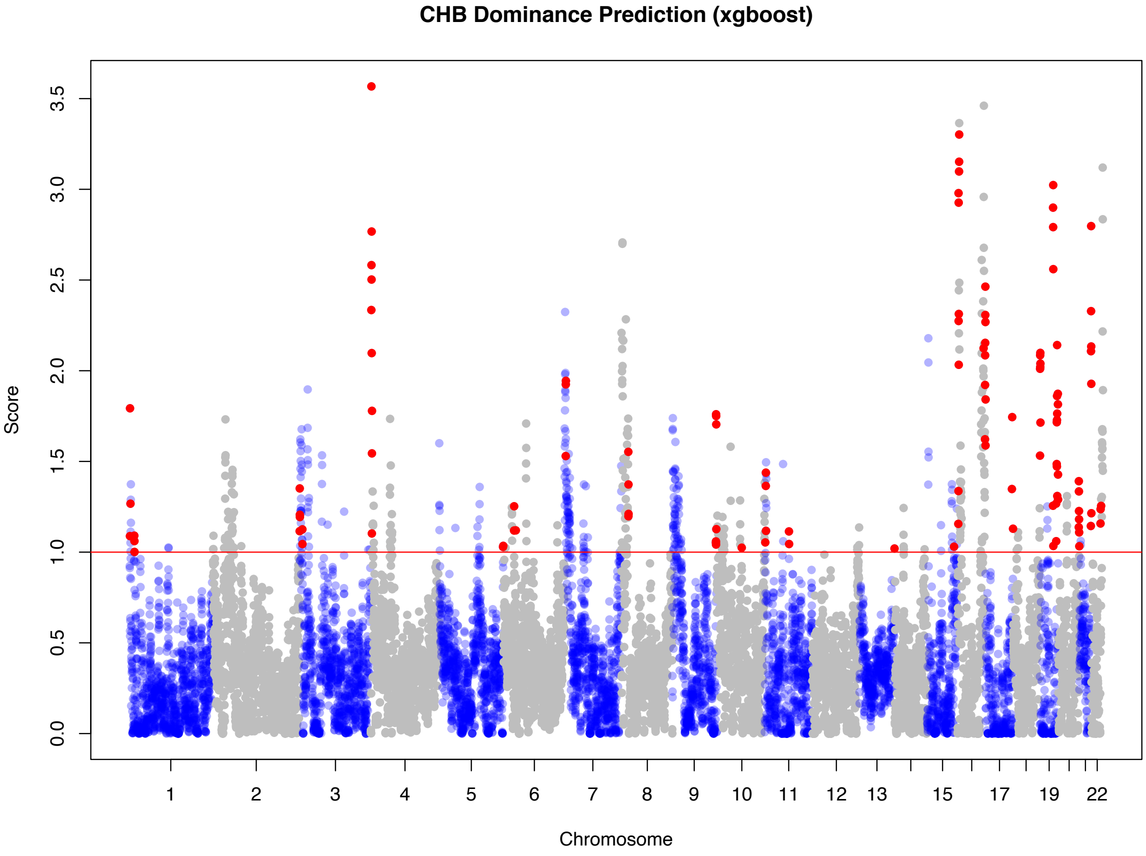


#### f) Recessive predictions in the Iberian population (IBS)

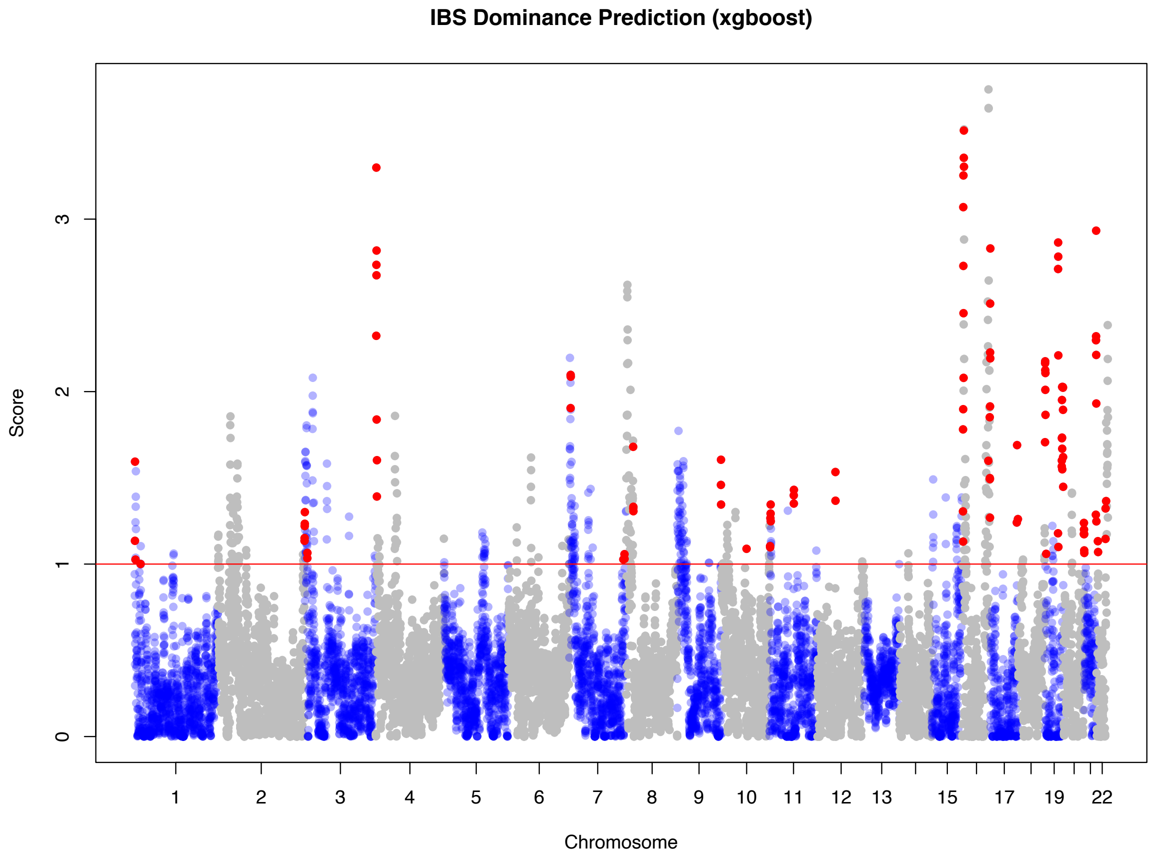


#### g) Recessive predictions in the British population (GBR)

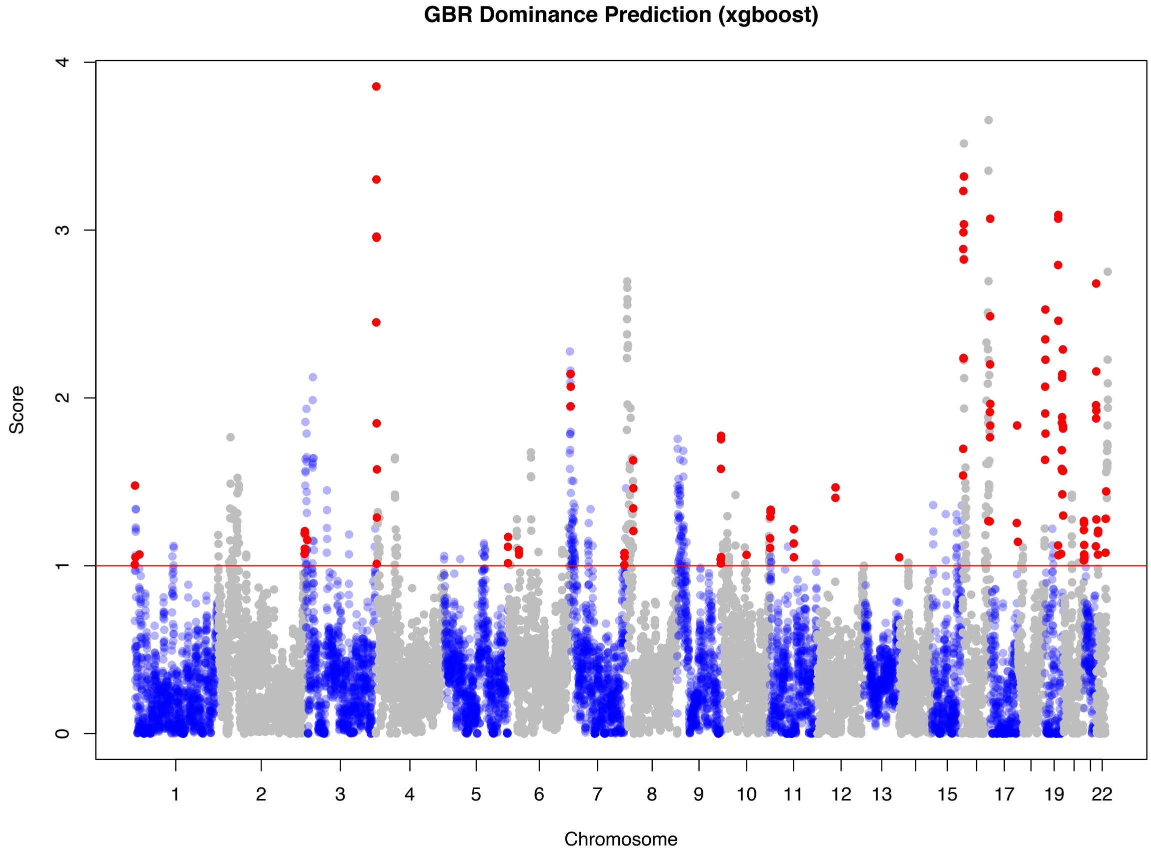


*From panel (**a**) through (**g**), we show the genome-wide distribution of regions predicted as recessive in 7 worldwide Eurasian populations included in the 1000 Genome Project, including CEU, FIN, CHS, CHB, JPT, IBS, and GBR. The significance threshold ($P_{recessive}$ = 0.9) is shown as red horizontal line, and regions with exon density >600/5MB are highlighted in red.*

**Supplementary Figure 11: Characteristics of recessive regions found exclusively in European or Asian populations**

**a) Recessive regions found only in European populations (at least one population among CEU, GBR, FIN, and IBS)**

**European–only recessive regions: total length = 22.2MB**

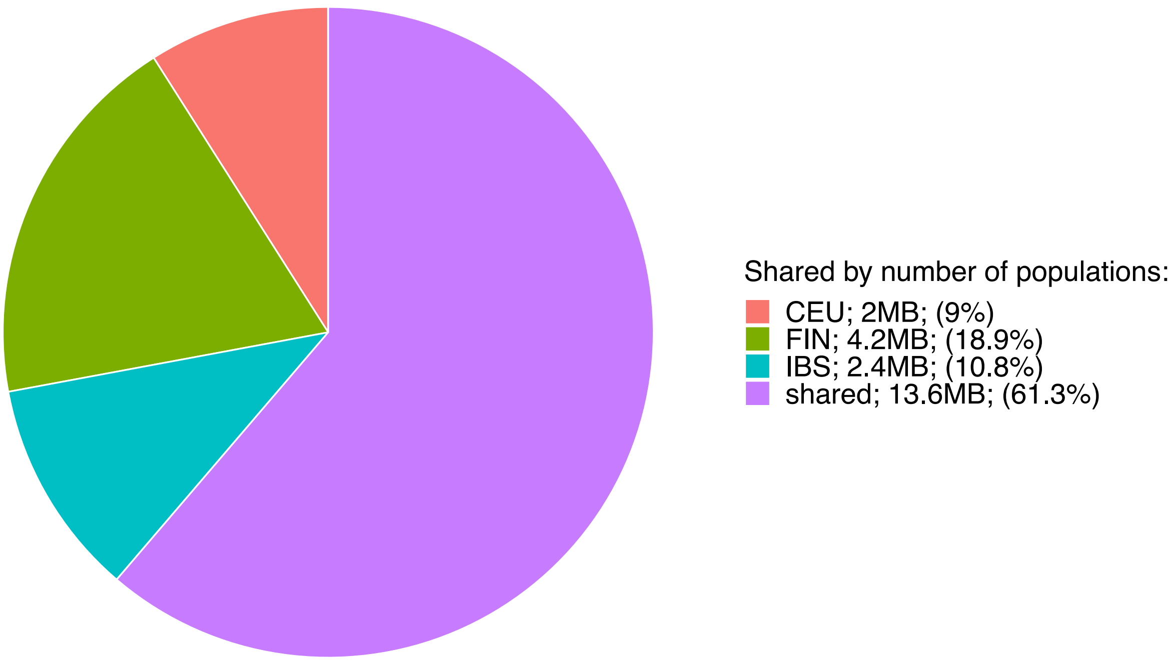


**b) Recessive regions found only in Asian populations (at least one population among CHB, CHS and JPT)**

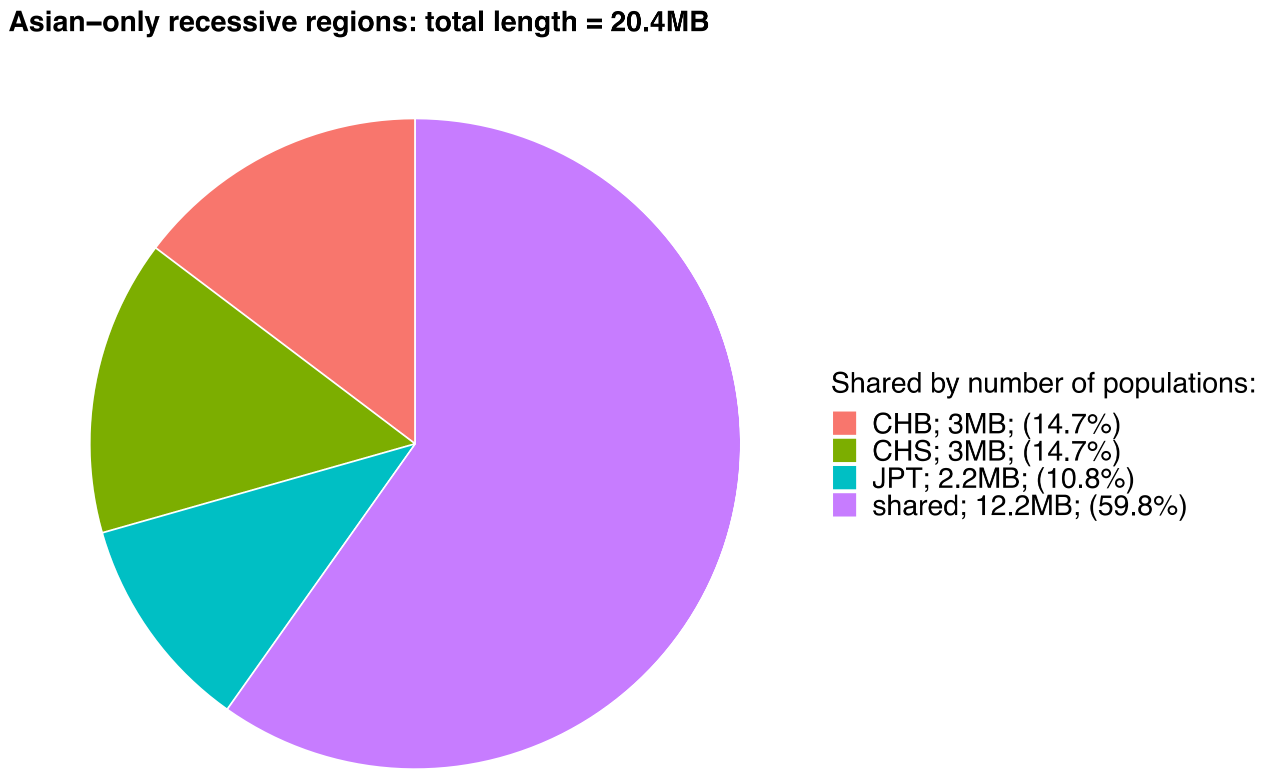


*Here we show the distribution of recessive regions found in (**a**) European populations only and (**b**) Asian populations only. There are approximately 20-22 MB total that are found in only one of the continents, among which most are shared by at least two populations on each continent. Among the regions that are found to be population-specific, Finnish population harbor the most population-specific recessive regions.*

**Supplementary Figure 12: Neanderthal ancestry distribution in different recessive score ranges**

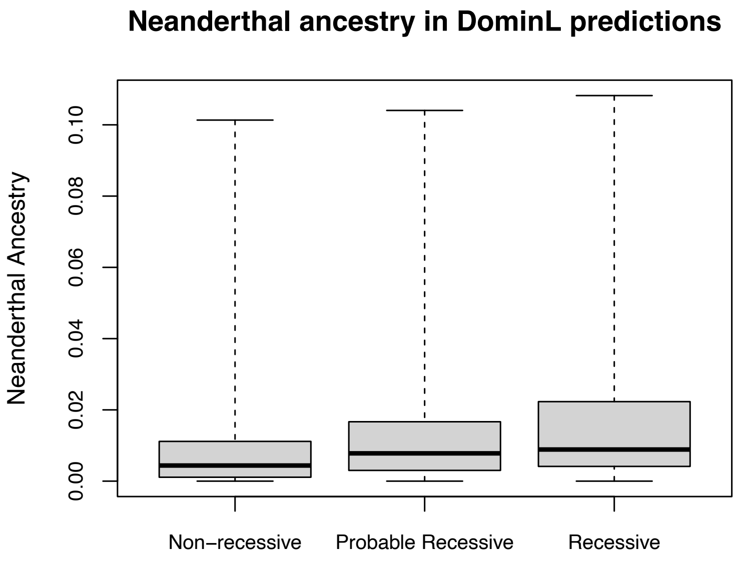


*We examine the validity of DominL prediction by plotting the distribution of Neanderthal ancestry in the 3 types of predictions in DominL, including Recessive ($P_{recessive}$ >=0.9), Probable Recessive ($P_{recessive}$ > 0.5 and <0.9), and Non-recessive ($P_{recessive}$ <=0.5). We show that the more DominL is confident at making a recessive call, the more Neanderthal ancestry there is in such region, consistent with what we expect from the underlying mechanism of DominL prediction, which relies heavily on the heterosis effect.*

**Supplementary Figure 13: Exon density distribution in different recessive score ranges**

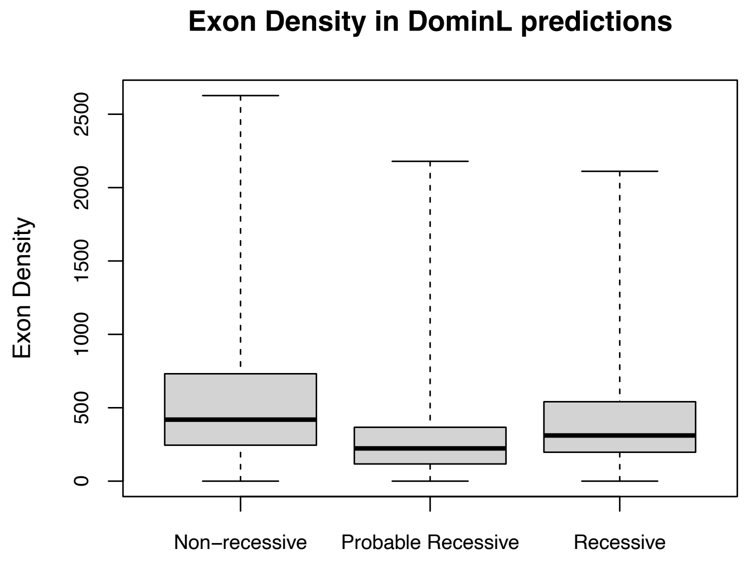


*Here we examine the distribution of exon density in the 3 types of predictions in DominL, including Recessive ($P_{recessive}$ >=0.9), Probable Recessive ($P_{recessive}$ > 0.5 and <0.9), and Non-recessive ($P_{recessive}$ <=0.5). We show that DominL predictions have little correlation with exon density, indicating DominL is not biased toward calling necessarily more recessive predictions in exon-dense regions.*

**Supplementary Figure 14: Feature distribution between recessive windows and all windows in empirical and training data**

a) **Pairwise feature comparisons in empirical data (CEU), with non-recessive predictions shown in black and recessive predictions shown in red**

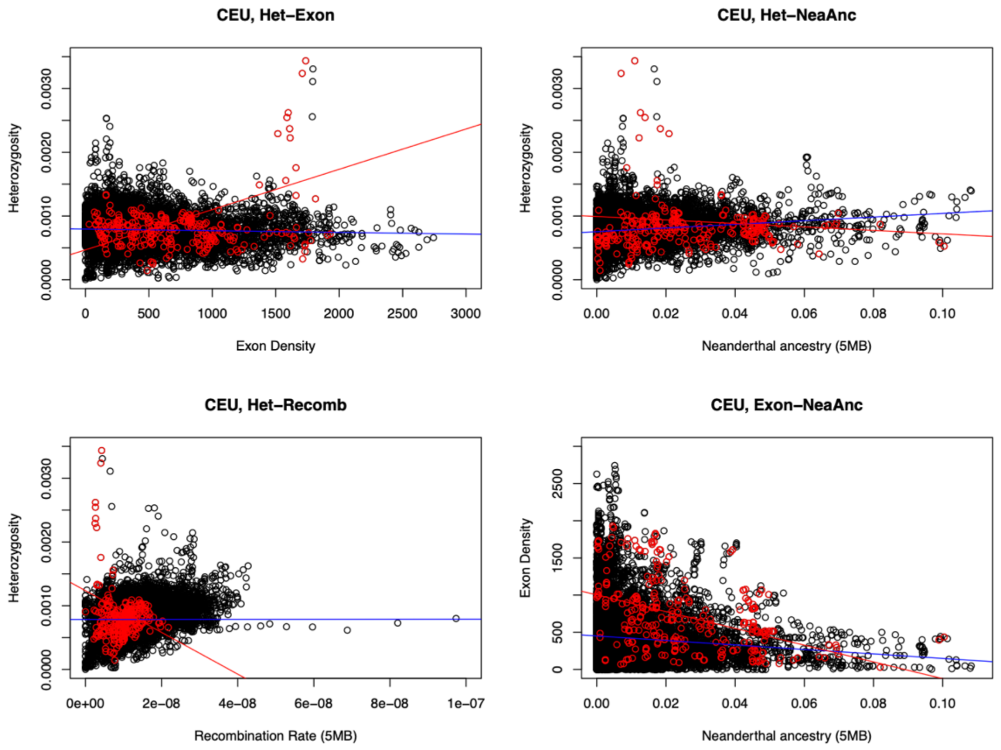


**b) Pairwise feature comparisons in the simulated training data, with non-recessive predictions shown in black and recessive predictions shown in red**

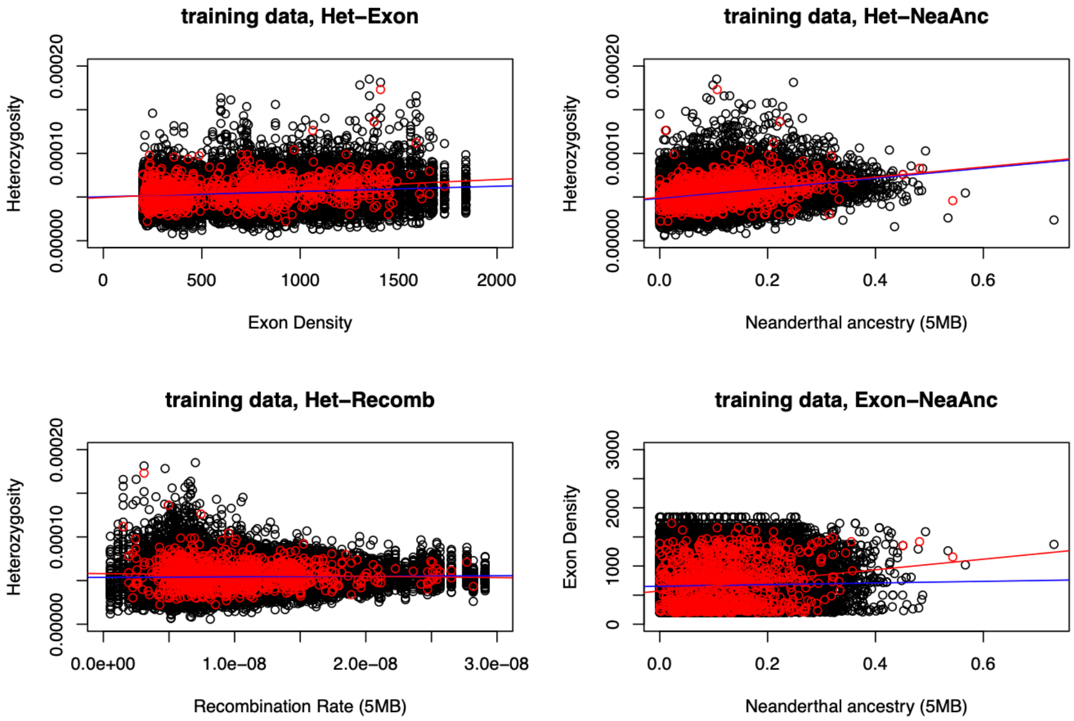


*We use pairwise distributions of 4 key features in (a) empirical data and (b) training data (simulations) to validate DominL by showing that the predicted recessive regions exhibit*

*biologically plausible relationships between features that are broadly consistent with simulation expectations. In each plot, the red line shows the linear regression of recessive predictions, and the blue line shows the linear regression of non-recessive predictions. The only pairwise feature comparison that shows a strong discrepancy between simulations and empirical data is the relationship between exon density and Neanderthal ancestry (the 4th panel). This observation can be readily explained by the fact that in simulations, we simulated fully recessive variants that lead to heterosis effects. Therefore, regions with more exons accumulated more fully recessive variants, which leads to more pronounced heterosis effects with increased Neanderthal ancestry. While in the empirical data, we generally observe the strong purifying selection against the Neanderthal ancestry, as additive deleterious effect likely accumulated throughout the human genome, even in regions with more recessive variants.*

**Supplementary Figure 15: The distribution of UKBB non-additive variants stratified by exon density in the "Non-recessive" category**

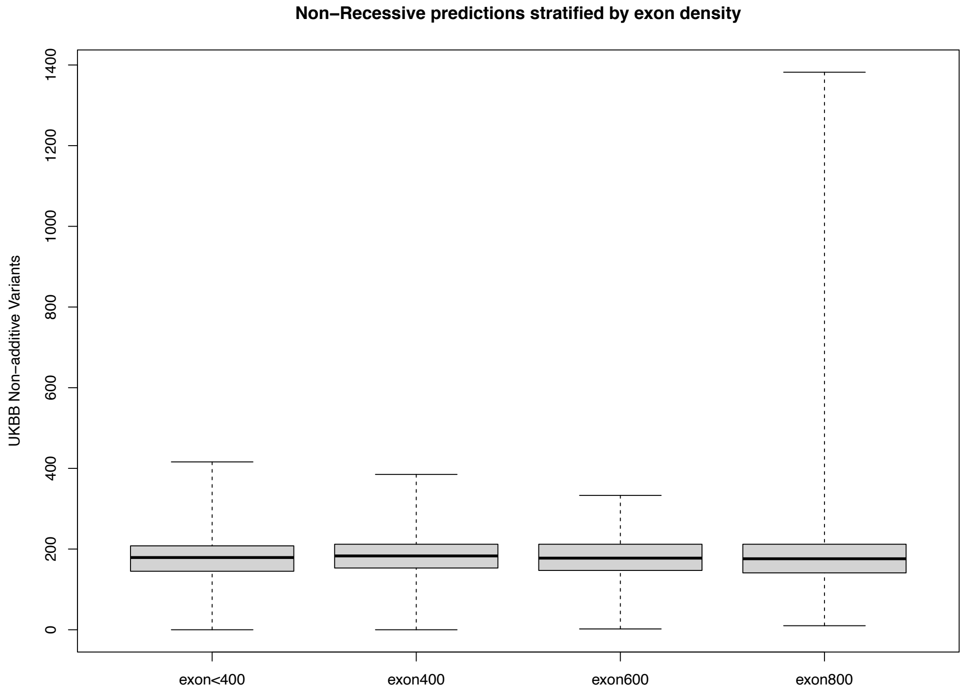


*We intersected the number of UKBB non-additive variants with the 1MB windows in our study that are classified as "Non-Recessive" ($P_{recessive}$ < 0.5), and stratified the distribution of such variants by the exon density in these windows. Unlike regions DominL confidently classified as "Recessive" shown in Fig. 4 ($P_{recessive}$ >= 0.9), in the "Non-Recessive" category, the number of non-additive UKBB variants shows a slight positive correlation with the exon density, with the highest exon density group having relatively the most UKBB variants.*

**Supplementary Figure 16: The distribution of UKBB non-additive variants stratified by exon density in the "Probable recessive" category**

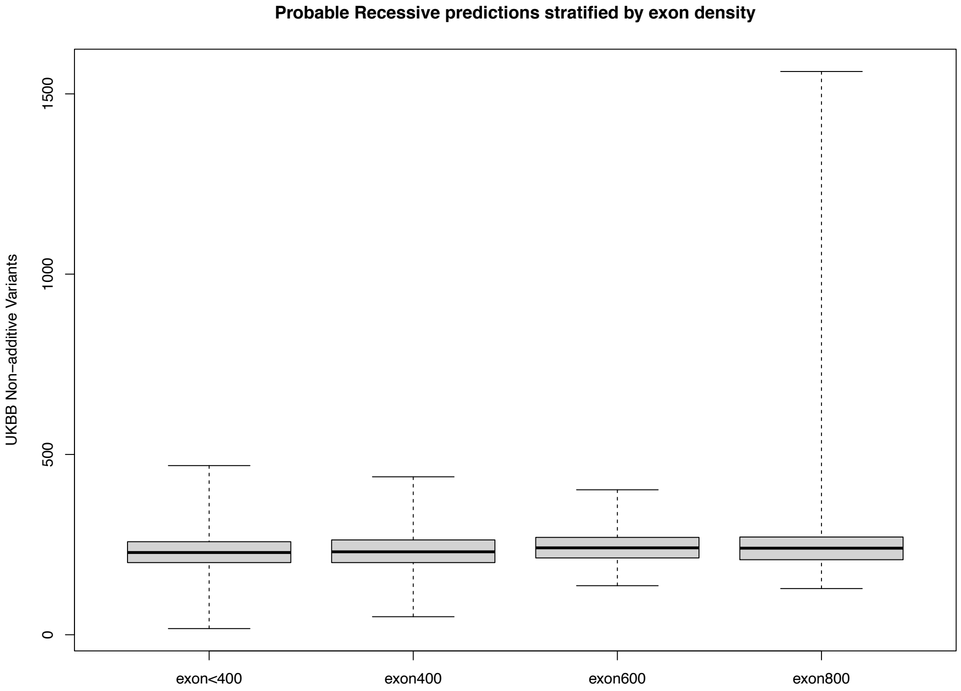


*We intersected the number of UKBB non-additive variants with the 1MB windows in our study that are classified as "Probable Recessive" ($P_{recessive}$ > 0.5 and <0.9), and stratified the distribution of such variants by the exon density in these windows. Unlike regions DominL confidently classified as "Recessive" shown in Fig. 4 ($P_{recessive}$ >= 0.9), in the "Probable Recessive" category, the number of non-additive UKBB variants shows a slight positive correlation with the exon density, with the highest exon density group having relatively the most UKBB variants.*

**Supplementary Figure 17: Exon Density Distribution on the genome**

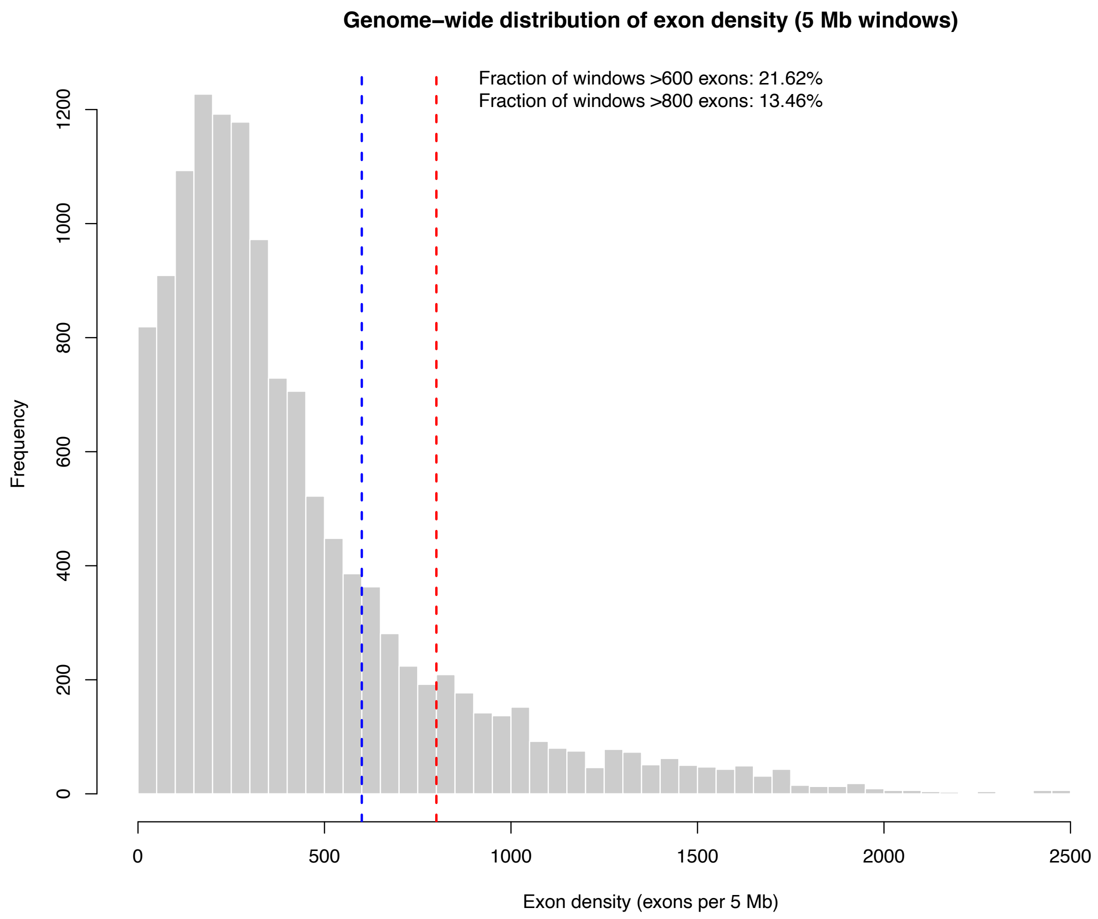


*We show the baseline genomic proportion of exon-dense regions to provide context for the observed enrichment. Specifically, exon-dense regions (>600 exons per 5 Mb) comprise 21.7% of the autosomes overall (blue dotted line), whereas 43.0% of recessive-predicted sequence falls within such regions, representing approximately a two-fold enrichment relative to the genomic baseline.*

**Supplementary Figure 18: Exon density in the test and control sets for genome biology analyses**

**a) Before matching**

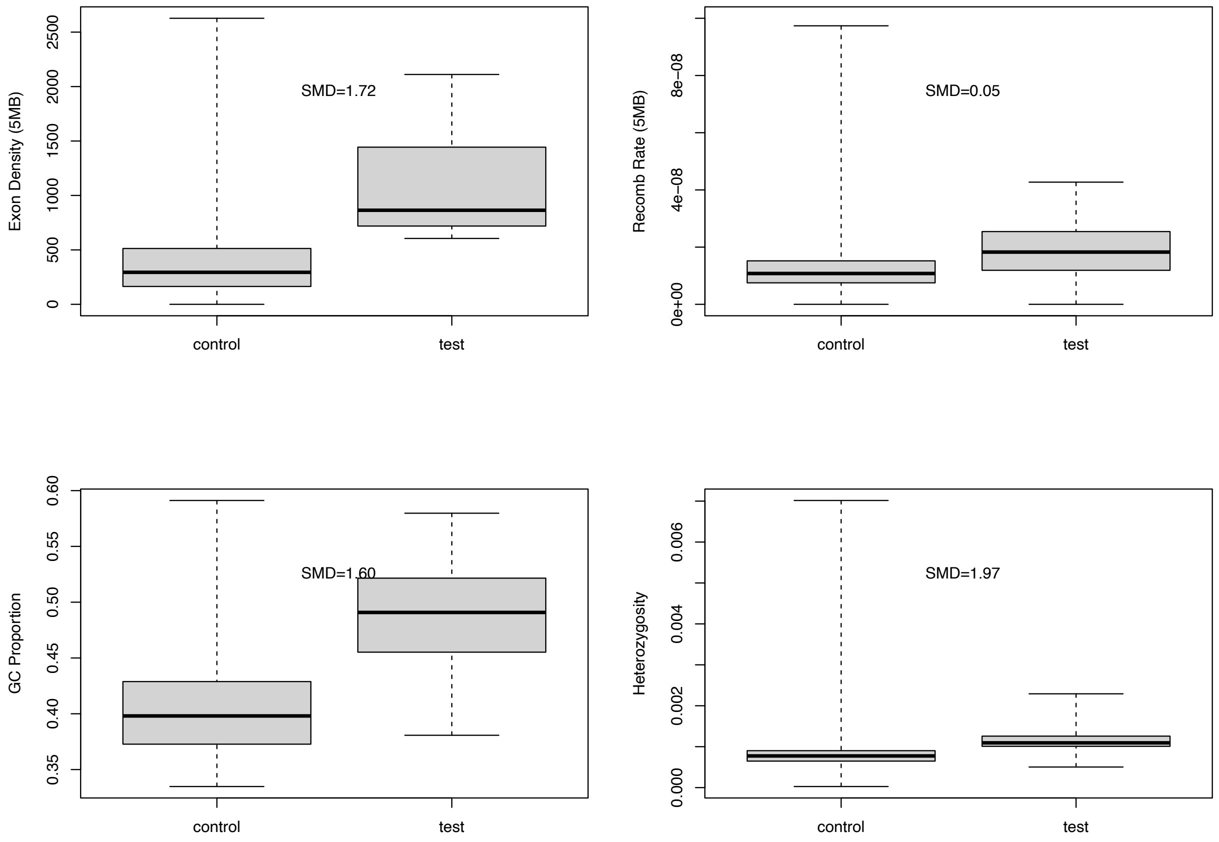


**b) Control for annotation analyses – matching exon density, recombination rate, and GC**

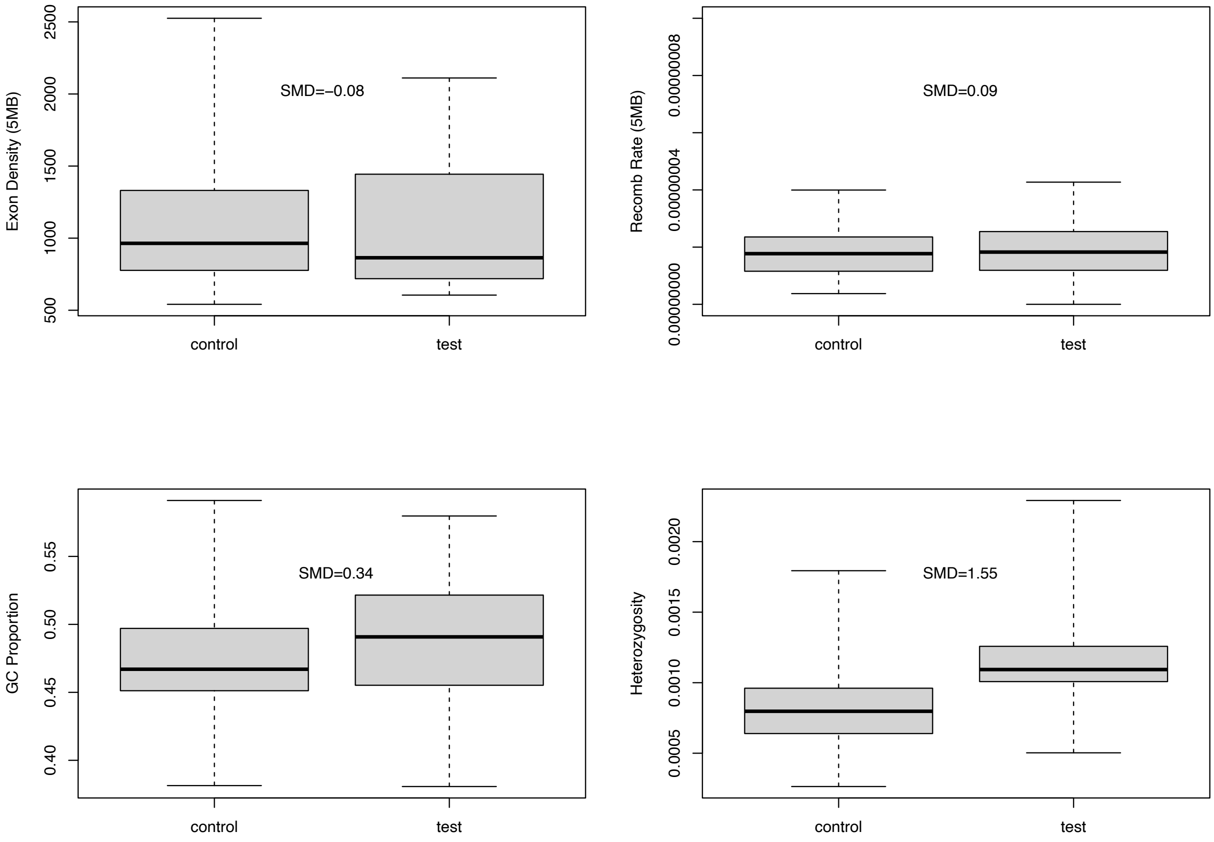


**c) Control for ROH analyses – matching exon density, recombination rate, and heterozygosity**

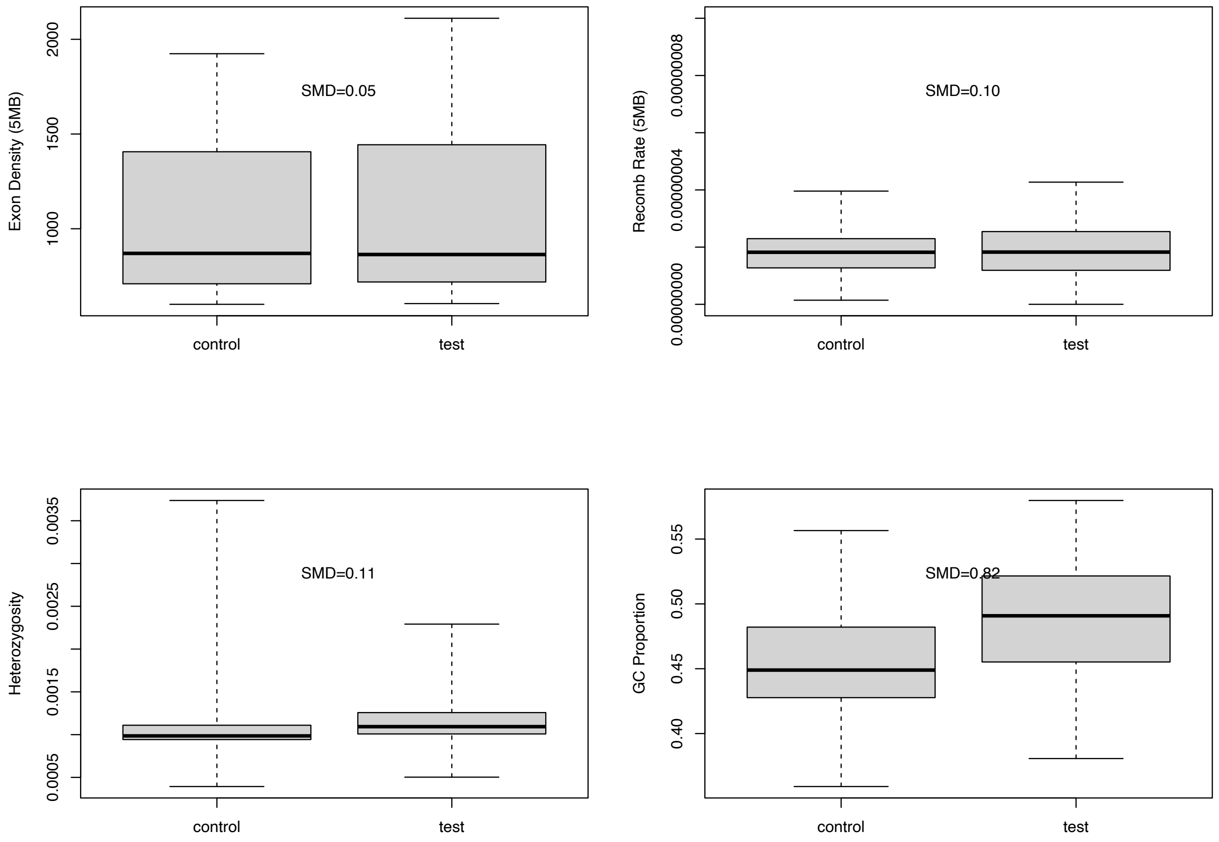


*We show in panel (a) the distribution of exon density, recombination rate, GC content, and Heterozygosity in recessive (test) and non-recessive regions (control) in terms of standardized mean differences SMD before matching on these parameters. Panel (b) shows the distribution of these parameters for the control set we used in annotation-based analyses where we match perfectly on exon density, recombination and GC. Panel (c) shows the distribution of these parameters for the control set we used in ROH analyses where perfect matching was achieved in exon density, recombination rate, and heterozygosity.*

**Supplementary Figure 19: ROH enrichment tests for in all 1KG populations**

a) **Intermediate, max length**

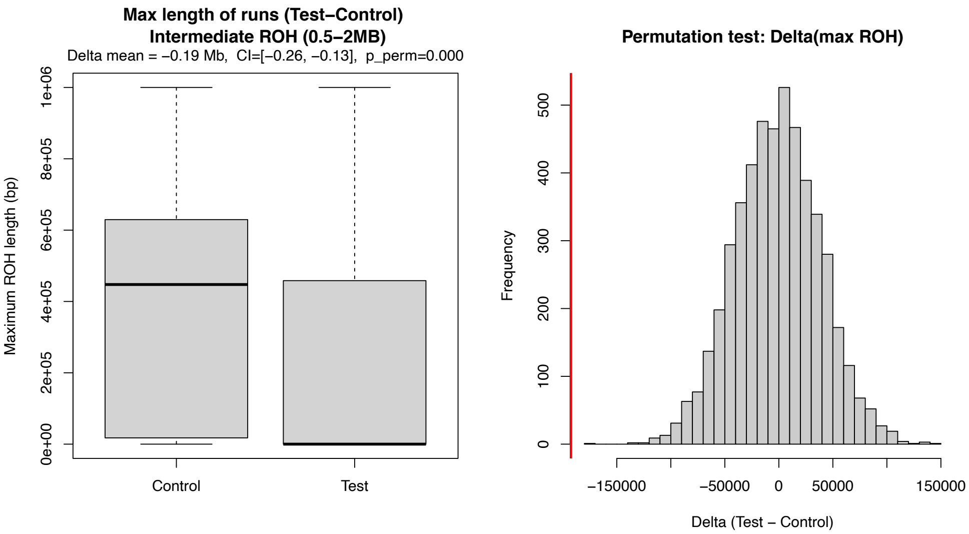


#### b) Intermediate, total length

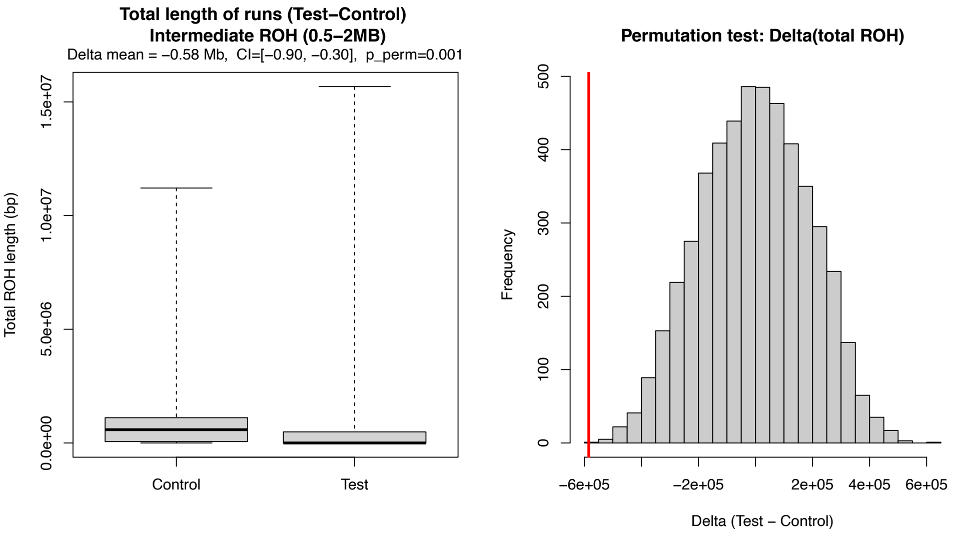


#### c) Intermediate, number of runs

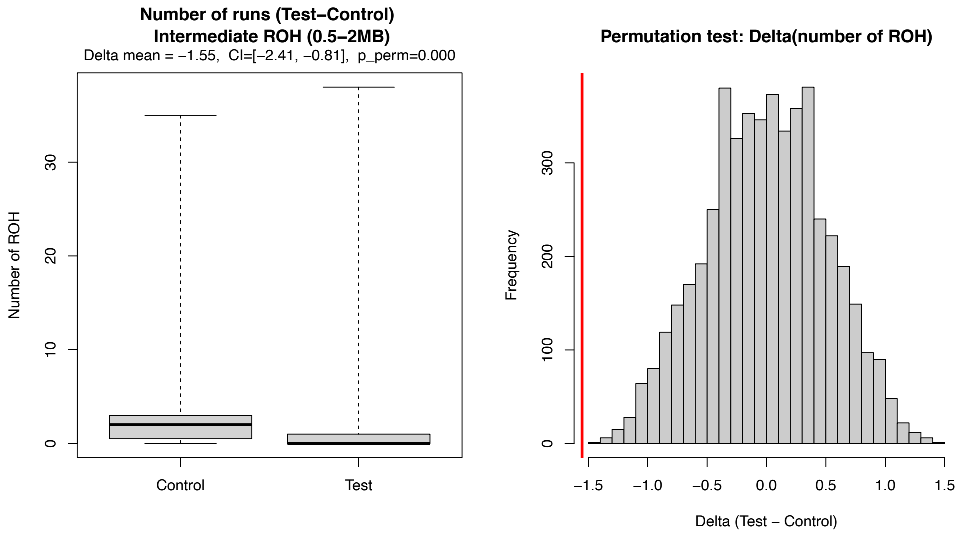


#### d) Long, max length

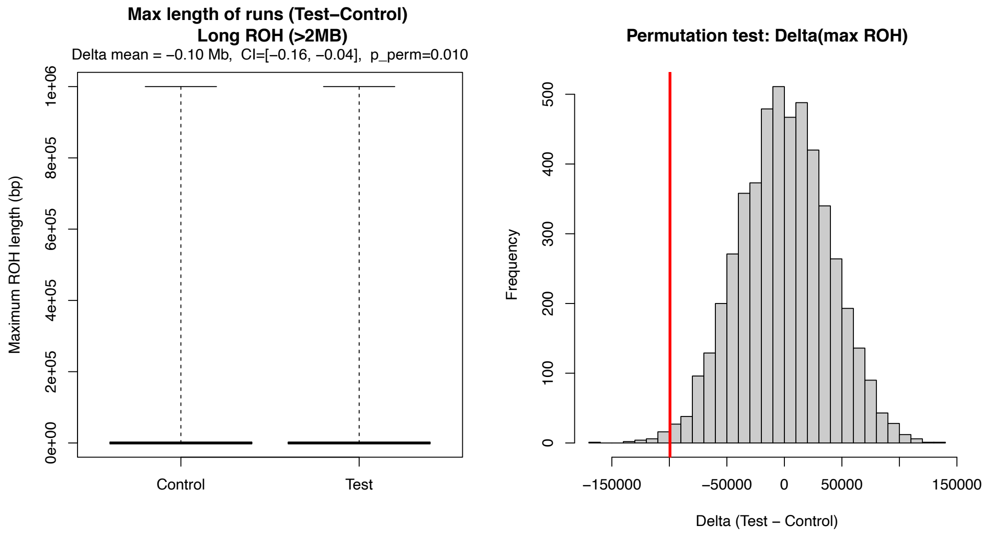


#### e) Long, total length

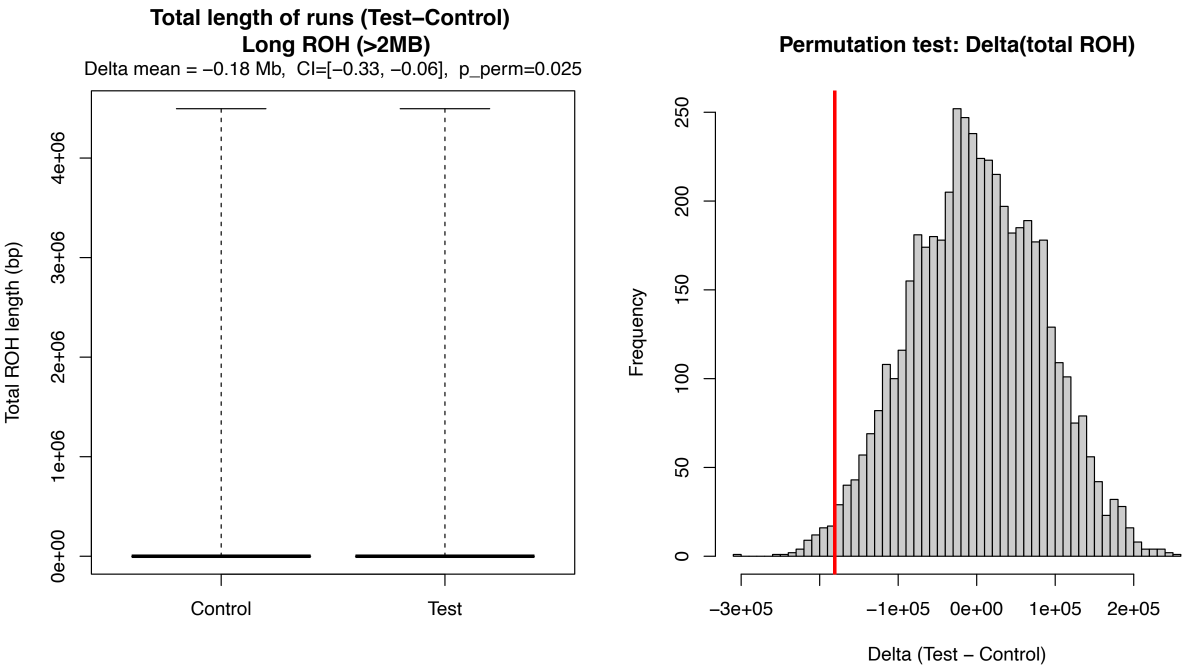


#### f) Long, number of runs

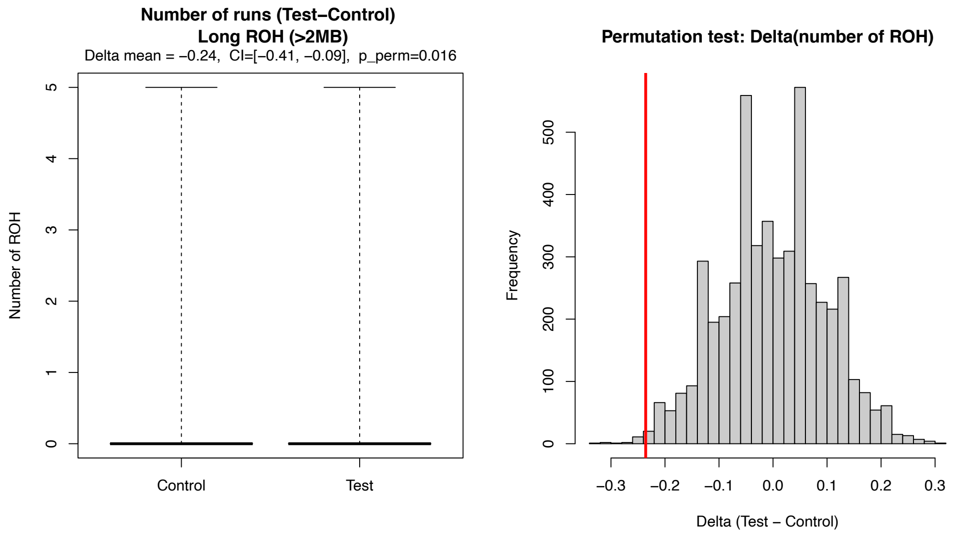


*We show in above panels the 1) maximum length of ROH, 2) total length of ROH, 3) number of ROH runs for intermediate length of ROHs (0.5-2MB, panel a-c) and for long ROHs (>2MB, panel d-f) that overlap with recessive windows and covariate-mathced control windows. Left side of the panel shows ROH distribution for the given metric between recessive (test) and control, and the right panel shows the permutation null distribution of the mean difference between test and control (test-control), with the observed value highlighted in red. Both intermediate and long ROHs are significantly depleted from recessive regions by all three tests,*

**Supplementary Figure 20: ROH enrichment tests for in all 1KG populations excluding HLA regions**

**a) Intermediate, max length, no HLA**

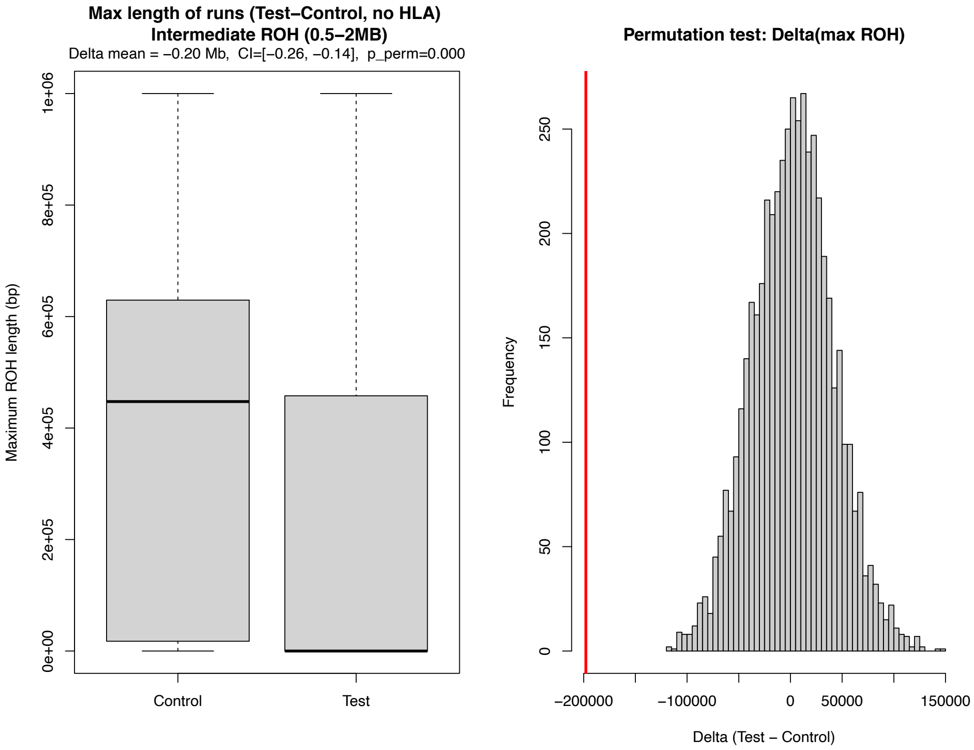


#### b) Intermediate, total length, no HLA

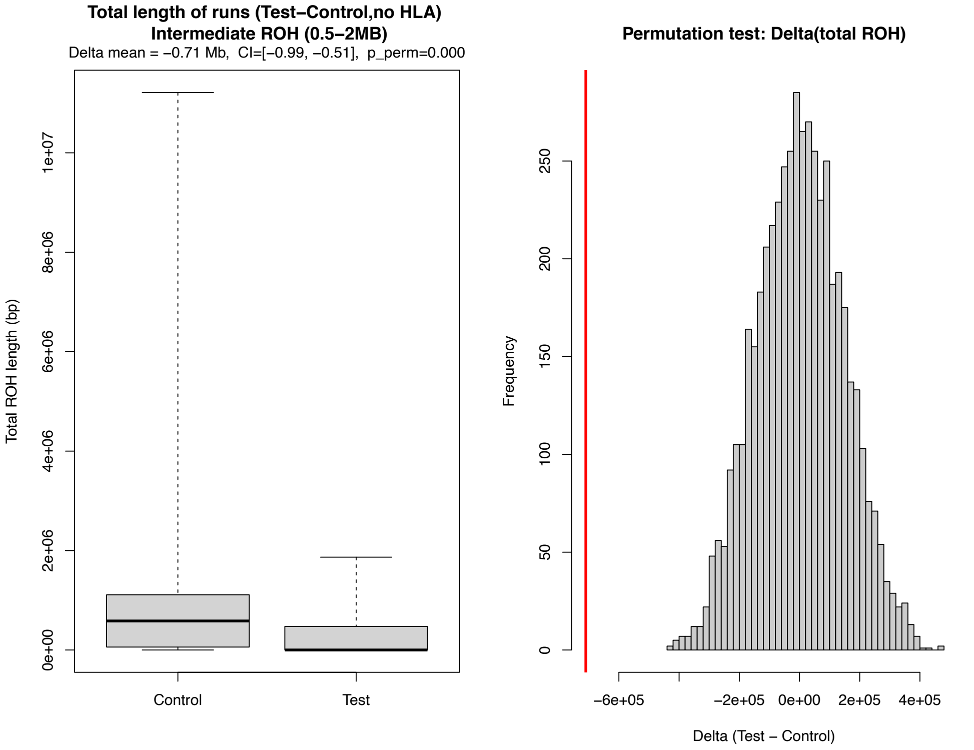


#### c) Intermediate, number of runs, no HLA

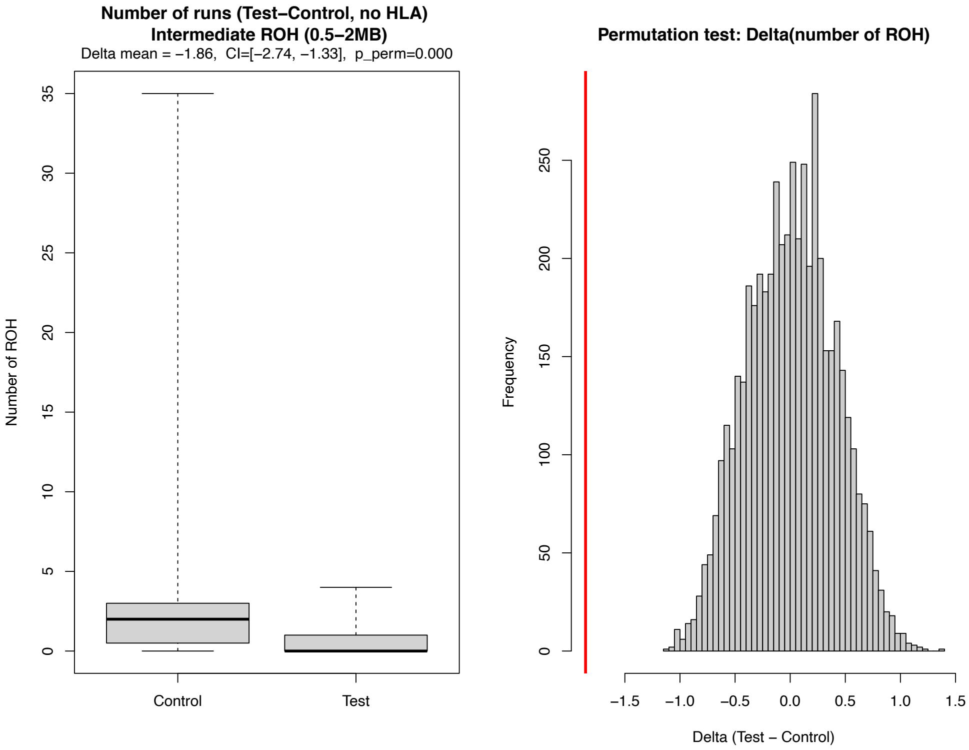


#### d) Long, max length, no HLA

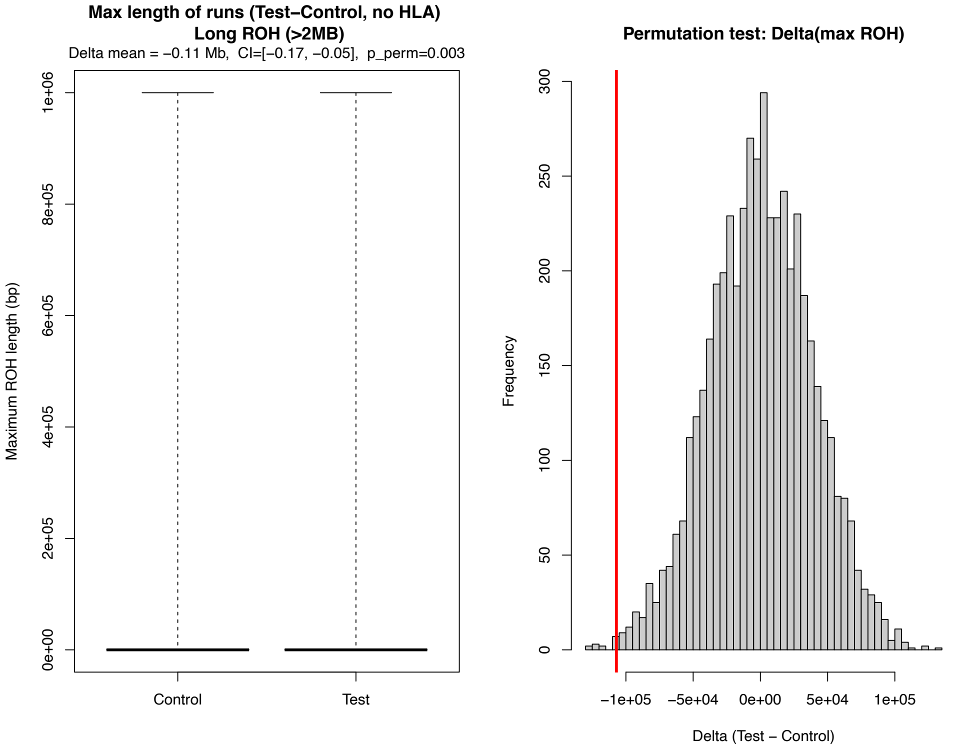


#### e) Long, total length, no HLA

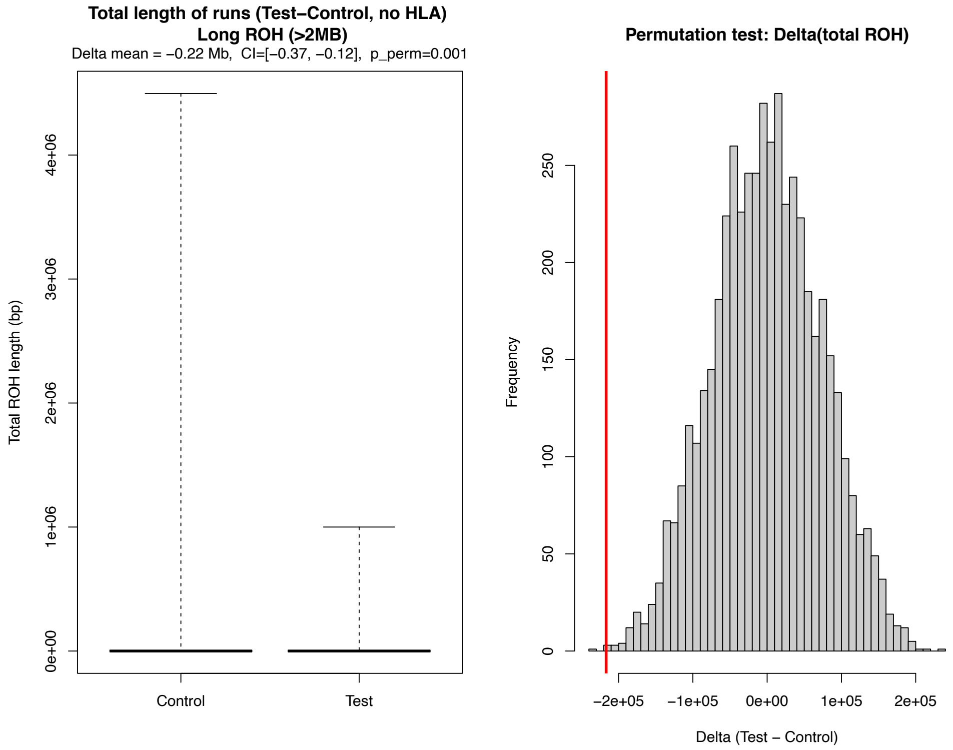


#### f) Long, number of runs, no HLA

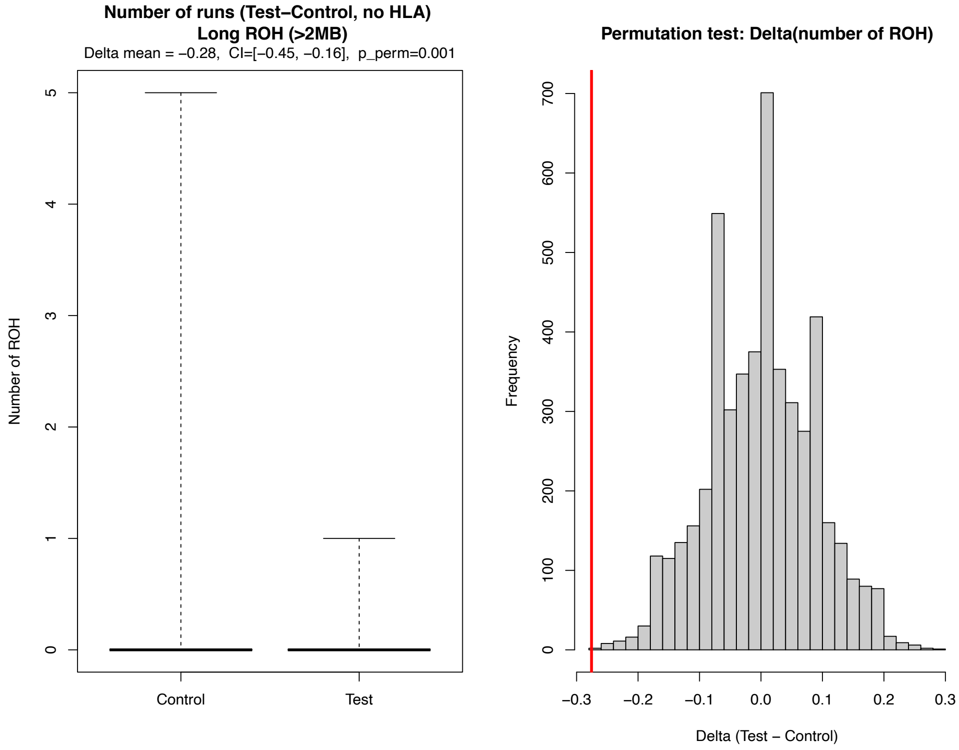


*We show in above panels the 1) maximum length of ROH, 2) total length of ROH, 3) number of ROH runs for intermediate length of ROHs (0.5-2MB, panel a-c) and for long ROHs (>2MB, panel d-f) that overlap with recessive windows and covariate-mathced control windows. Left side of the panel shows ROH distribution for the given metric between recessive (test) and*

*control, and the right panel shows the permutation null distribution of the mean difference between test and control (test-control), with the observed value highlighted in red. The main difference between this figure and Fig. S19 is that the HLA region on Chromosome 6 is removed from the test set. Both intermediate and long ROHs are significantly depleted from recessive regions by all three tests, and the depletion pattern is more significant than when the HLA region is included in the test set.*

**Supplementary Figure 21: McVicker's B value distribution in control regions, OMIM gene regions, and recessive regions**

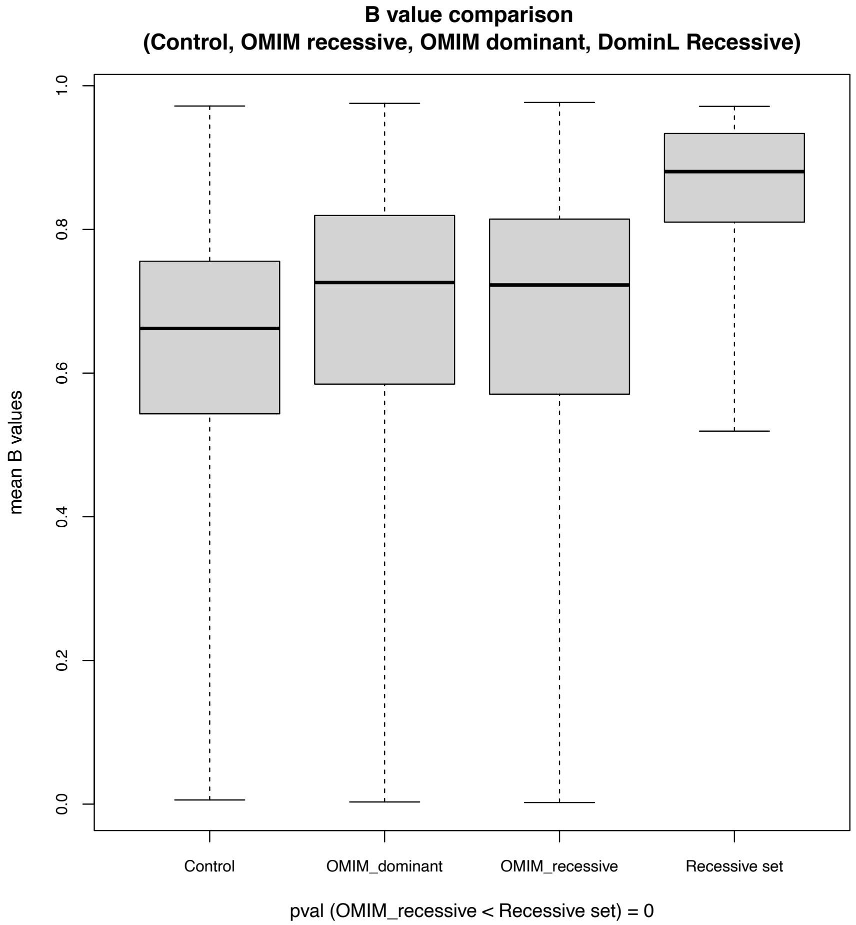


*The distribution of McVicker's B values in the following four groups: 1) control regions, defined as randomly sampled 1MB regions on the genome with total length and exon density distribution matching the DominL detected recessive regions; 2) autosomal dominant gene regions in OMIM database; 3) autosomal recessive gene regions in OMIM database; and 4) DominL detected recessive regions, defined as regions with exon density >=600 exons/5MB, and are shared by at least 6 human populations. The DominL detected recessive regions experienced the least historical background selection (highest B values), and both OMIM gene types experienced strong historical BGS.*

### Supplementary Figure 22: Chromosome-end localization analyses
#### a) Genome-wide genomic-feature maps

Exon density

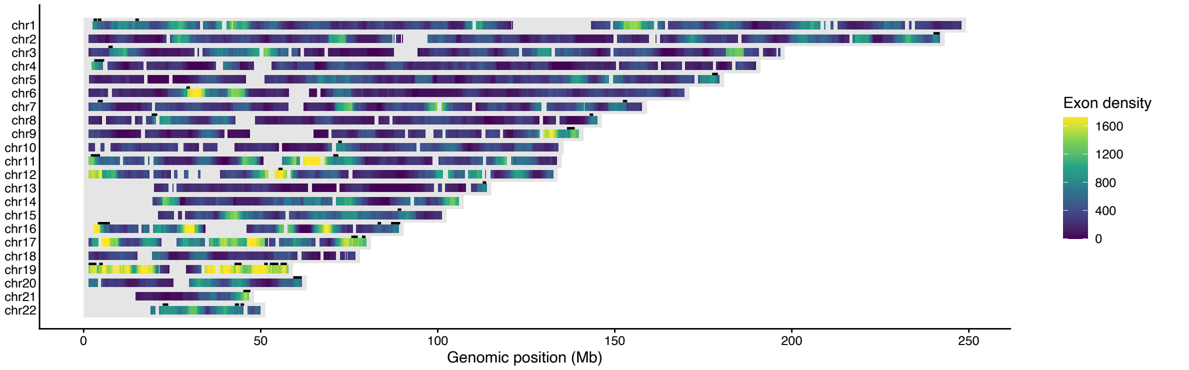


Recombination rate

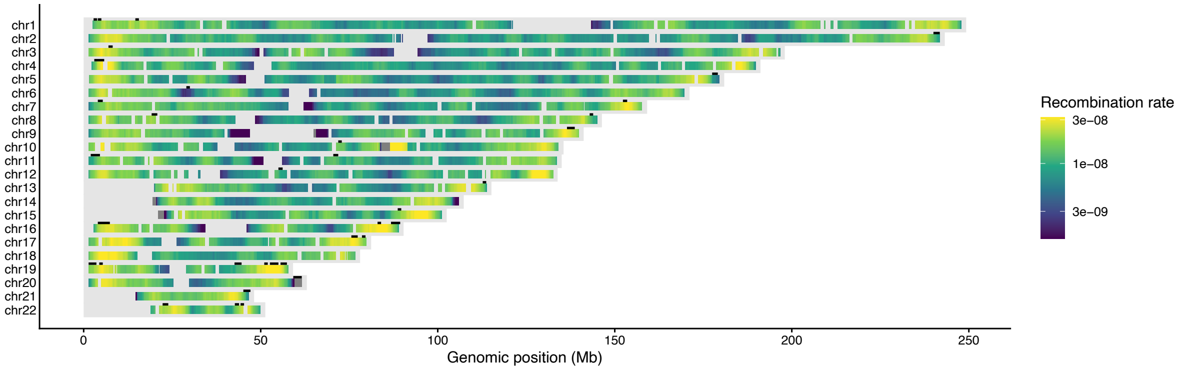


Neanderthal ancestry

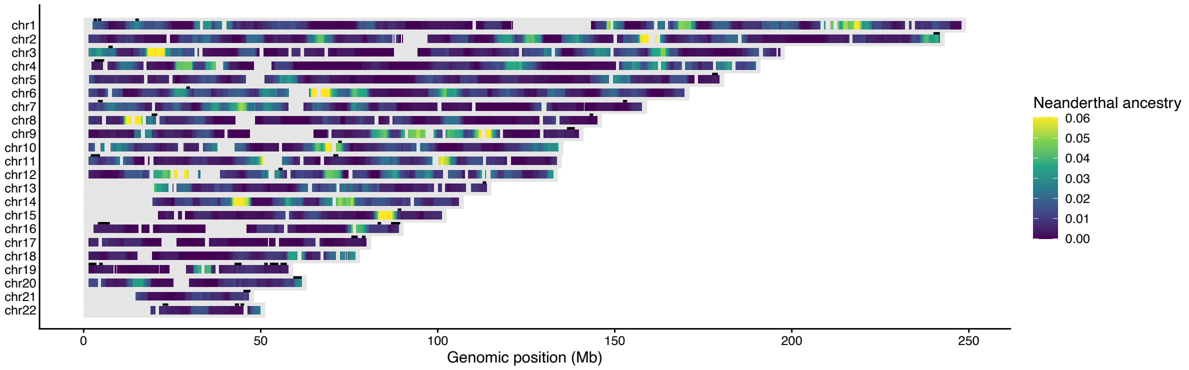


DominL probability

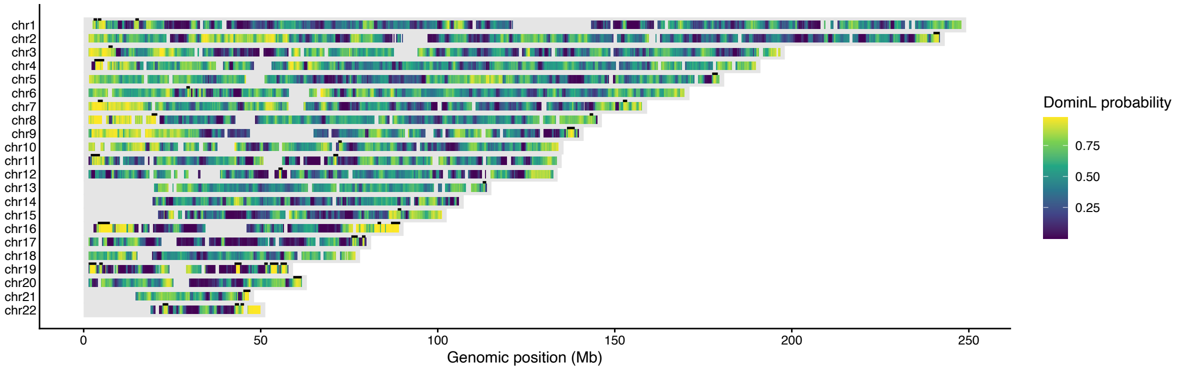

#### b) Median distance to chromosome end across all genomic windows, stratified by exon density

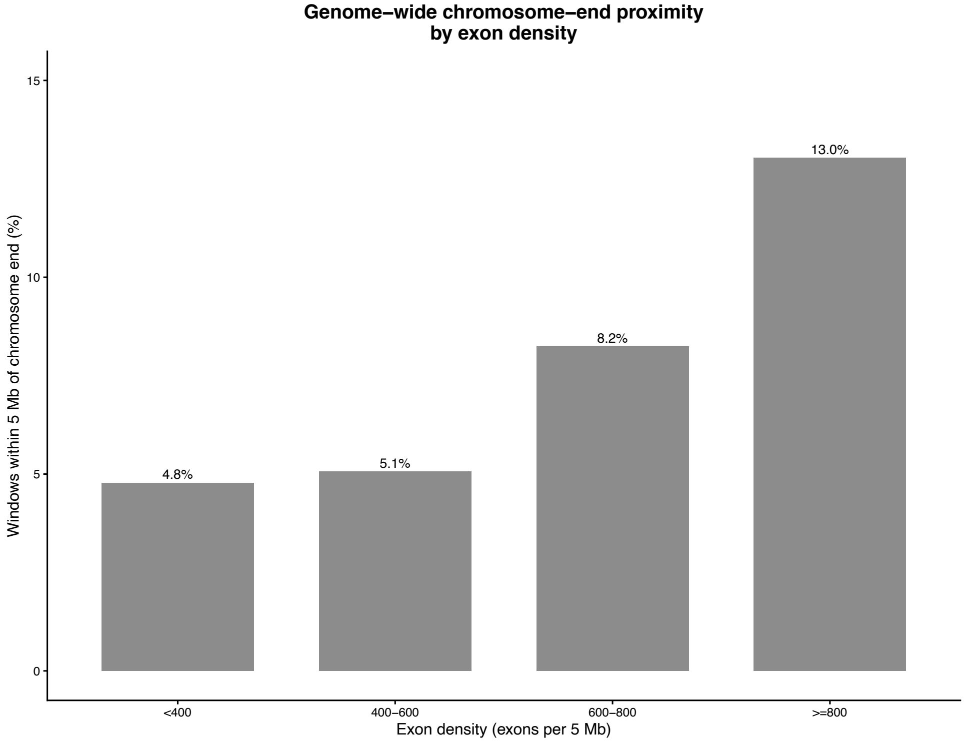


#### c) Fraction within 5 Mb of chromosome ends for DominL-predicted vs other windows within each exon-density bin

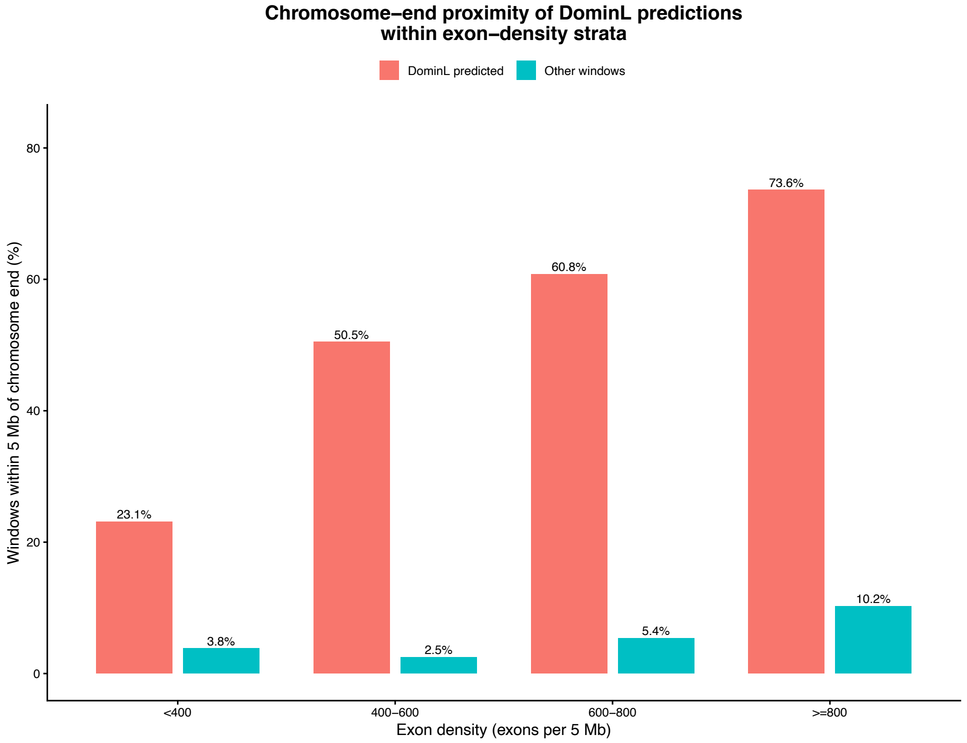

**Supplementary Figure 23: GO enrichment test for different groups of recessive windows**

a) **Molecular functions enriched in recessive regions that are found in European populations only (region detected in at least one population among CEU, GBR, FIN, IBS)**

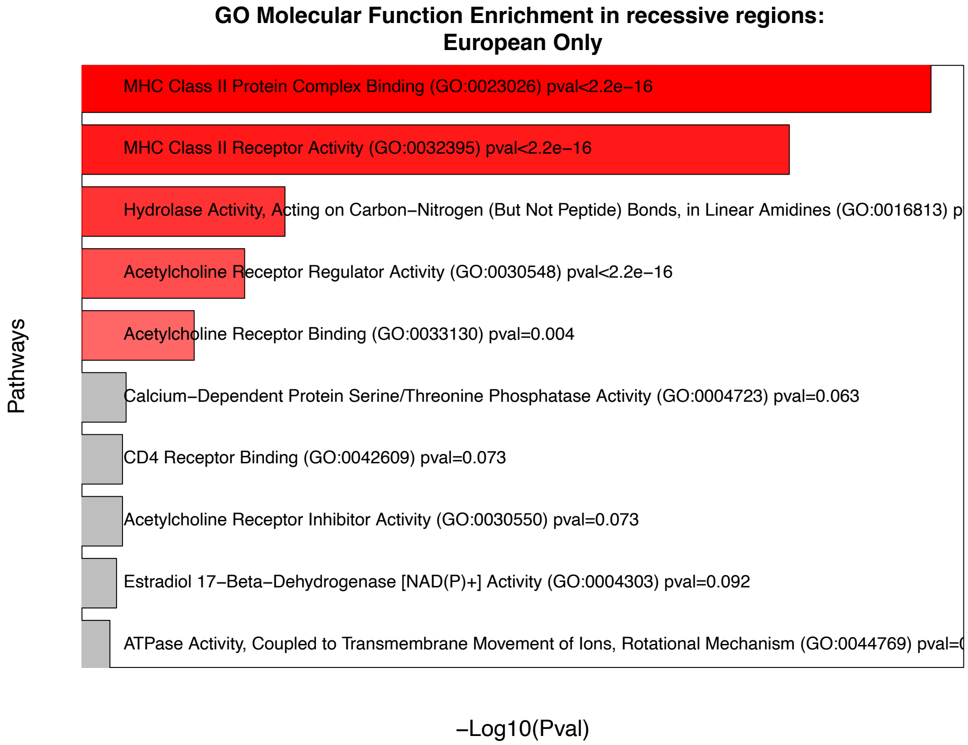


b) **Biological processes enriched in recessive regions that are found in European populations only (region detected in at least one population among CEU, GBR, FIN, IBS)**

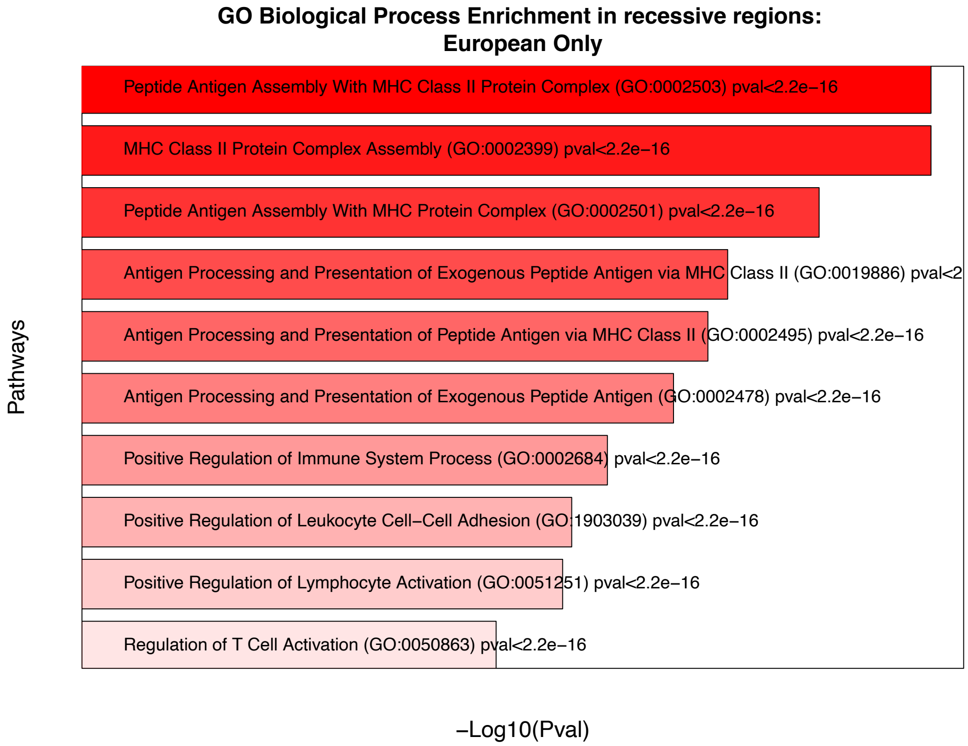


#### c) Molecular functions enriched in recessive regions that are found in Asian populations only (region detected in at least one population among CHB, CHS, JPT)

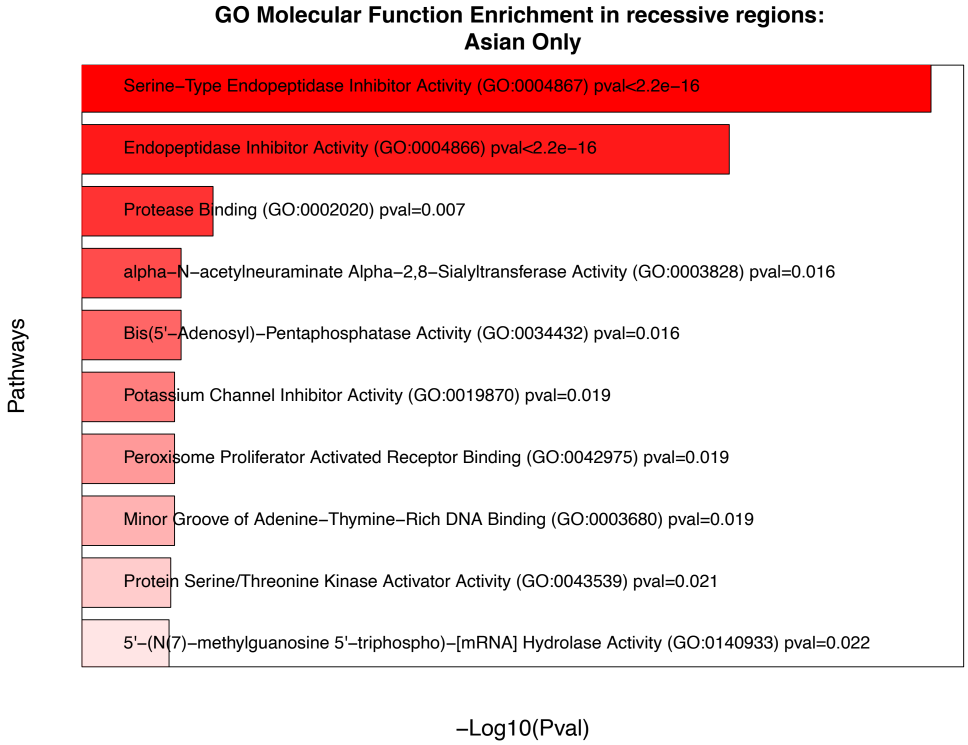

d) **Biological processes enriched in recessive regions that are found in Asian populations only (region detected in at least one population among CHB, CHS, JPT)**

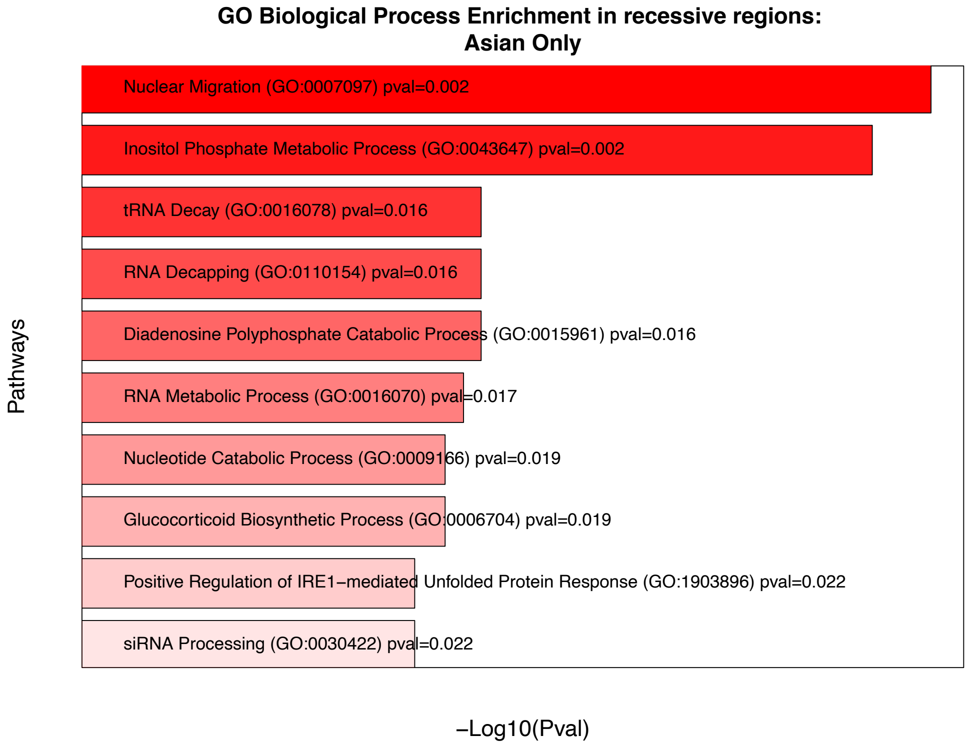


*Gene Oncology (GO) enrichment analyses using Enrichr for (**a**) molecular functions in recessive regions found in European populations only, (**b**) human biological processes in recessive regions found in European populations only, (**c**) molecular functions in recessive regions found in Asian populations only, and (**d**) human biological processes in recessive regions found in Asian populations only. The top 10 processes are shown, with ascending p-values for each analysis, with red indicating a significant p-value and gray indicating non-significant p-values. The intensity of the red color as well as the length of the bars both indicate how significant a process is from the enrichment test.*

**Supplementary Table 1: Feature list, importance score, and feature selection results**

| Feature | Description | Importance Score | If passed feature selection |
|---|---|---|---|
| num_seg_p2 | Number of segregating sites in outgroup population | 0.180 | X |
| **num_private_seg_p2** | **Number of private segregating sites in outgroup population** | **0.147** | **Yes** |
| **exon_window** | **Exon Density in 1MB window** | **0.109** | **Yes** |
| num_variant_window | Number of segregating sites in 1MB window | 0.092 | X |
| **mean_introg_anc** | **Mean Neanderthal introgressed ancestry in 5MB window** | **0.088** | **Yes** |

| | | | |
|---|---|---|---|
| **watterson_theta_p3** | **Watterson's estimate of genetic diversity in recipient population** | **0.066** | **Yes** |
| **divergence_p3_p1** | **Nucleotide divergence between recipient and donor populations** | **0.054** | **Yes** |
| **exon_density** | **Exon Density in 5MB window** | **0.028** | **Yes** |
| **num_private_seg_p3** | **Number of private segregating sites in recipient population** | **0.028** | **Yes** |
| **divergence_p3_p2** | **Nucleotide divergence between recipient and outgroup populations** | **0.026** | **Yes** |
| **num_seg_p1** | **Number of segregating sites in Neanderthal (donor) population** | **0.023** | **Yes** |
| recrate_window | Mean recombination rates in 1MB window | 0.017 | X |
| mean_recrate | Mean recombination rates in 5MB window | 0.015 | X |
| df_p3_p1 | Density of fixed differences between donor and recipient populations | 0.014 | X |
| **U50** | **Number of U50 alleles in recipient population (Racimo et al. 2016)** | **0.009** | **Yes** |
| **U0** | **Number of U0 alleles in recipient population (Racimo et al. 2016)** | **0.009** | **Yes** |
| **Het** | **Heterozygosity** | **0.009** | **Yes** |
| **num_seg_p3** | **Number of segregating sites in recipient population** | **0.008** | **Yes** |
| garud_h1 | H1 statistic for detecting soft sweeps (Garud et al. 2015) | 0.008 | X |
| RD | Sequence Divergence Ratio between donor and recipient population (Racimo et al. 2016) | 0.008 | X |
| **D** | **ABBA-BABA Statistic (Green et al. 2010)** | **0.007** | **Yes** |
| hap_diversity_p3 | Haplotype diversity in recipient population | 0.007 | X |
| df_p3_p2 | Density of fixed differences between recipient and outgroup populations | 0.007 | X |
| **Q95** | **95% quantile of derived allele frequency in recipient population (Racimo et al. 2016)** | **0.007** | **Yes** |
| **introg_anc_window** | **mean Neanderthal introgressed ancestry in 1MB window** | **0.007** | **Yes** |

| | | | |
|---|---|---|---|
| **U20** | **Number of U20 alleles in recipient population (Racimo et al. 2016)** | **0.006** | **Yes** |
| fD | ABBA-BABA Statistic (Martin et al. 2014) | 0.006 | X |
| **num_private_seg_p1** | **Number of private segregating sites in Neanderthal (donor) population** | **0.006** | **Yes** |
| **windowed_tajima_d_p3** | **Tajima's D in recipient population** | **0.006** | **Yes** |
| **U80** | **Number of U80 alleles in recipient population (Racimo et al. 2016)** | **0.005** | **Yes** |
| garud_h2_h1 | H2/H1 statistic for detecting soft sweeps (Garud et al. 2015) | 0.000 | X |
| garud_h12 | H12 statistic for detecting soft sweeps (Garud et al. 2015) | 0.000 | X |

*This table summarizes all features used to train DominL. From left to right, the columns show the name of the feature, the biological meaning, feature importance score in XGBoost before feature selection, and whether a given feature is retained after feature selection.*

**Supplementary Table 2: Summary of recessive regions detected by *DominL* ($P_{recessive}$ > 0.9).**

| Chromosome | Start position | End position | Exon density (number of exons per 5MB) | Detected in populations |
|---|---|---|---|---|
| **1** | **2200000** | **3800000** | **837** | **CEU,FIN,IBS,GBR,CHB,CHS,JPT** |
| **1** | **4000000** | **6400000** | **611** | **CEU,FIN,IBS,GBR,CHB,CHS,JPT** |
| **1** | **13800000** | **15800000** | **862** | **CEU,FIN,GBR,CHB,CHS,JPT** |
| **1** | **17600000** | **19200000** | **634** | **CEU,FIN,IBS** |
| 1 | 102400000 | 103600000 | 273 | CEU,GBR |
| 1 | 105600000 | 107200000 | 132 | CEU,FIN,IBS,GBR,CHB,CHS,JPT |
| 2 | 1000000 | 3400000 | 225 | CEU,FIN,IBS,GBR,CHB,CHS,JPT |
| 2 | 30600000 | 31600000 | 561 | GBR,CHB,JPT |
| 2 | 33000000 | 36600000 | 446 | CEU,FIN,IBS,GBR,CHB,CHS,JPT |
| 2 | 37400000 | 38800000 | 280 | CEU,FIN,IBS,GBR,JPT |
| 2 | 39800000 | 42800000 | 336 | CEU,FIN,IBS,GBR,CHB,CHS,JPT |
| 2 | 43000000 | 44000000 | 344 | CEU,CHB,CHS |
| 2 | 46400000 | 48000000 | 384 | CEU,FIN,IBS,CHB,CHS,JPT |

| | | | | |
|---|---|---|---|---|
| 2 | 51000000 | 54400000 | 254 | CEU,FIN,IBS,GBR,CHB,CHS,JPT |
| 2 | 56200000 | 58000000 | 272 | CEU,FIN,IBS,GBR,CHB,CHS,JPT |
| 2 | 75600000 | 78000000 | 347 | CEU,FIN,IBS,GBR,CHB,CHS |
| 2 | 236200001 | 238600001 | 597 | FIN,IBS,CHS,JPT |
| **2** | **239800001** | **242000001** | **848** | **CEU,FIN,IBS,GBR,CHB,CHS,JPT** |
| 3 | 1000000 | 4800000 | 397 | CEU,FIN,IBS,GBR,CHB,CHS,JPT |
| **3** | **1200000** | **8200000** | **778** | **CEU,FIN,IBS,GBR,CHB,CHS,JPT** |
| 3 | 20800000 | 23400000 | 105 | CEU,FIN,IBS,GBR,CHB,CHS,JPT |
| 3 | 24200000 | 25800000 | 99 | CEU,FIN,IBS,GBR,CHB,CHS,JPT |
| 3 | 59200000 | 61000000 | 548 | CEU,FIN,IBS,GBR,CHB,CHS,JPT |
| 3 | 74600000 | 75600000 | 51 | CHB,CHS,JPT |
| 3 | 125100001 | 126300001 | 375 | CEU,FIN,IBS,GBR,CHB,CHS,JPT |
| 3 | 189700001 | 190700001 | 295 | CHB |
| 3 | 193700001 | 194900001 | 144 | CEU,FIN,IBS,GBR,CHB,JPT |
| 3 | 195100001 | 196900001 | 344 | CEU,FIN,IBS,GBR,JPT |
| **4** | **3000000** | **5800000** | **937** | **CEU,FIN,IBS,GBR,CHB,CHS,JPT** |
| 4 | 6400000 | 10800000 | 354 | CEU,FIN,IBS,GBR,CHB,CHS,JPT |
| 4 | 11000000 | 12800000 | 28 | CEU,FIN,IBS,GBR |
| 4 | 57500001 | 59700001 | 306 | CEU,FIN,IBS,GBR,CHB,CHS,JPT |
| 4 | 59900001 | 61900001 | 342 | CEU,FIN,IBS,GBR,CHB,CHS,JPT |
| 4 | 63100001 | 64900001 | 342 | CEU,FIN,IBS,GBR,CHB,CHS,JPT |
| 4 | 69500001 | 70500001 | 442 | CHS |
| 4 | 188500001 | 190300001 | 186 | CEU,FIN,IBS |
| 5 | 1200000 | 3800000 | 236 | CEU,FIN,IBS,GBR,CHB,CHS,JPT |
| 5 | 45200000 | 46200000 | 76 | CEU,GBR,CHB,JPT |
| 5 | 109100001 | 110500001 | 260 | CEU,FIN,IBS,GBR,CHB,CHS,JPT |
| 5 | 111100001 | 112100001 | 297 | CHS |
| 5 | 113300001 | 116300001 | 190 | CEU,FIN,IBS,GBR,CHB,CHS,JPT |
| 5 | 116900001 | 118700001 | 273 | CEU,FIN,IBS,GBR,CHS,JPT |
| **5** | **177500001** | **179100001** | **981** | **CEU,FIN,GBR,CHB,CHS,JPT** |
| 6 | 6000000 | 7000000 | 207 | GBR,CHB,CHS |
| 6 | 23000000 | 24800000 | 406 | CEU,FIN,IBS,GBR,CHB,CHS,JPT |
| **6** | **28800000** | **30600000** | **1609** | **CEU,FIN,GBR,CHB,CHS,JPT** |
| **6** | **32200000** | **33200000** | **1681** | **FIN** |

| | | | | |
|---|---|---|---|---|
| **6** | **33400000** | **34400000** | **1467** | **CHB** |
| 6 | 66500001 | 68700001 | 59 | CEU,FIN,IBS,GBR,CHB,CHS,JPT |
| 6 | 86500001 | 87700001 | 162 | CEU,IBS,GBR |
| 6 | 150100001 | 151300001 | 249 | FIN,IBS,GBR,CHB,CHS,JPT |
| 6 | 166500001 | 167900001 | 276 | CEU,FIN,IBS,GBR,CHS,JPT |
| 6 | 168300001 | 170100001 | 391 | CEU,FIN,GBR,CHS |
| **7** | **1000000** | **17200000** | **634** | **CEU,FIN,IBS,GBR,CHB,CHS,JPT** |
| 7 | 21200000 | 22600000 | 414 | CHB,CHS,JPT |
| 7 | 45200000 | 46200000 | 583 | FIN |
| 7 | 51000000 | 54000000 | 139 | CEU,FIN,IBS,GBR,CHB,CHS,JPT |
| 7 | 61700001 | 63100001 | 56 | CEU,FIN,IBS,GBR,CHB,CHS,JPT |
| 7 | 67300001 | 68500001 | 195 | CEU,FIN,IBS,GBR,CHB,CHS,JPT |
| 7 | 146100001 | 147100001 | 159 | CEU |
| **7** | **149300001** | **150300001** | **720** | **IBS** |
| **7** | **152100001** | **153700001** | **665** | **CEU,FIN,IBS,GBR,CHS** |
| 7 | 154500001 | 156300001 | 318 | CEU,FIN,IBS,GBR,CHB,CHS,JPT |
| 7 | 156500001 | 158100001 | 281 | CEU,FIN,IBS,GBR,CHB,CHS,JPT |
| 8 | 1000000 | 5600000 | 210 | CEU,FIN,IBS,GBR,CHB,CHS,JPT |
| 8 | 6000000 | 7400000 | 104 | CEU,FIN,IBS,GBR,CHB,CHS,JPT |
| 8 | 8000000 | 13800000 | 337 | CEU,FIN,IBS,GBR,CHB,CHS,JPT |
| 8 | 14200000 | 17000000 | 291 | CEU,FIN,IBS,GBR,CHB,CHS,JPT |
| **8** | **17200000** | **20800000** | **864** | **CEU,FIN,IBS,GBR,CHB,CHS,JPT** |
| **8** | **142900001** | **143900001** | **724** | **CEU** |
| 9 | 6000000 | 13000000 | 237 | CEU,FIN,IBS,GBR,CHB,CHS,JPT |
| 9 | 13600000 | 15800000 | 87 | CEU,FIN,IBS,GBR,CHB,CHS,JPT |
| 9 | 17000000 | 19800000 | 279 | CEU,FIN,IBS,GBR,CHB,CHS,JPT |
| 9 | 22200000 | 23800000 | 107 | FIN,CHB,CHS,JPT |
| 9 | 24000000 | 25200000 | 109 | CHB,CHS,JPT |
| 9 | 25600000 | 27400000 | 142 | CEU,IBS,GBR,CHB |
| 9 | 82900001 | 84100001 | 81 | CEU,FIN |
| 9 | 105300001 | 106700001 | 318 | IBS |
| **9** | **136500001** | **139100001** | **1384** | **CEU,FIN,IBS,GBR,CHB,CHS,JPT** |
| 10 | 2000000 | 3400000 | 283 | CEU,FIN,CHB,CHS,JPT |
| 10 | 4400000 | 5800000 | 360 | CEU,FIN,IBS,GBR,CHB,CHS,JPT |

| | | | | |
|---|---|---|---|---|
| 10 | 7600000 | 8600000 | 245 | CEU,FIN |
| 10 | 13800000 | 17600000 | 347 | CEU,FIN,IBS,GBR,CHB,CHS,JPT |
| 10 | 18600000 | 21600000 | 349 | CEU,FIN,IBS,GBR,CHB,CHS,JPT |
| 10 | 42300001 | 43500001 | 105 | CEU,FIN,IBS,GBR,CHB,CHS,JPT |
| 10 | 67500001 | 69100001 | 285 | CEU,FIN,GBR,CHB,CHS,JPT |
| **10** | **71900001** | **72900001** | **842** | **CEU,FIN,IBS,GBR,CHB** |
| 10 | 131700001 | 134500001 | 312 | CEU,FIN,IBS,GBR,CHB,CHS,JPT |
| **11** | **2000000** | **6200000** | **1354** | **CEU,FIN,IBS,GBR,CHB,CHS,JPT** |
| 11 | 49600000 | 51000000 | 66 | CEU,FIN,IBS,GBR,CHB,CHS,JPT |
| **11** | **70500001** | **71900001** | **794** | **CEU,FIN,IBS,GBR,CHB,CHS,JPT** |
| 11 | 80700001 | 81900001 | 107 | FIN |
| 11 | 97900001 | 98900001 | 118 | CHS |
| 11 | 130900001 | 131900001 | 161 | CEU,FIN,IBS,GBR |
| 12 | 38200001 | 39400001 | 256 | CHS,JPT |
| **12** | **55000000** | **56200001** | **2111** | **CEU,FIN,IBS,GBR,CHS,JPT** |
| 12 | 126200001 | 127800001 | 197 | CEU,FIN,IBS,CHB,CHS,JPT |
| 12 | 129000001 | 130800001 | 336 | CHS,JPT |
| 12 | 131000001 | 132200001 | 589 | CEU,FIN,GBR,CHB,CHS,JPT |
| 13 | 23300001 | 24300001 | 264 | FIN |
| **13** | **112700001** | **113700001** | **743** | **CEU,FIN,GBR,CHB,CHS,JPT** |
| 14 | 42100001 | 44300001 | 135 | CEU,FIN,IBS,GBR,CHB,CHS,JPT |
| **14** | **94700001** | **96700001** | **607** | **CHS,JPT** |
| 14 | 98700001 | 99700001 | 360 | CHB |
| 15 | 23700001 | 25700001 | 306 | CEU,FIN,IBS,GBR,CHB,CHS,JPT |
| 15 | 57700001 | 58900001 | 436 | CEU,FIN,IBS,GBR,CHB,CHS,JPT |
| **15** | **85900001** | **89700001** | **710** | **CEU,IBS,GBR,CHB,CHS,JPT** |
| **15** | **92300001** | **93300001** | **624** | **CHB,CHS,JPT** |
| 15 | 97900001 | 99500001 | 207 | CEU,GBR,CHS,JPT |
| 15 | 99700001 | 101500001 | 227 | CEU,FIN,IBS,GBR,CHB,CHS,JPT |
| **16** | **3800000** | **8600000** | **1701** | **CEU,FIN,IBS,GBR,CHB,CHS,JPT** |
| 16 | 8800000 | 14200000 | 460 | CEU,FIN,IBS,GBR,CHB,CHS,JPT |
| 16 | 74600001 | 76400001 | 223 | CEU,FIN,IBS,GBR,CHB,CHS,JPT |
| **16** | **76600001** | **89400001** | **738** | **CEU,FIN,IBS,GBR,CHB,CHS,JPT** |
| 17 | 14200000 | 15200000 | 307 | JPT |

| | | | | |
|---|---|---|---|---|
| **17** | **75600001** | **77400001** | **1421** | **CEU,FIN,IBS,GBR,CHB,CHS,JPT** |
| **17** | **78600001** | **79600001** | **1185** | **CEU,FIN,IBS,GBR,CHB,CHS,JPT** |
| 18 | 9800000 | 11200000 | 492 | CEU,FIN,GBR,CHB,CHS,JPT |
| 18 | 13400000 | 15200000 | 352 | CEU,FIN,IBS,GBR,CHB,CHS,JPT |
| 18 | 70200001 | 71400001 | 108 | JPT |
| 18 | 75400001 | 77200001 | 309 | CEU,FIN,IBS,GBR,CHB,CHS,JPT |
| **19** | **1400000** | **3800000** | **1730** | **CEU,FIN,IBS,GBR,CHB,CHS,JPT** |
| **19** | **4400000** | **5400000** | **1369** | **CEU,FIN,IBS,JPT** |
| 19 | 21800000 | 23200000 | 221 | CEU,FIN,IBS,GBR,CHB,CHS,JPT |
| 19 | 28600001 | 30400001 | 71 | CEU,IBS,GBR,CHS,JPT |
| **19** | **42600001** | **44600001** | **1689** | **CEU,FIN,IBS,GBR,CHB,CHS,JPT** |
| **19** | **51000000** | **52200001** | **1641** | **CEU,FIN,GBR,CHB,CHS,JPT** |
| **19** | **52600001** | **55000001** | **1593** | **CEU,FIN,IBS,GBR,CHB,CHS,JPT** |
| **19** | **55600001** | **57400001** | **1682** | **CEU,FIN,IBS,GBR,CHB,CHS,JPT** |
| 20 | 22800000 | 25000000 | 148 | CEU,FIN,IBS,GBR,CHB,CHS,JPT |
| 20 | 46400001 | 47400001 | 468 | CEU,GBR |
| **20** | **59200001** | **61800001** | **801** | **CEU,FIN,IBS,GBR,CHB,CHS,JPT** |
| 21 | 17700001 | 18900001 | 127 | GBR,CHB,CHS,JPT |
| **21** | **44700001** | **47100001** | **1525** | **CEU,FIN,IBS,GBR,CHB,CHS,JPT** |
| **22** | **22300001** | **24100001** | **939** | **CEU,FIN,IBS,GBR** |
| **22** | **42700001** | **43900001** | **704** | **CEU,FIN,IBS,GBR,CHB,CHS,JPT** |
| **22** | **44300001** | **45500001** | **632** | **CEU,FIN,IBS,GBR,CHB,CHS,JPT** |
| 22 | 46100001 | 50300001 | 589 | CEU,FIN,IBS,GBR,CHB,CHS,JPT |

*From left to right, each column corresponds to the chromosome name, start position, end position, exon density (number of exons per 5MB window), and the population(s) the region is detected in. All positions refer to hg19. In the main text, we only discuss regions that have exon density >=600 (highlighted in bold). We also merged overlapping windows in each row, so the total length of each region can exceed 1MB. We show the mean exon density of all overlapping windows in the 4$^{th}$ column.*

#### Supplementary Table 3: Sensitivity of empirical analyses to chromosome-end distance

| Analysis | Metric | Original effect | Original CI | Original pval | With chrom end effect | With chrom end CI | With chrom end pval |
|---|---|---|---|---|---|---|---|
| B value | delta mean | 0.05 | 0.0, 0.1 | 0.0002* | 0.04 | 0.02, 0.07 | 0.0045* |
| LOEUF | delta mean | 0.13 | 0.022, 0.211 | 0.0002* | 0.06 | -0.04, 0.16 | 0.277 |
| UKBB non-additive | IRR | 1.48 | 1.44, 1.52 | 4.46e-184* | 1.41 | 1.37, 1.45 | 1.92e-145* |
| UKBB additive | IRR | 0.93 | 0.78, 1.10 | 0.38 | 0.94 | 0.79, 1.12 | 0.495 |
| Intermediate ROH count | delta mean | -1.55 | -2.37, -0.81 | 0.0002* | -0.89 | -1.74, -0.05 | 0.067 |
| Intermediate ROH total | delta MB | -0.58 | -0.90, -0.29 | 0.0004* | -0.29 | -0.60, 0.04 | 0.116 |
| Intermediate ROH max | delta MB | -0.19 | -0.26, -0.12 | 0.0002* | -0.12 | -0.18, -0.05 | 0.0028* |
| Long ROH count | delta mean | -0.24 | -0.41, -0.08 | 0.02* | -0.16 | -0.33, 0.00 | 0.087 |
| Long ROH total | delta MB | -0.18 | -0.32, -0.05 | 0.025* | -0.12 | -0.26, 0.02 | 0.137 |
| Long ROH max | delta MB | -0.01 | -0.16, -0.03 | 0.012* | -0.09 | -0.16, -0.01 | 0.023* |

**annotates significant p-values.*

**Supplementary Table 4: GO biological processes enriched in recessive regions (showing all significant pathways)**

| Term | Overlap | P-value | Odds Ratio | Combined Score | Genes |
|---|---|---|---|---|---|
| Peptide Antigen Assembly With MHC Class II Protein Complex (GO:0002503) | 11/13 | 1.24E-20 | 536.019704 | 24568.03127 | *HLA-DRB5;HLA-DMA;HLA-DMB;HLA-DRA;HLA-DOA;HLA-DOB;HLA-DQA2;HLA-DQA1;HLA-DQB2;HLA-DRB1;HLA-DPA1* |
| MHC Class II Protein Complex Assembly (GO:0002399) | 11/13 | 1.24E-20 | 536.019704 | 24568.03127 | *HLA-DRB5;HLA-DMA;HLA-DMB;HLA-DRA;HLA-DOA;HLA-DOB;HLA-DQA2;HLA-* |

| | | | | | |
|---|---|---|---|---|---|
| | | | | | *DQA1;HLA-DQB2;HLA-DRB1;HLA-DPA1* |
| Peptide Antigen Assembly With MHC Protein Complex (GO:0002501) | 11/18 | 4.84E-18 | 153.109782 | 6104.402022 | *HLA-DRB5;HLA-DMA;HLA-DMB;HLA-DRA;HLA-DOA;HLA-DOB;HLA-DQA2;HLA-DQA1;HLA-DQB2;HLA-DRB1;HLA-DPA1* |
| Antigen Processing and Presentation of Exogenous Peptide Antigen via MHC Class II (GO:0019886) | 11/25 | 6.35E-16 | 76.5277973 | 2677.908788 | *HLA-DMA;HLA-DRB5;HLA-DMB;HLA-DRA;HLA-DOA;HLA-DOB;HLA-DQA2;HLA-DQA1;HLA-DQB2;HLA-DRB1;HLA-DPA1* |
| Antigen Processing and Presentation of Peptide Antigen via MHC Class II (GO:0002495) | 11/27 | 1.82E-15 | 66.9550493 | 2272.320455 | *HLA-DMA;HLA-DRB5;HLA-DMB;HLA-DRA;HLA-DOA;HLA-DOB;HLA-DQA2;HLA-DQA1;HLA-DQB2;HLA-DRB1;HLA-DPA1* |
| Antigen Processing and Presentation of Exogenous Peptide Antigen (GO:0002478) | 1/11 | 1.14E-14 | 53.553202 | 1719.294065 | *HLA-DMA;HLA-DRB5;HLA-DMB;HLA-DRA;HLA-DOA;HLA-DOB;HLA-DQA2;HLA-DQA1;HLA-DQB2;HLA-DRB1;HLA-DPA1* |

| | | | | | |
|---|---|---|---|---|---|
| Positive Regulation of Immune System Process (GO:0002684) | 1/12 | 3.88E-13 | 35.6840722 | 1019.794535 | *HLA-DMA;HLA-DRB5;HLA-DMB;HLA-DRA;HLA-DOA;HLA-DOB;HLA-DQA2;HLA-DQA1;HLA-DQB2;HLA-DRB1;HLA-DPA1* |
| Positive Regulation of Leukocyte Cell-Cell Adhesion (GO:1903039) | 1/13 | 2.60E-12 | 28.9227799 | 771.5574571 | *HLA-DMA;HLA-DRB5;HLA-DMB;HLA-DRA;HLA-DOA;HLA-DOB;HLA-DQA2;HLA-DQA1;HLA-DQB2;HLA-DRB1;HLA-DPA1* |
| Positive Regulation of Lymphocyte Activation (GO:0051251) | 1/14 | 4.21E-12 | 27.4367816 | 718.6364144 | *HLA-DMA;HLA-DRB5;HLA-DMB;HLA-DRA;HLA-DOA;HLA-DOB;HLA-DQA2;HLA-DQB2;HLA-DQA1;HLA-DRB1;HLA-DPA1* |
| Regulation of T Cell Activation (GO:0050863) | 1/15 | 1.46E-10 | 18.755423 | 424.7344451 | *HLA-DMA;HLA-DRB5;HLA-DMB;HLA-DRA;HLA-DOA;HLA-DOB;HLA-DQA2;HLA-DQB2;HLA-DQA1;HLA-DRB1;HLA-DPA1* |
| Positive Regulation of Response to Stimulus (GO:0048584) | 1/16 | 4.10E-09 | 13.1822052 | 254.5863873 | *HLA-DMA;HLA-DRB5;HLA-DMB;HLA-DRA;HLA-DOA;HLA-DOB;HLA-DQA2;HLA-DQB2;HLA-DQA1;HLA-DRB1;HLA-DPA1* |

| | | | | | |
|---|---|---|---|---|---|
| Regulation of Immune Response (GO:0050776) | 1/17 | 5.17E-09 | 12.863256 | 245.4465432 | *HLA-DMA;HLA-DRB5;HLA-DMB;HLA-DRA;HLA-DOA;HLA-DOB;HLA-DQA2;HLA-DQB2;HLA-DQA1;HLA-DRB1;HLA-DPA1* |
| Positive Regulation of Immune Response (GO:0050778) | 1/18 | 5.79E-09 | 12.7094769 | 241.0668329 | *HLA-DMA;HLA-DRB5;HLA-DMB;HLA-DRA;HLA-DOA;HLA-DOB;HLA-DQA2;HLA-DQB2;HLA-DQA1;HLA-DRB1;HLA-DPA1* |
| Positive Regulation of T Cell Activation (GO:0050870) | 11/111 | 3.02E-08 | 10.6672906 | 184.7036232 | *HLA-DRB5;HLA-DMA;HLA-DMB;HLA-DRA;HLA-DOA;HLA-DOB;HLA-DQA2;HLA-DQA1;HLA-DRB1;HLA-DQB2;HLA-DPA1* |
| + Reg of CD4-+, CD25-+, Alpha-Beta Regulatory T Cell Differentiation (GO:0032831) | 2/5 | 0.00111561 | 62.2106918 | 422.9302774 | *HLA-DRA;HLA-DRB1* |
| Regulation of T-helper Cell Differentiation (GO:0045622) | 2/6 | 0.00166162 | 46.6556604 | 298.5943724 | *HLA-DRA;HLA-DRB1* |

| | | | | | |
|---|---|---|---|---|---|
| Positive Regulation of CD4-positive, Alpha-Beta T Cell Differentiation (GO:0043372) | 2/7 | 0.00230989 | 37.3226415 | 226.5691213 | *HLA-DRA;HLA-DRB1* |
| Regulation of CD4-positive, Alpha-Beta T Cell Activation (GO:2000514) | 2/7 | 0.00230989 | 37.3226415 | 226.5691213 | *HLA-DRA;HLA-DRB1* |
| Positive Regulation of Memory T Cell Differentiation (GO:0043382) | 2/8 | 0.00305819 | 31.1006289 | 180.0705493 | *HLA-DRA;HLA-DRB1* |
| Regulation of Memory T Cell Differentiation (GO:0043380) | 2/9 | 0.00390431 | 26.6563342 | 147.8273119 | *HLA-DRA;HLA-DRB1* |
| Positive Regulation of Immune Effector Process (GO:0002699) | 1/3 | 0.00665041 | 8.51055579 | 42.66407473 | *HLA-DMB;HLA-DRA;HLA-DRB1* |
| Positive Regulation of CD4-positive, Alpha-Beta T Cell Activation (GO:2000516) | 2/12 | 0.00700826 | 18.6566038 | 92.5491674 | *HLA-DRA;HLA-DRB1* |
| Regulation of CD4-positive, Alpha-Beta T Cell Differentiation (GO:0043370) | 2/12 | 0.00700826 | 18.6566038 | 92.5491674 | *HLA-DRA;HLA-DRB1* |

| | | | | | |
|---|---|---|---|---|---|
| Regulation of Immune Effector Process (GO:0002697) | 2/14 | 0.00952804 | 15.5455975 | 72.34168303 | *HLA-DRA;HLA-DRB1* |
| Positive Regulation of Regulatory T Cell Differentiation (GO:0045591) | 2/17 | 0.01394328 | 12.4345912 | 53.1299907 | *HLA-DRA;HLA-DRB1* |
| Positive Regulation of Alpha-Beta T Cell Activation (GO:0046635) | 2/18 | 0.01557677 | 11.6568396 | 48.51546698 | *HLA-DRA;HLA-DRB1* |
| Immunoglobulin Mediated Immune Response (GO:0016064) | 2/19 | 0.017288 | 10.9705882 | 44.51582211 | *IGLL5;IGLL1* |
| B Cell Mediated Immunity (GO:0019724) | 2/20 | 0.01907514 | 10.360587 | 41.0213934 | *IGLL5;IGLL1* |
| Antigen Processing and Presentation of Endogenous Peptide Antigen (GO:0002483) | 2/21 | 0.02093637 | 9.81479643 | 37.9466293 | *HLA-DRA;HLA-DRB1* |
| Focal Adhesion Assembly (GO:0048041) | 2/22 | 0.02286992 | 9.32358491 | 35.22387553 | *BCR;RCC2* |
| Regulation of Neurotransmitter Receptor Activity (GO:0099601) | 2/23 | 0.02487406 | 8.87915544 | 32.79897702 | *LYNX1;PSCA* |

| | | | | | |
|---|---|---|---|---|---|
| Cellular Response to Acetylcholine (GO:1905145) | 2/24 | 0.02694706 | 8.47512864 | 30.62810787 | *LYNX1;PSCA* |
| Cell-Substrate Junction Assembly (GO:0007044) | 1/2 | 0.03828686 | 6.90391335 | 22.52504262 | *BCR;RCC2* |
| Regulation of Chemotaxis (GO:0050920) | 1/3 | 0.04073854 | 6.65700809 | 21.3062915 | *RARRES2;PPM1F* |
| Positive Regulation of Locomotion (GO:0040017) | 1/4 | 0.04581325 | 6.21257862 | 19.15451 | *RARRES2;PPM1F* |
| Positive Regulation of T Cell Mediated Cytotoxicity (GO:0001916) | 1/5 | 0.04581325 | 6.21257862 | 19.15451 | *HLA-DRA;HLA-DRB1* |

**Supplementary Table 5: GO molecular functions enriched in recessive regions (showing all significant pathways)**

| Term | Overlap | P-value | Odds Ratio | Combined Score | Genes |
|---|---|---|---|---|---|
| MHC Class II Protein Complex Binding (GO:0023026) | 11/24 | 3.59E-16 | 82.4187192 | 2931.06645 | *HLA-DRB5;HLA-DMA;HLA-DMB;HLA-DRA;HLA-DOA;HLA-DOB;HLA-DQA2;HLA-DQA1;HLA-DQB2;HLA-DRB1;HLA-DPA1* |

| | | | | | |
|---|---|---|---|---|---|
| MHC Class II Receptor Activity (GO:0032395) | 7/8 | 1.15E-13 | 669.057971 | 19931.7407 | *HLA-DRA;HLA-DOB;HLA-DQA2;HLA-DQA1;HLA-DQB2;HLA-DRB1;HLA-DPA1* |
| Hydrolase Activity, Acting on Carbon-Nitrogen (But Not Peptide) Bonds, in Linear Amidines (GO:0016813) | 3/9 | 9.68E-05 | 46.8720379 | 433.252073 | *PADI6;PADI3;PADI4* |
| Acetylcholine Receptor Regulator Activity (GO:0030548) | 3/15 | 5.00E-04 | 23.42891 | 178.089489 | *LYNX1;PSCA;SLURP1* |
| Acetylcholine Receptor Binding (GO:0033130) | 2/9 | 0.00390431 | 26.6563342 | 147.827312 | *LYNX1;PSCA* |
| Calcium-Dependent Protein Serine/Threonine Phosphatase Activity (GO:0004723) | 1/6 | 0.0625143 | 18.5737089 | 51.4930066 | *PPM1F* |
| CD4 Receptor Binding (GO:0042609) | 1/7 | 0.07254839 | 15.4773083 | 40.604742 | *HLA-DRB1* |

| | | | | | |
|---|---|---|---|---|---|
| Acetylcholine Receptor Inhibitor Activity (GO:0030550) | 1/7 | 0.07254839 | 15.4773083 | 40.604742 | *LYNX1* |
| Estradiol 17-Beta-Dehydrogenase [NAD(P)+] Activity (GO:0004303) | 1/9 | 0.092297 | 11.6068075 | 27.6560471 | *HSD17B8* |
| ATPase Activity, Coupled to Transmembrane Movement of Ions, Rotational Mechanism (GO:0044769) | 1/12 | 0.12113783 | 8.44003414 | 17.8154462 | *ATP6V0E2* |
